# MK2 Expression Promotes Non-Small Cell Lung Cancer Cell Death and Predicts Survival

**DOI:** 10.1101/2021.11.30.470656

**Authors:** Othello Del Rosario, Karthik Suresh, Medha Kallem, Gayatri Singh, Anika Shah, Yun Xin, Nicolas M. Philip, Haiyang Jiang, Franco D’Alessio, Meera Srivastava, Alakesh Bera, Larissa A. Shimoda, Michael Merchant, Madhavi J. Rane, Carolyn E. Machamer, Jason Mock, Robert Hagan, Todd M. Kolb, Mahendra Damarla

## Abstract

Non-small cell lung cancers demonstrate intrinsic resistance to cell death even in response to chemotherapy. Previous work suggested that defective nuclear translocation of active caspase 3 may play a role in resistance to cell death. Separately, our group has identified that mitogen activated protein kinase activated protein kinase 2 (MK2) is required for nuclear translocation of active caspase 3 in the execution of apoptosis. This study demonstrates a relatively low expression of MK2 in non-small cell lung carcinoma cell lines compared to small cell carcinoma cell lines. Further, overexpression of MK2 in non-small cell lung carcinoma cell lines results in increased caspase 3 activity and caspase 3 mediated cell death. Higher MK2 transcript levels were observed in patients with earlier-stage non-small cell lung cancer. Higher expression of MK2 is associated with better survival in patients with early stage non-small cell lung cancer across two independent clinical datasets. Using data sets spanning multiple cancer types, we observed improved survival with higher MK2 expression was unique to lung adenocarcinoma. Mechanistically, MK2 promotes nuclear translocation of caspase 3 leading to PARP1 cleavage and execution of cell death. While MK2 can directly phosphorylate caspase 3, neither phosphorylation status of caspase 3 nor the kinase activity of MK2 impacts caspase 3 activation, nuclear translocation and execution of cell death. Rather, a non-kinase function of MK2, specifically trafficking via its nuclear localization sequence, is required for caspase 3 mediated cell death. In summary this study highlights the importance of a non-enzymatic function of MK2 in the execution of apoptosis, which may be leveraged in the adjunctive treatment of NSCLC or other conditions where regulation of apoptosis is crucial.

## Introduction

Apoptosis is a highly regulated form of programmed cell death, a process mediated by a group of enzymes known as caspases, a family of cysteine proteases. Upon activation (i.e. cleavage), caspases act to amplify death stimuli via initiator caspases-2 and -9 (intrinsic pathway) or caspases-8 and -10 (extrinsic pathway). Once initiated, cell dismantling proceeds via the executioner caspases-3 and -7(1). Traditionally, activation of caspase 3, the major executioner caspase, has heralded cell death(1–3).

Apoptosis plays a key role in tumoral response to cytotoxic chemotherapy, which remains a mainstay of treatment for many cancers(4, 5). However, chemotherapy resistance in lung cancer is a significant problem that potentially limits effective treatment for patients with lung cancers. Since tumor death following chemotherapy administration is thought to proceed, in part, via apoptosis, the molecular basis of apoptosis resistance in tumors is a major focus of investigation(4, 6, 7). Small cell lung cancer (SCLC) is typically more sensitive to chemotherapy-induced tumor death compared to non-small cell lung cancer (NSCLC). Previous work had sought to identify differences in activation of the apoptotic cascade between NSCLC and SCLC to explain the inherent differences in response to chemotherapy(6, 7). Joseph et. al. showed that NSCLCs are indeed able to activate the apoptotic machinery in response to chemotherapy, even to the point of activating caspase 3. However, they observed that active caspase 3 was sequestered in the cytoplasm of NSCLC cell lines, unlike SCLC cell lines, where active caspase-3 translocated to the nucleus. They hypothesized that defective nuclear translocation of active caspase 3 may be a potential mechanism of chemotherapy resistance in NSCLC(7). However, the fundamental mechanism(s) by which caspase 3 translocates to the nucleus are poorly understood.

Independently, our group also recently identified a disconnect between activation of the apoptotic cascade, i.e. activation of caspase 3, and the execution of apoptosis(8). We observed that endotoxin induces caspase 3 activation, endothelial apoptosis and endothelial permeability. Notably, loss of the signaling molecule mitogen activated protein kinase activated protein kinase 2 (MK2) resulted in prevention of both apoptosis and endothelial barrier dysfunction; yet, to our surprise, caspase 3 activation was preserved. Despite being activated, when MK2 was absent, caspase-3 remained sequestered in the cytoplasm, preventing endotoxin-induced apoptosis. This cytoplasmic sequestering of caspase 3 under conditions of MK2 depletion mirror prior findings of cytoplasmic caspase 3 localization in chemo-resistant NSCLCs (9).

Thus, we hypothesized that loss of MK2 in NSCLC may be the reason why caspase 3, despite being active, fails to translocate to the nucleus to affect apoptosis. We further hypothesized that reduced basal expression of MK2 in NSCLCs mediates resistance to cell death, and high MK2 levels may be associated with improved survival in patients with non-small cell lung cancer.

### Methods and Materials

Detailed methods are in the online Supplement.

The Johns Hopkins University Institutional Animal Care and Use Committee approved all animal protocols. Male C57BL/6J (wild type, WT) mice aged 10-12 weeks (Jackson Laboratory, Bar Harbor, ME) and *MK2^-/-^* mice, C57BL/6J background (9) were exposed to intravenous (IV) PBS or lipopolysaccharide (LPS, 0127:B8, product # L3129, Sigma) via retro-orbital injection (10) for up to 6hrs.

Cell lines: Non-small cell lung carcinoma (NSCLC) cell lines- H23 and A549 cell lines were purchased from ATCC (Manassas, VA). Small cell lung carcinoma (SCLC) cell line- H446 was a kind gift from Dr. Phil Dennis. Cells were maintained in full culture media as according to ATCC’s recommendations.

Adenoviral vectors: adenoviral vectors encoding wild-type MK2 (Ad-WT MK2), constitutively active MK2 (Ad-Active-MK2; T222E, T334E)(11), dominant negative MK2 (Ad-Dom Neg-MK2; K93R)(12), a mutated nuclear export sequence MK2 (Ad-Mut-NES; L360A) (13) and a mutated nuclear localization sequence MK2 (Ad-Mut-NLS-MK2; K372A, K373A, K388A, K389A) (14, 15) were directly purchased from Vector Builder (Chicago, IL). The sequences of the plasmids encoding these vector were verified by Sanger sequencing (The Genetics Resources Core Facility, Johns Hopkins University).

Immunoblot analyses: Phospho-specific anti-total antibodies directed at HSP27(p- HSP27- CST-2401; t-HSP27- CST-2402) and anti-total antibodies directed at MK2 (CST-3042), caspase 3 (CST-9662), β-tubulin (CST-5346), GAPDH (CST-3683), PARP1 (CST-9542) (Cell Signaling, Boston, MA) were used.

Quantitative-PCR: For quantitative PCR, we isolated total RNA from cells with Trizol (ThermoFisher Scientific; Waltham, MA) and purified with a commercially available kit (QIAgen; Germantown, MD). Subsequently prepared cDNA (GE First-Strand cDNA Synthesis Kit; Niskayuna, NY) was used as the template for quantitative PCR using the BIORAD iQ5 thermal cycler. Primer sets for human Mapkapk2 (#PPH01783A) and β- Actin (#PPH00073G) were purchased from QIAgen. Background subtracted amplification data were analyzed using open-source software(16) to estimate Ct values and amplification efficiency. Target gene expression was normalized to a reference gene using the comparative Ct method(17).

Nuclear and cytosolic fractionation: Cells were trypsinized and re-suspended in PBS. Nuclear and cytosolic fractions of the resulting cell suspensions were generated using NE-PER Nuclear and Cytoplasmic Extraction Reagents (Thermo Fisher Scientific, Rockford, IL).

Brightfield microscopy: Six hours after sub-confluent plating, H23 cells were infected with Ad-eGFP or Ad-WT MK2 and imaged over time. A mark was placed under side each dish to allow for sequential imaging of the same region. All images were anonymized and number of cells per image were counted by blinded investigators (AS, GS, MK). Cell counts obtained by each investigator were subsequently averaged together.

In vitro kinase assay: Reactions were performed in 50 mM sodium β-glycerophosphate containing 10 mM Mg-acetate, 0.1 mM ATP, 2 uCi γ-^32^P-ATP (New England Nuclear, PerkinElmer, Waltham, MA) and 170 ng (6 U) of recombinant active MK2 (Millipore, Dundee, UK). Substrates were recombinant active caspase-3 (250 ng, Enzo, Farmingdale, NY) or recombinant HSP27 (14 ng, Enzo, Farmingdale, NY). Reactions were incubated at 30°C for 30 or 60 min, and resolved on 4-12% NUPAGE gels (Life Technologies, Carlsbad, CA). After staining with Coomassie Blue, gels were dried and phosphorylated proteins were detected on a Molecular Imager FX phosphorimager (BioRad, Hercules, CA) using Quantity One software.

Bio-ID assay: Adenoviral vectors encoding wild-type MK2 fused with a mutant of the *Escherichia coli* biotin protein ligase BirA (18) that is smaller and more efficient (19) on the c-terminus (Ad-WT-MK2-BioID-C) or the n-terminus (Ad-WT-MK2-BioID-N) were purchased from Vector Builder (Chicago, IL). H23 cells were infected with Ad-WT MK2, Ad-WT-MK2-BioID-C or Ad-WT-MK2-BioID-N for 24 hours after which the media was changed to include biotin for an additional 18 hours, after which cell lysates were prepared. Cell lysates were incubated with streptavidin fused Sepharose beads overnight at 4°C with gentle shaking. Beads were washed four times with buffer to elute off non-specific proteins bound to the sepharose beads. The buffer was aspirated and beads were re-suspended in Laemmli buffer for Western blotting.

Co-Immunoprecipitation: H23 cells were infected with Ad-eGFP, Ad-WT MK2, Ad-Mut- NLS-MK2 or Ad-Mut-NES-MK2 and 48 hrs afterwards cell lysates were prepared. Appropriate cell lysates (50 µg) were treated with anti-MK2 (CST-3042, 1:100 dilution) or isotype control antibody (SC-2027X, 1:100) for overnight incubation at 4°C with gentle shaking. Protein A sepharose beads (GE Healthcare 17-5280-01; 10 µl) were added on the next day and incubated with shaking for 3 hrs at 4°C. Beads were washed two times with Krebs+ buffer to elute off non-specific proteins bound to the protein A sepharose beads. The buffer was aspirated and beads were re-suspended in 20 µl 2x Laemmli sample buffer. The samples were separated by 4-12% SDS-PAGE and immunoblotted.

Flow cytometry: Following experimental exposures, cell cultures were trysinized and single cell suspension was generated. 4’,6-Diamidino-2-Phenylindole, Dihydrochloride (DAPI) was used to stain DNA of cells as a way to quantify condensed and fragmented nuclei, a hall mark of apoptosis(20). Data acquisition was performed on a custom FACSAria II instrument running FACSDiva acquisition software (BD Biosciences, San Jose, CA). A singly-stained aliquot of H23 cells for the DAPI fluorochrome and unstained cells were used to compensate for background auto-fluorescence. 5 x 10^4^ events were obtained per sample and analyzed using FCS Express 6 (De Novo Software, Pasadena, CA).

Statistical analysis: Data are shown as means (± SD). Since data is obtained using cell lines, biological replicates are not feasible. Data from separate individual cultures (each individual culture represents N of 1) are plotted for each condition. Sample size is identified in Figure Legends. The specific statistical test and post-hoc testing performed is identified within each Figure Legend. A *P* value of less than 0.05 was considered significant. Data were analyzed using GraphPad Prism 8 (La Jolla, CA).

### TCGA Data access and analysis

#### Clinical and mRNA expression accession

Clinical data for TCGA datasets was accessed using the *TCGABiolinks* package (21–23) in R/Bioconductor (24, 25). mRNA expression data was obtained using OncoLnc (26). Patient level clinical and mRNA data were merged using individual TCGA sample identifiers. The code used to analyze the TCGA data is presented in markdown format in **Supplemental Data**. All raw data and code pertaining to TCGA analyses performed herein are available at: github.com/suresh-lab/TCGA-casp3/

#### Data Filtering/Processing

After downloading the LUAD dataset, we parsed entries based on sample type (normal tissue and primary tumors). For tumor samples, we restricted our analyses to tumors identified on histology as adenocarcinoma-type, and to patients who had either a date of death or a date of last follow-up. Since mRNA transcript levels can vary based on processing, filtering and normalization parameters, we wanted to verify the accuracy of mRNA transcript counts obtained from OncoLnc web interface. So, these values were compared to normalized RSEM transcript levels for the MAPKAPK2 gene that were downloaded and extracted from the TCGA-LUAD dataset using the *TCGAbiolinks* package (**Supplemental Figure 1**). Since the values were exactly correlated, we chose to continue using transcript levels (for MAPKAPK2 as well as other genes) obtained from OncoLnc given ease of access. For our validation cohort, MAPKAPK2 mRNA transcript levels (log mRNA expression Seq V2 RSEM) and clinical data for the validation cohort was obtained via cBioportal(27, 28).

#### Choice of Models

We explored the association between MK2 transcript levels and survival using the following models:

##### Model 1

Cox Proportional Hazards model of time to death in one year (right censored at one year of follow-up), stratified by tumors stage, adjusted for sex and smoking status, using MK2 as a dichotomous variable (high / low).

##### Model 2

Logistic regression for risk at one year, stratified by tumor stage (early vs. late stage), and adjusted for sex and smoking status, using MK2 as a dichotomous variable (high / low).

In the TCGA-LUAD dataset, we observed that MK2 levels were significantly lower in late-stage tumor samples. Thus, we opted to stratify our analyses by tumor stage (early vs. late stage). Such a baseline imbalance was not noted in our validation cohort; thus, in this cohort, we chose to include stage as a covariate in our multivariate analysis. In both TCGA and validation datasets, we initially modeled MK2 transcript levels as a continuous variable and noted an association between MK2 transcript levels and survival at one year (**Supplemental Data**). However, we reasoned that examining MK2 transcript levels as “high” vs “low” (defined as top 1/3 vs. bottom 2/3 of transcript levels) may have more translational relevance in terms of risk-stratifying early stage NSCLC patients. Thus, for TCGA-LUAD, models were run with a) stratification by tumor stage and b) dichotomization of MK2 as “high” and “low”.

##### Censoring

We initially right censored the TCGA-LUAD dataset at one year because we were interested in early survival effects of changes in tumor MK2 levels. However, in the validation dataset, there were too few events (deaths) to perform a meaningful survival analysis censored at one year. Thus, we opted to look additionally at 2-year mortality so that the same analysis could be performed in both cohorts.

##### Model construction and testing

Cox proportional hazards models were constructed using the *survminer* (29) and *survival* packages in R (30). Proportional hazard assumption testing was performed by examining Schoenfield residuals and Q-Q plots. During initial exploration of the data, we observed that patients were entered into the TCGA dataset over a long period of time (1991–2013). The number of patients enrolled (per year) was skewed, with the majority of patients being enrolled after 2005 (**Supplemental Figure 2A**). To determine whether time of enrollment could be confounding our results, we performed a jackknife analysis whereby we examined the effect of sequential exclusion of one particular year of data (i.e. excluding all patients diagnosed in that year) on the point estimate results for Model 1 (**Supplemental Figure 2B**). We observed that the MK2 hazard ratio and confidence intervals were very stable even with specific years were excluded, suggesting that imbalances in enrollment time were unlikely to play a substantial role in our results. Some survival models (e.g. modeling survival across the entire span of survival time in the validation cohort) failed to meet proportionality hazard assumption testing. Thus, these models were not used.

##### Model results

For survival analysis, Kaplan-Meier curves and Hazard ratios were calculated. For logistic regression, odds ratios as well as predicted probabilities were calculated. Point estimates are provided with 95% confidence intervals. A *P* value < 0.05 was accepted as statistically significant.

## Results

### Non-Small Cell Lung Cancers have Lower MK2 Expression

To first test the hypothesis that NSCLCs have lower MK2 expression, we obtained previously characterized NSCLC and SCLC cell lines(7). As can be seen in **Figure 1A**, there is markedly reduced MK2 protein expression in the H23 NSCLC cell line compared to the H446 SCLC cell line. In order to ensure that this was not an isolated finding, a different NSCLC cell line, A549, was also tested. As can be seen in **Figures 1B-C** and **Supplemental Figure 3** there is marked reduction in MK2 protein expression in both H23 and A549 cells as compared to the H446 cells. Furthermore, lower protein expression in H23 and A549 cells was accompanied by lower MK2 mRNA expression, **Figure 1C**. Next, we interrogated the database of the Broad institute’s dependency map Consortium (www.depmap.org), which provides proteomic data across a variety of cancer cell lines. There we identified proteomic data for MK2 expression from 64 NSCLC cell lines and compared them to data from 13 SCLC cell lines(31). As can be seen in **Figure 1D**, MK2 protein expression in NSCLC cell lines was decreased compared to SCLC cell lines.

**Figure 1:**
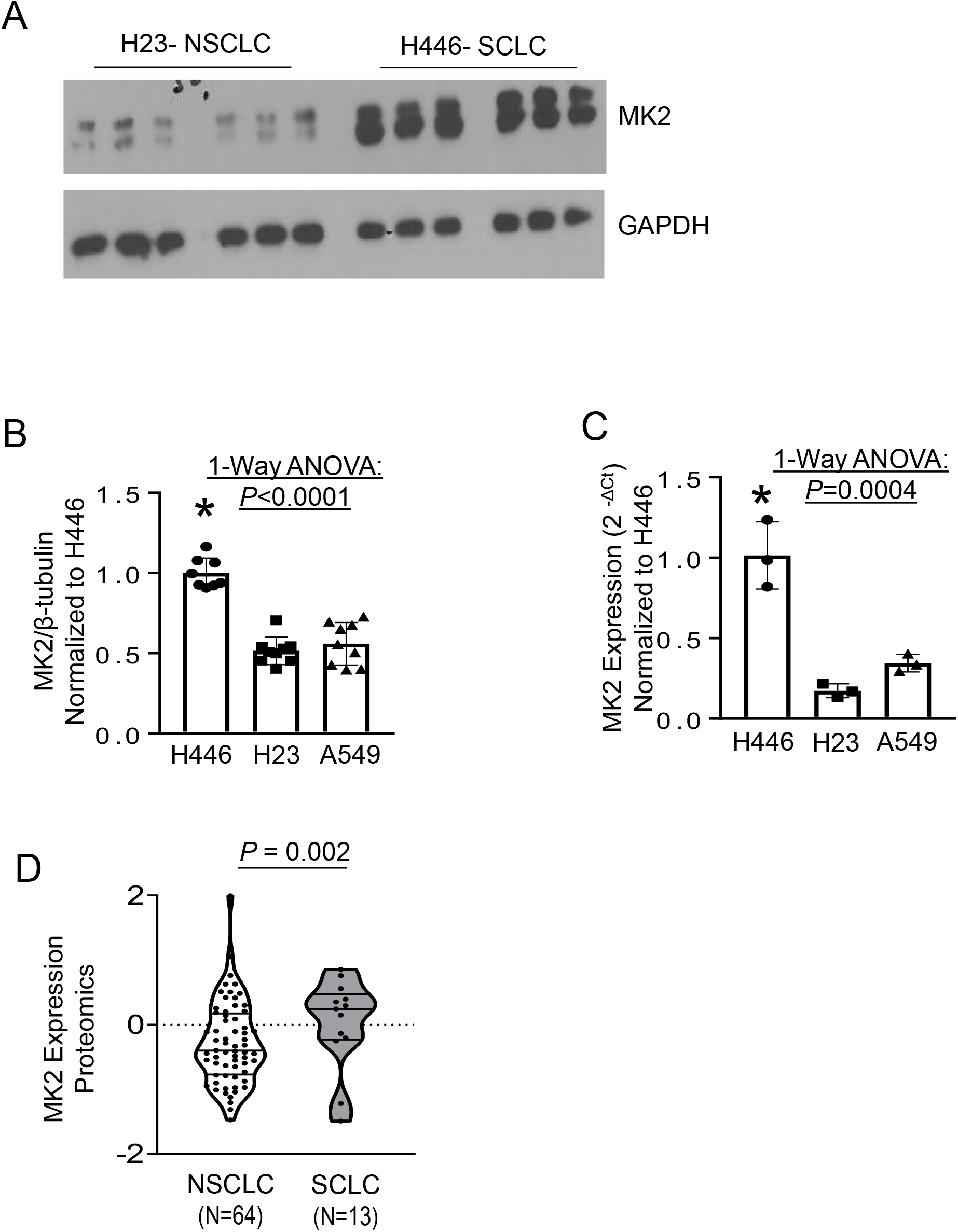
Reduced expression of MK2 in non-small cell lung carcinoma cell lines. **A.** Cell lysates from the NSCLC cell line, H23, and the SCLC cell line, H446 were immunoblotted with antibodies recognizing total MK2. As shown, there is markedly reduced MK2 expression in H23 cells compared to H446 cells. **B.** Quantification of immunoblot analyses of MK2 expression shows increased MK2 expression in SCLC cell line, H446, compared to the NSCLC cell lines, H23 and A549. *, *P* < 0.05 vs all other conditions using post-hoc Dunnett’s multiple comparison test. N= 9 per group. **C.** Real- time PCR analysis demonstrates significantly reduced MK2 gene expression in the NSCLC cell lines, H23 and A549 as compared to the SCLC cell line, H446. *, *P* < 0.05 vs all other conditions using post-hoc Dunnett’s multiple comparison test. N= 3 per group. **D.** Relative protein expression of MK2 using proteomics of lung cancer cell lines examined by the Cancer Dependency Map project (www.depmap.org) shows significantly lower MK2 expression in NSCLC cell lines compared to SCLC cell lines using Wilcoxon Rank Sum Test.

### Expression of MK2 Induces Cell Death in H23 NSCLC cells

We reasoned that if the loss of MK2 confers resistance to cell death in NSCLC, then forced expression of MK2 should sensitize these cells to apoptosis. To test this hypothesis, we overexpressed MK2 in NSCLC cells. Using the H23 cells as a model cell type (given endogenously low amount of MK2), we first verified that we could significantly express MK2 protein using adenoviral gene delivery in H23 cells, **Supplemental Figure 4A**. We then developed a work flow for using flow cytometry to measure and quantify cell death by DAPI staining(20), **Supplemental Figure 4B.** Using these approaches, we observed a dose-dependent increase in cell death at 48 hrs based on Ad-WT MK2 multiplicity of infection, **Figure 2A**. To determine the longer-term effects of MK2 expression, H23 cells were infected with Ad-eGFP or Ad-WT MK2 and imaged over time. As shown in **Figure 2B-C**, H23 cells infected with Ad-eGFP continue to grow over time, whereas those infected with Ad-WT MK2 failed to grow and in fact have significantly reduced cell numbers at 144 hrs following plating. These data show that in the H23 NSCLC cell line, expression of MK2 leads to increased cell death.

**Figure 2:**
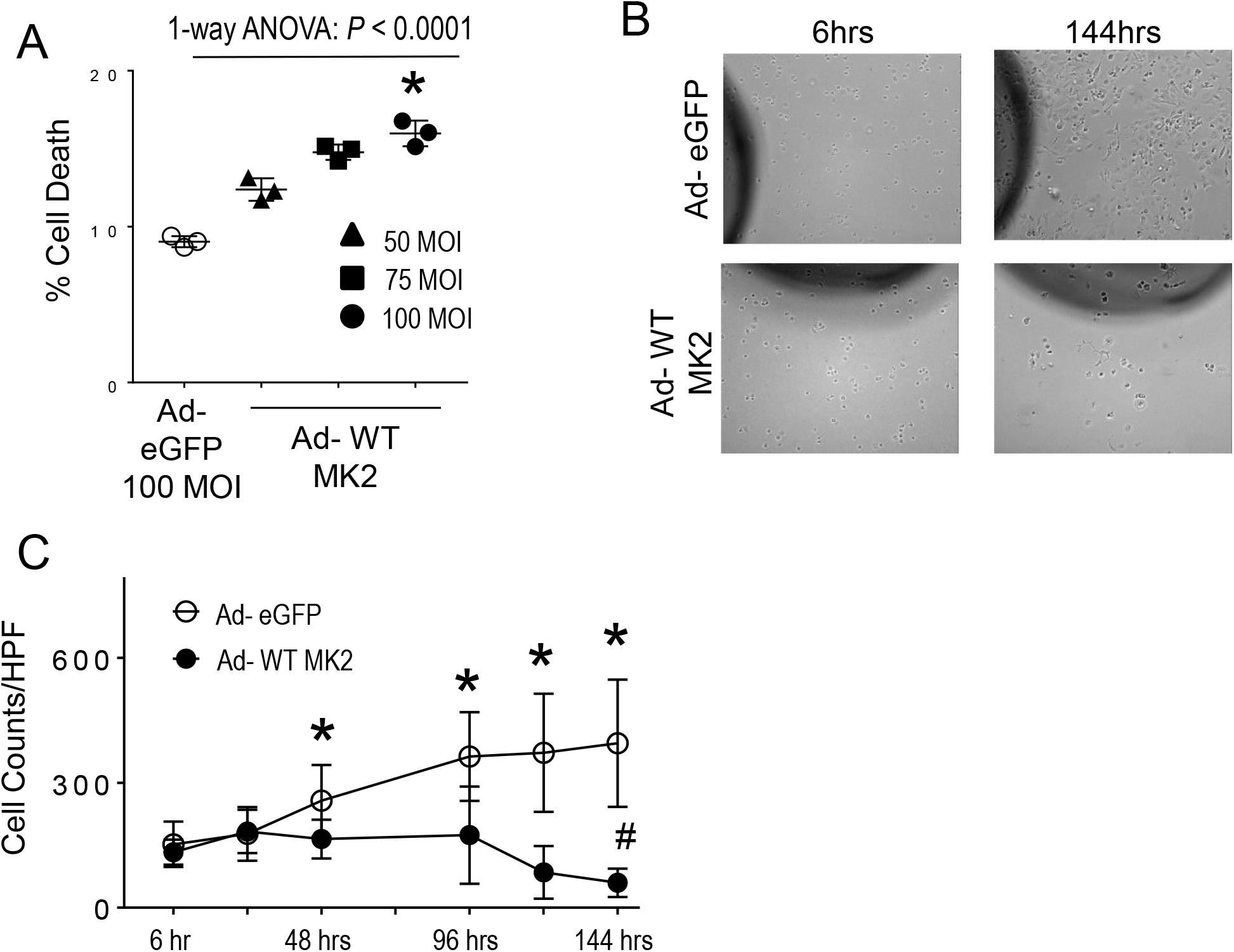
MK2 expression leads to caspase 3 mediated apoptosis in H23 NSCLC cells. H23 NSCLC cells were infected with adenovirus encoding wild type MK2 or eGFP (Ad-WT MK2 and Ad-eGFP, respectively) for up to 144hrs for analyses. **A.** After 48 hours of infection, there is a dose-dependent increase in cell death based on Ad-WT MK2 multiplicity of infection. *, *P* < 0.05 vs Ad-eGFP and 50 MOI using Dunnett’s multiple comparison test. **B.** Time lapse bright-field microscopy was used to quantify the effect of MK2 expression on H23 cells. Six hours after sub-confluent plating, H23 cells were infected with Ad-eGFP or Ad-WT MK2 and imaged over time, up to 144hrs. **C.** Quantification of cell counts demonstrates Ad-eGFP infected cells continue to grow over time compared Ad-WT MK2 infected cells. H23 cells infected with Ad-WT MK2 resulted in reduced cell counts compared to baseline. N=22 individual cultures per group were analyzed over time. *, *P* < 0.05 vs Ad-WT MK2 by un-paired Mann Whitney test. #, *P* < 0.05 vs Ad-WT MK2 6hr by un-paired Mann Whitney test.

### Expression of MK2 Induces Cell Death in A549 NSCLC cells

Having shown expression of MK2 induces cell death in H23 NSCLC cells, we sought to determine if this process could be relevant in other NSCLC cells. As shown in **Figure 1**, MK2 protein expression was low at baseline in both A549 and H23 NSCLC cells. When we infected A549 cells with Ad-WT-MK2 (**Figure 3A**), we observed significant increase in caspase 3 activity and cell death as compared to Ad-eGFP, **Figure 3B-C**. The intrinsically low level of apoptosis in A549 cells as compared to H23 cells has been noted previously(32).

**Figure 3:**
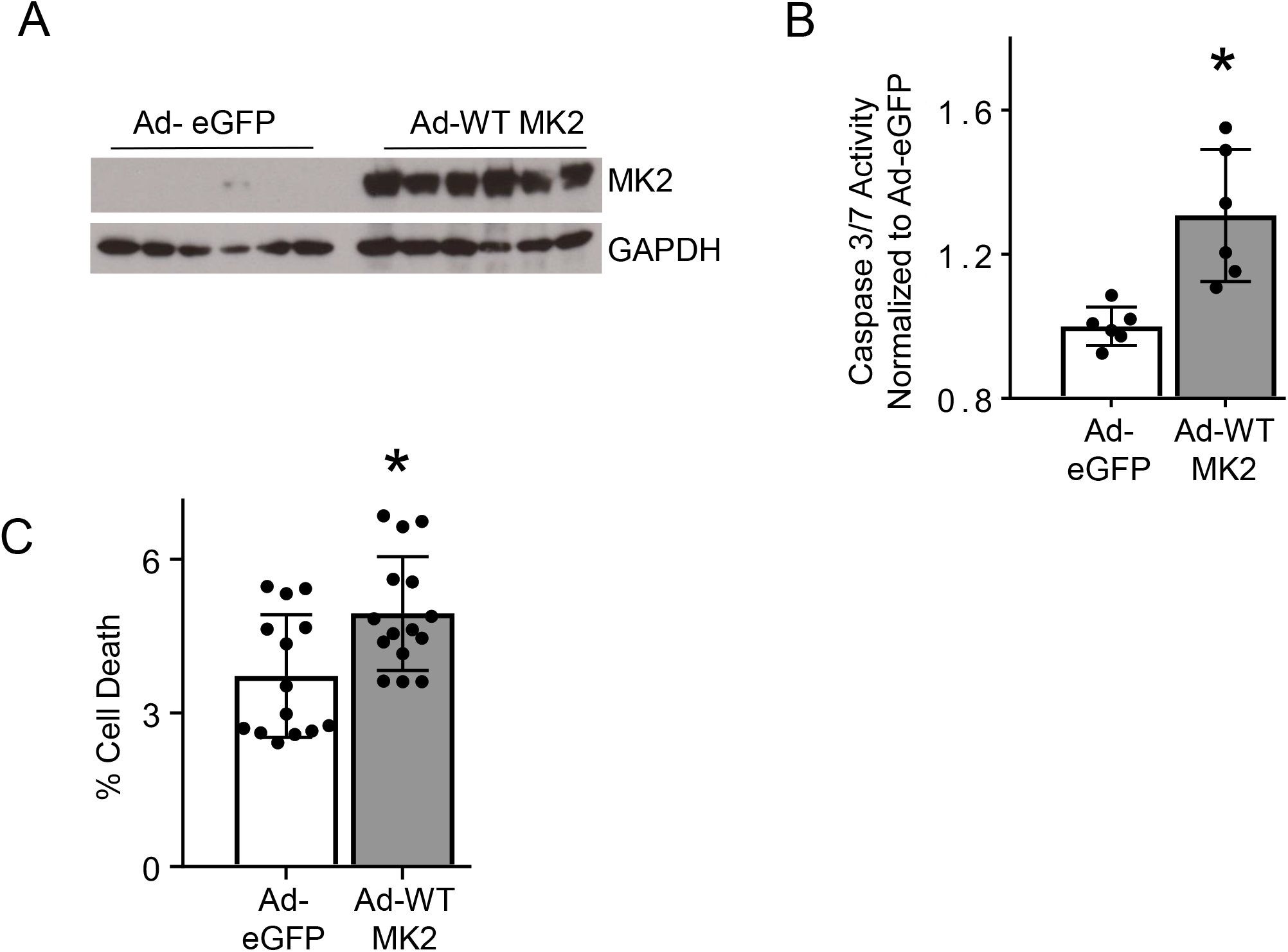
MK2 expression leads to cell death in A549 NSCLC cells. **A.** A549 cells infected with Ad-WT MK2 leads to marked increase in MK2 protein expression. **B.** A549 NSCLC cells were infected with Ad-eGFP or Ad-WT MK2 and 48hrs afterwards lysates were assessed for caspase 3/7 activity. Ad-WT MK2 infected A549 cells exhibited significantly increased caspase 3/7 activity compared to cells infected with Ad-eGFP. *, *P*< 0.05 using unpaired t test. N=6 per group **C.** A549 NSCLC cells were infected Ad- eGFP or Ad-WT MK2 and 48hrs afterwards cells were assessed for cell death using flow cytometry. Ad-WT MK2 infected A549 cells exhibited significantly increased cell death compared to cells infected with Ad-eGFP. *, *P*< 0.05 using unpaired t-test. N=15 per group.

### Low Tumor MK2 Expression is Associated with Late Stage Lung Adenocarcinoma

Given our data showing that MK2 over-expression leads to increased cell death in NSCLC cell lines, we sought to test if MK2 expression was associated with clinical outcomes in patients with NSCLC. Because many publicly available databases only have information for mRNA transcript, we first sought to determine the relationship between MK2 mRNA transcript to protein expression, to ascertain whether mRNA transcript level was a reasonable surrogate for protein expression. As shown in **Supplemental Figure 5**, we observed a direct, statistically significant correlation between MK2 mRNA and MK2 protein expression across NSCLC cell lines. With these data in hand, we turned to using the TCGA-LUAD dataset (n=483 patients), which contains tumor (and adjacent normal tissue) RNA-seq data for a large number of lung cancer patients. We observed a significantly lower level of MK2 transcript in late-stage (i.e. Stage III and IV) lung adenocarcinoma samples compared to early stage (i.e. Stage I and II), **Figure 4A**. This difference in MK2 transcript levels was not seen in adjacent normal tissue samples (n=56), **Figure 4B**.

**Figure 4:**
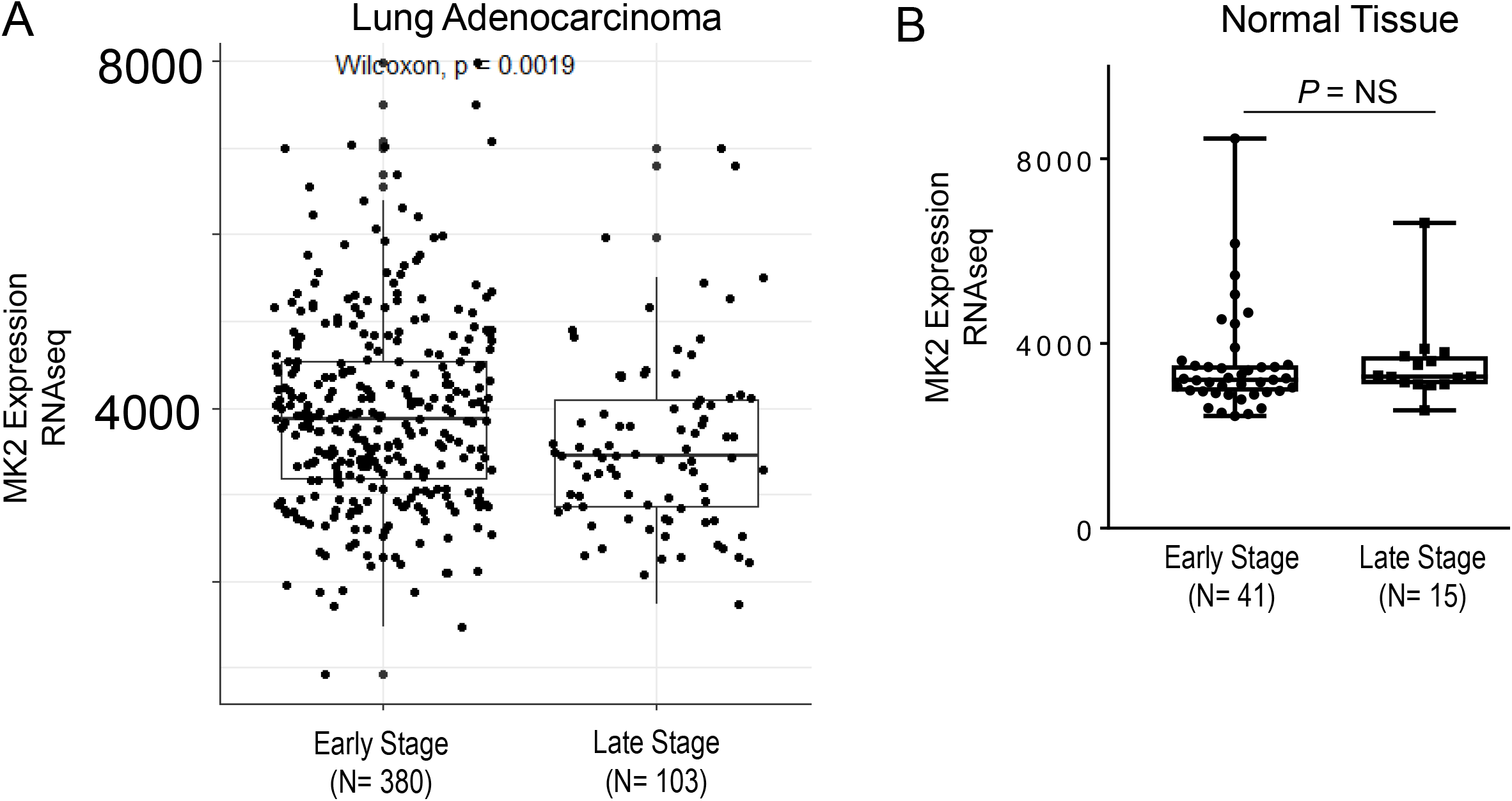
MK2 expression is lower in tumors of patients with late stage lung adenocarcinoma. **A.** Using The Cancer Genome Atlas Lung Adenocarcinoma (TCGA- LUAD) database, there is significantly lower MK2 expression in tumors of patients diagnosed with late stage (stage III and stage IV) as compared to those diagnosed with early stage (stage I and stage II). **B.** In a subset of patients from the TCGA-LUAD database, adjacent normal lung tissue was also sampled. There is no difference in MK2 expression in normal lung tissue between patients diagnosed with late stage and early stage lung adenocarcinoma using Wilcoxon Rank Sum Test.

### High Tumor MK2 Expression is Associated with Survival in Early Stage Lung Adenocarcinoma

Next, we modeled the effect of MK2 transcript level on overall survival in the TCGA-LUAD cohort using univariate (**Figure 5**) and multivariate (**Table 1**) Cox proportional hazards and multivariate logistic regression (**Table 1**). In both models, the association between high MK2 transcript level and death was stratified by tumor stage (early/late), and adjusted for age, sex and smoking status. High MK2 transcript level (defined as transcripts in the top tertile) was associated with improved survival at one year (HR 0.24, 95%CI: 0.07-0.80; OR: 0.22, 95% CI: 0.06-0.77) and two years (HR 0.52, 95%CI: 0.28-0.98; OR: 0.49, 95%CI: 0.25,0.98) in early stage, but not late stage, NSCLC. Similarly, the proportion of patients who died at one year, stratified by cancer stage, correlated more strongly with MK2 transcript tertile in early (R^2^ = 0.84) than late (R^2^ = 0.4) stage NSCLC (**Supplementary Figure 6**). To validate these results, we examined the association between MK2 transcript levels and survival in an independent cohort of 305 Asian patients with lung adenocarcinoma(33). As discussed in Methods, due to a small number of deaths occurring in the first year, we only performed 2-year analyses with this dataset. In multivariate analysis, we again observed improved survival associated with higher MK2 transcript levels (HR: 0.1, 95%CI: 0.01-0.81; OR: 0.08, 95%CI: 0.004, 0.154) after adjustment for age, sex, smoking and cancer stage (**Figure 5C**, **Table 2**). Interestingly, unlike the TCGA cohort, smoking and male sex were also significantly associated with mortality in the validation cohort; however, the association between MK2 and mortality remained significant even after adjustment for these covariates. Additionally, sensitivity analyses showed that adjustment for caspase 3 transcript levels did not change the results (**Supplemental Data**).

**Figure 5.**
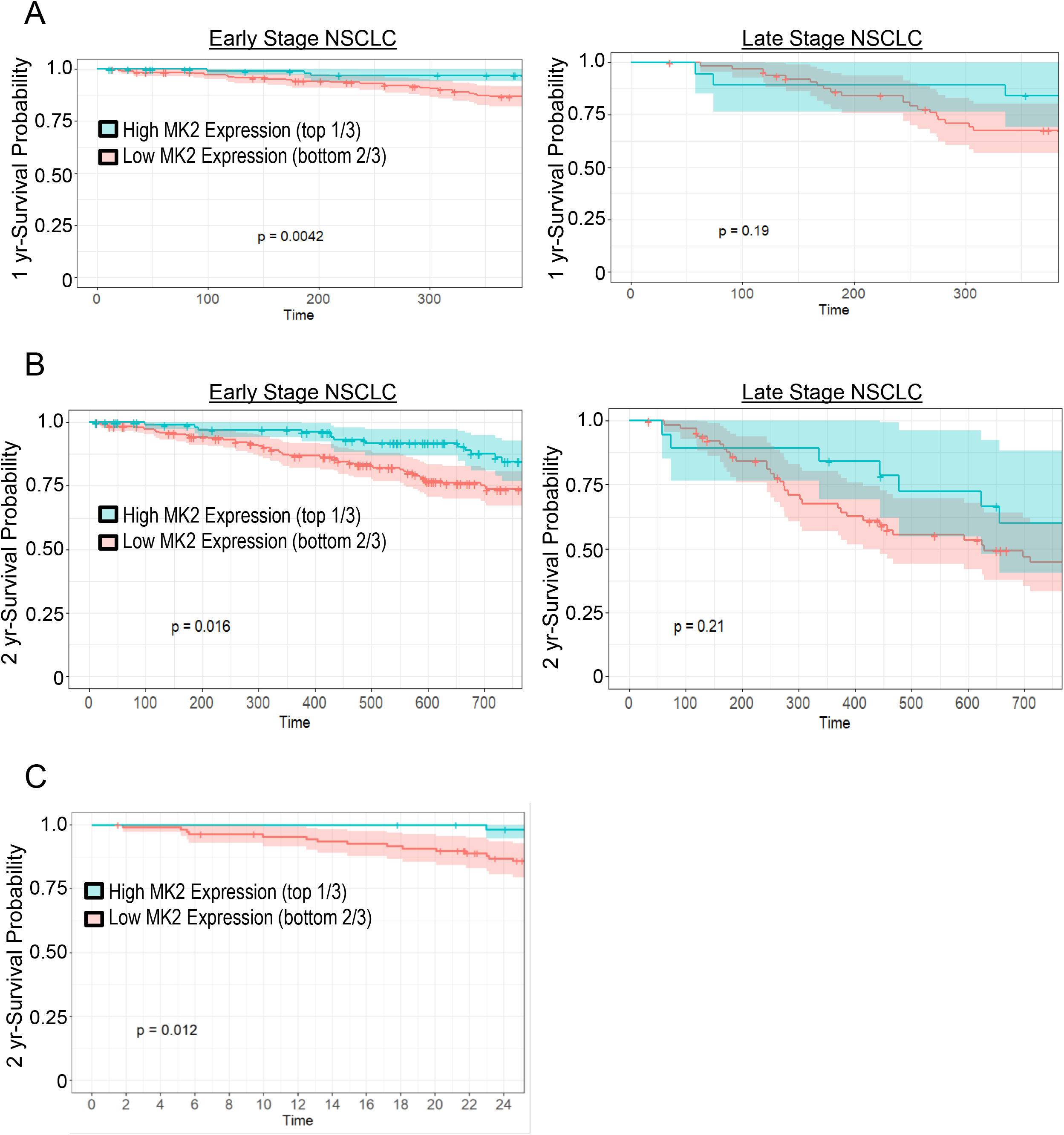
Higher MK2 Expression is associated with improved survival in patients with NSCLC. Kaplan-Meier curves showing 1-year (**A**) and 2-year (**B**) survival in patients with early stage (left column, N= 312) and late stage (right column, N= 86) NSCLC in the TCGA-LUAD dataset, stratified by high (teal) and low (red) MK2 transcript levels. Time is shown in days. **C.** Kaplan-Meier curve showing 2 year survival in patients with lung adenocarcinoma in the validation dataset (N=169), stratified by high (teal) and low (red) MK2 transcript levels. Time is shown in months.

**Table 1:** MK2 expression and death at 1 and 2 years in TCGA-LUAD (N=398)

| Endpoint | 1 year mortality |  |  |  | 2 year mortality |  |  |  |
| --- | --- | --- | --- | --- | --- | --- | --- | --- |
| Tumor Stage | Early Stage |  | Late Stage |  | Early Stage |  | Late Stage |  |
| Model Type | **Model 1<sup>a</sup>**<br>HR (95% CI) | **Model 2<sup>b</sup>**<br>OR (95% CI) | **Model 1**<br>HR (95% CI) | **Model 2**<br>OR (95% CI) | **Model 1<sup>a</sup>**<br>HR (95% CI) | **Model 2<sup>b</sup>**<br>OR (95% CI) | **Model 1**<br>HR (95% CI) | **Model 2**<br>OR (95% CI) |
| High MK2 expression<sup>c</sup> | **0.24 (0.07-0.80)** | **0.22 (0.06,0.77)** | 0.48<br>(0.14,1.66) | 0.48<br>(0.12,1.92) | **0.52 (0.28,0.98)** | **0.49 (0.25,0.98)** | 0.61<br>(0.27,1.40) | 0.70<br>(0.23,2.03) |
| Male Sex | 2.2 (0.99-5.11) | 2.20 (0.92, 5.22) | 1.21<br>(0.53,2.76) | 1.40<br>(0.51,3.78) | 1.52<br>(0.89,2.60) | 1.47<br>(0.81,2.70) | 1.0 (0.53,1.89) | 0.97<br>(0.40,2.34) |
| Smoking | 1.06<br>(0.42,2.69) | 1.18<br>(0.44,3.14) | 0.81<br>(0.35,1.90) | 0.68<br>(0.24,1.90) | 1.36<br>(0.71,2.60) | 1.45<br>(0.71,2.95) | 0.90<br>(0.46,1.75) | 0.77<br>(0.30,1.95) |
| Age | 0.98 (0.94, 1.02) | 0.98<br>(0.94,1.02) | 1.02<br>(0.98,1.07) | 1.02<br>(0.97,1.08) | 1.0<br>(0.97,1.03) | 1.0<br>(0.97,1.03) | 1.01<br>(0.98,1.05) | 1.01<br>(0.97,1.06) |

<sup>a</sup> Model 1: Cox Proportional hazards modeling of time to death, right censored at 2 years, as a function of high MK2 expression (dichotomous: top 1/3 vs bottom 2/3 of transcript expression), adjusted for age, sex and smoking status
<sup>b</sup> Model 2: Logistic regression modeling death at 2 years as a function of high MK2 expression, adjusted for age, sex, and smoking status
<sup>c</sup> Top 1/3 of MK2 transcript levels (reference: bottom 2/3)

**Table 2:** MK2 expression and 2-year mortality in validation cohort (n=169)

| Model type | Early Stage |  |
| --- | --- | --- |
| Variable | **Model 1<sup>a</sup>**<br>HR (95% CI) | **Model 2<sup>b</sup>**<br>OR (95% CI) |
| High MK2 expression<sup>c</sup> | **0.10 (0.01,0.81)** | **0.08 (0.004,0.154)** |
| Male Sex | 1.29 (0.40,4.20) | 1.32 (0.33,5.28) |
| Smoking | **3.44 (1.01, 11.74)** | **4.38 (1.15,18.89)** |
| Age | 1.02 (0.98,1.08) | 1.04 (0.98,1.10) |
| Late Stage<sup>d</sup> | **3.81 (1.42,10.21)** | **4.99 (1.54,16.83)** |

<sup>a</sup> Model 1: Cox Proportional hazards modeling of time to death, right censored at 2 years, as a function of high MK2 expression (dichotomous: top 1/3 vs bottom 2/3 of transcript expression), adjusted for age, sex smoking status and tumor stage
<sup>b</sup> Model 2: Logistic regression modeling death at 2 years as a function of high MK2 expression, adjusted for age, sex, smoking status and tumor stage
<sup>c</sup> Top 1/3 of MK2 transcript levels (reference: bottom 2/3)
<sup>d</sup> Early Stage: Stage I/II; Late Stage: Stage III/IV

### MK2 Expression Leads to Caspase 3 Mediated Apoptosis in H23 NSCLC cells

Having observed survival benefits with increased MK2 levels across multiple lung cancer datasets, we explored whether the mechanism by which MK2 expression may be affecting cell death is caspase 3 dependent, i.e. apoptosis. After infection of H23 cells with Ad-WT MK2, there is marked cleavage of caspase 3, the active form, as compared to Ad-eGFP, **Figure 6A**. We have previously noted that MK2 expression was not required for caspase 3 activation in mouse lung tissues or human microvascular lung endothelial cells(8). To determine whether MK2 is required for caspase 3 activation in NSCLC, we first performed a dose titration of etoposide, a commonly used chemotherapeutic agent, in H23 cells to find a dose that would induce cell death. As can be seen in **Supplemental Figure 7**, with 96µg/ml dosing of etoposide we observed statistically significant increase in cell death. We then measured caspase 3 activity in

**Figure 6:**
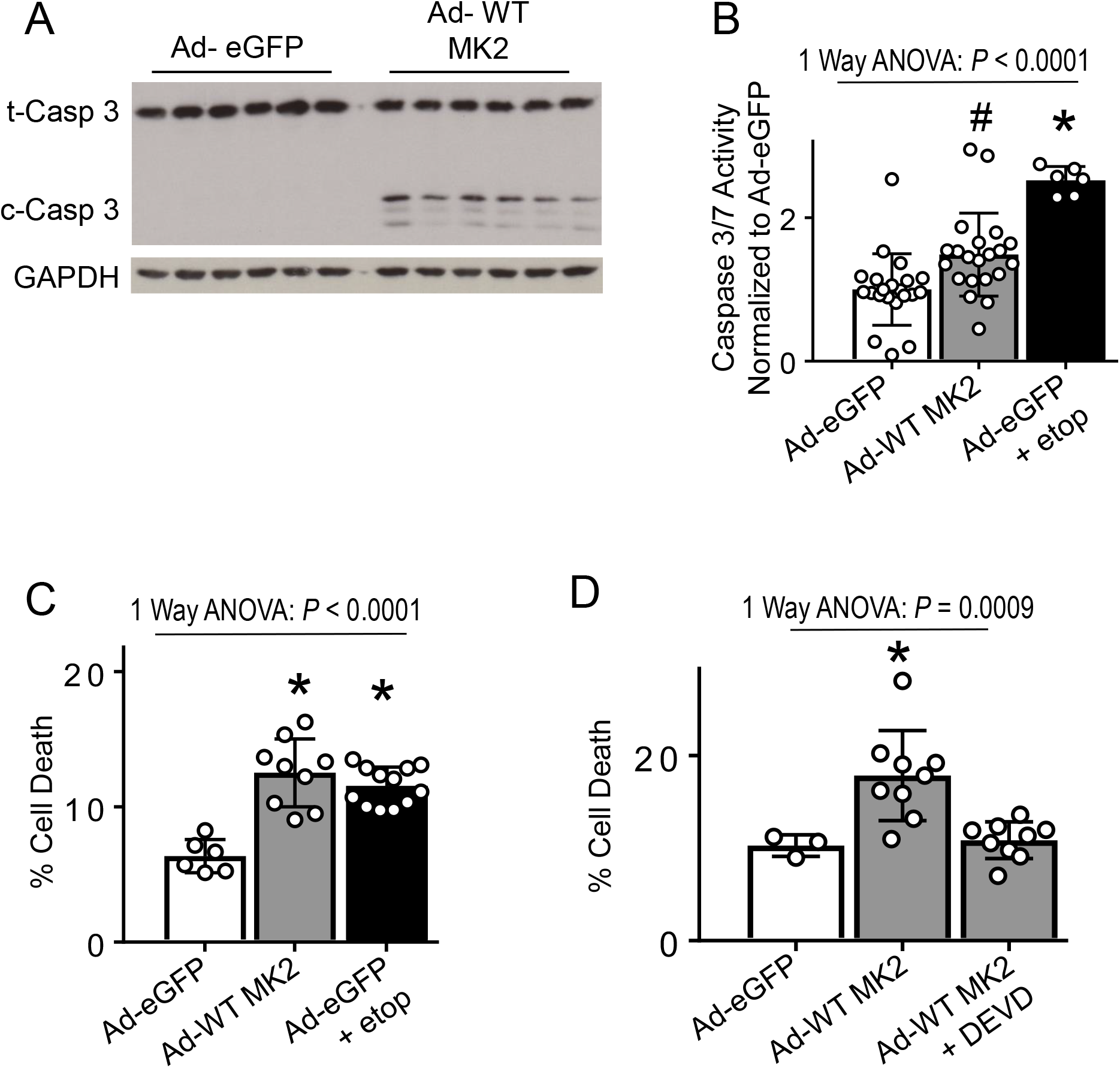
MK2 expression leads to caspase 3 mediated apoptosis in H23 NSCLC cells. **A.** Cell lysates from H23 cells infected with Ad-eGFP or Ad-WT MK2 for 48 hours were analyzed for protein expression. As shown, there is significant increase in cleaved caspase 3 in lysates from cells infected with Ad-WT MK2. t-Casp 3, total caspase 3; c- Casp 3, cleaved caspase 3. **B.** H23 cells were infected with Ad-eGFP for 24 hours and then treated with vehicle or etoposide 96ug/mL for 24hrs or Ad-WT MK2. Cell lysates were analyzed for caspase 3 activity. As shown, there is significant increase in caspase 3/7 activity with MK2 expression. Ad-eGFP infected H23 cells treated with etoposide exhibited the greatest increase in caspase 3/7 activity. #, *P* < 0.05 vs Ad-eGFP. *, *P* < 0.05 vs all others. All post-hoc analyses using Tukey’s multiple comparisons test. N= 6- 21 per group. **C.** H23 cells were infected with Ad-eGFP for 24 hours and then treated with vehicle or etoposide 96ug/mL for 24hrs or Ad-WT MK2. Cells were then analyzed for cell death using flow cytometry. As shown, there is significant increase in cell death with Ad-WT MK2. There is no difference in cell death between Ad-WT MK2 infected cells and Ad-eGFP infected cells treated with etoposide. *, *P* < 0.05 vs Ad-eGFP. All post-hoc analyses using Tukey’s multiple comparisons test. N=6-12 per group. **D.** H23 cells were infected with Ad-WT MK2 for 24 hours and then pre-treated with vehicle or DEVD (caspase 3 inhibitor, 50ug/mL, 1hr) and then exposed to etoposide 96ug/mL for 24hrs. Cells were then analyzed for cell death using flow cytometry. Pre-treatment with DEVD abrogated the MK2 expression-induced cell death. *, *P* < 0.05 vs all others. All post-hoc analyses using Tukey’s multiple comparisons test. N= 3-9 per group.

Ad-WT MK2 or Ad-eGFP infected H23 cells treated with vehicle or 96µg/ml of etoposide. Ad-WT MK2 infected H23 cells showed a significant increase in caspase 3 activity. However, Ad-eGFP infected cells exposed to etoposide demonstrated even higher caspase 3 activity than with Ad-WT MK2 infection alone, **Figure 6B**, demonstrating MK2 expression is not required for caspase 3 activation. Despite having a significantly lower amount of caspase 3 activity, Ad-WT MK2 infected cells exposed to vehicle had a similar amount of cell death when compared to Ad-eGFP infected H23 cells exposed to etoposide, **Figure 6C**. Given the disproportionate effect of MK2 expression on cell death (in relation to caspase 3 activity), we sought to determine if apoptosis, i.e. caspase 3-mediated cell death, was responsible for the cell death observed. Therefore, after infection of H23 cells with Ad-WT MK2, cells were either pre-treated with DEVD, a caspase 3 specific inhibitor, or its diluent, DMSO. As shown in **Figure 6D**, pretreatment with DEVD completely abrogated MK2 expression-induced cell death. These data indicate that MK2 over-expression was sufficient to induce and potentiate caspase 3-mediated apoptosis in H23 NSCLC cells.

### MK2 Directly Phosphorylates Caspase 3 but does not Affect its Activity

We next explored potential mechanistic interactions between MK2 and caspase 3 that might explain the synergy between MK2 and caspase 3 with regards to apoptosis in NSCLC. Because MK2 is a kinase, we first sought to determine if MK2 could directly phosphorylate caspase 3. In wild type mice intravenous LPS exposure, activates MK2 and induces caspase 3-mediated cell death(8). In lung homogenates obtained from LPS exposed mice, 2-Dimensional immunoblotting for caspase 3 demonstrated a shift toward the positive electrode on an isoelectric gradient, **Supplemental Figure 8**. Treatment of lung homogenates of WT mice exposed to IV LPS with active recombinant serine/threonine phosphatase, PP2A, reversed these charge-based shifts, suggestive of a LPS-induced phosphorylation event on caspase 3. Additionally, 2-Dimensional immunoblotting for caspase 3 of lung homogenates from *MK^-/-^* mice exposed to LPS is similar to that of lung homogenates from WT animals exposed to LPS and treated with active recombinant PP2A, suggestive of a MK2-dependent phosphorylation event on caspase 3, as MK2 is a serine/threonine kinase, **Supplemental Figure 8**. Therefore, we tested the hypothesis that synergistic interaction between MK2 and caspase 3 on cell death is mediated by direct phosphorylation of caspase 3. Using recombinant proteins in cell free assays, we observed that activated MK2 is able to directly phosphorylate caspase 3, **Figure 7A**. We next tested if MK2 could phosphorylate caspase 3 *in vitro* using H23 cells. 2-Dimensional gel immunoblotting of caspase 3 in Ad-WT MK2 infected H23 cells, demonstrated a leftward shift in the isoelectric focus towards the positive electrode, **Figure 7B**, compared to Ad-eGFP infected cells. Additionally, treatment of Ad-WT MK2 cell lysates with PP2A normalized the leftward shift of caspase 3, **Figure 7B**. To further isolate the effects of MK2’s kinase activity on caspase 3, we generated functional mutations of MK2 that alter its enzymatic activity: a constitutively active mutant of MK2 (Ad-Active-MK2) and a dominant negative mutant of MK2 (Ad-Dom-Neg- MK2), as previously described(11, 12). As shown in **Figure 7C**, lysates of H23 cells infected with Ad-Active-MK2 demonstrated a significant phosphorylation of HSP27, while lysates of H23 cells infected with Ad-Dom-Neg-MK2 demonstrate an inability to phosphorylate HSP27, **Figure 7C**. Utilizing these kinase activity mutant constructs of MK2, we performed two-dimensional immunoblotting for caspase 3. As seen in **Figure 7D**, there is a leftward shift toward the positive electrode on an isoelectric gradient in Ad-Active-MK2 infected H23 cells that is not present in Ad-Dom Neg-MK2 infected cells, suggestive of a MK2-dependent phosphorylation event on caspase 3. Given the evidence for phosphorylation of caspase 3, we next sought to determine if the phosphorylation status of caspase 3 altered its enzymatic activity. As shown in **Figure 7E**, treatment of lysates of H23 cells infected with Ad-eGFP or Ad-WT MK2 with PP2A did not change caspase 3 activity. Furthermore, infection of H23 cells with either Ad- Active-MK2 or Ad-Dom-Neg-Mk2, does not differentially affect caspase 3 activity, **Figure 7F**, despite having a differential effect on caspase 3 phosphorylation (**Figure 7D**). In sum, these data suggest that while MK2 phosphorylates caspase 3, this function does not affect caspase 3 activity.

**Figure 7:**
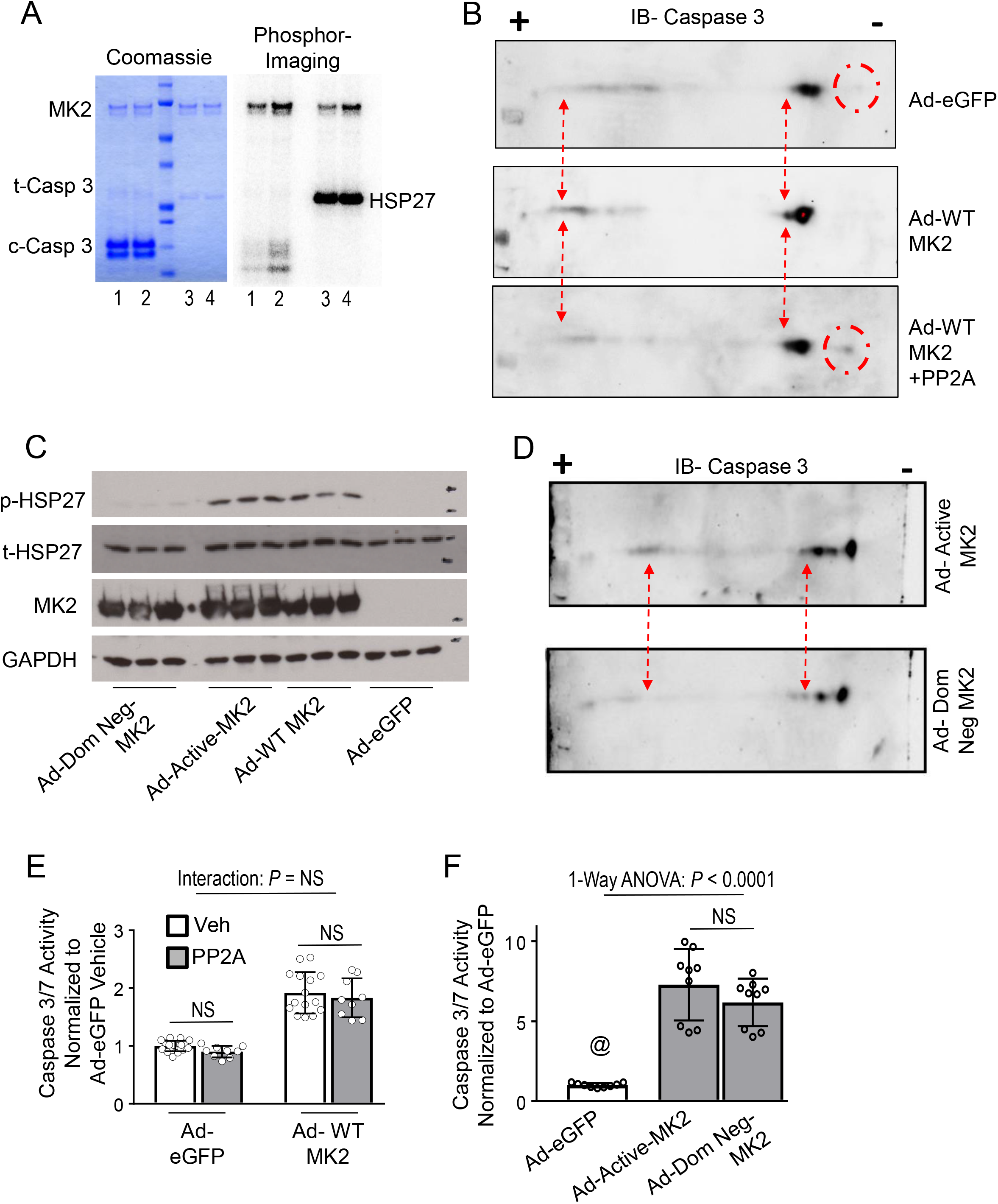
MK2 dependent phosphorylation of caspase 3 does not affect caspase 3 activity. **A.** Kinase activity assays of recombinant activated MK2 (1μM) and recombinant caspase 3 (100μM) with ^32^P-ATP. Coomassie stained gel shows relative protein amounts. Phosphorimaging shows phosphorylation of caspase 3. Lanes: 1- 30min reaction; 2- 60min reaction; 3, 4- 100nM of HSP27, 30min reaction as positive control. **B.** H23 cells were infected with Ad-eGFP or Ad-WT-MK2 (wild-type MK2) and cell lysates underwent 2-Dimensional immunoblotting for caspase 3. As shown, there is a shift toward the positive electrode on an isoelectric gradient in Ad-MK2 infected H23 cells (red dash arrows). Additionally, a band is present toward the negative electrode on an isoelectric gradient in Ad-eGFP infected cells that is not present in Ad-MK2 infected cells (red dash circles). Treatment of cell lysates from Ad-MK2 infected H23 cells with active recombinant serine/threonine phosphatase, PP2A, reverses these charge based shifts, suggestive of a MK2-dependent phosphorylation event. **C.** H23 cells were infected with Ad-eGFP, Ad-WT-MK2, Ad-Active-MK2 (constitutively active mutant of MK2) or Ad-Dom Neg-MK2 (a dominant negative mutant of MK2) and 48hrs afterwards, cell lysates were analyzed for protein expression. As shown, there is significant phosphorylation of HSP27 (the canonical substrate of MK2) in Ad-WT-MK2 and Ad- Active-MK2 but not Ad-eGFP and Ad-Dom Neg-MK2 infected cells. **D.** H23 cells were infected with Ad-Dom Neg-MK2 or Ad-Active-MK2 and cell lysates underwent 2- Dimensional immunoblotting for caspase 3. As shown, there is a shift toward the positive electrode on an isoelectric gradient in Ad-Active-MK2 infected H23 cells that is not present in Ad-Dom Neg-MK2 infected cells (red dash arrows), suggestive of a MK2- dependent phosphorylation event. **E.** H23 cells were infected with Ad-eGFP or Ad-MK2 for 48 hours and cell lysates were treated with active recombinant PP2A or vehicle and analyzed for caspase 3/7 activity. As shown, there is no interaction between MK2 expression and PP2A treatment on caspase 3/7 activity. Furthermore, there is no within group effect of PP2A treatment on caspase 3/7 activity using post-hoc Tukey’s multiple comparisons test. N= 9-18 per group. **F.** H23 cells were infected with Ad-eGFP, Ad- Active-MK2 or Ad-Dom Neg-MK2 for 48hrs and cell lysates were analyzed for caspase 3 activity. As shown, both MK2 constructs resulted in significant increase caspase 3 activity compared to Ad-eGFP infected cells. Furthermore, there was no difference in caspase 3 activity when comparing Ad-Active-MK2 or Ad-Dom Neg-MK2. @, *P* < 0.05 vs all others. All post-hoc analyses using Dunn’s multiple comparisons test. N= 9 per group.

### Enzymatic Function of MK2 is Dispensable in Nuclear Translocation of Caspase 3 and Resultant Apoptosis

We have previously identified the presence of MK2 is required for nuclear translocation of caspase 3 and subsequent apoptosis(8). Having demonstrated that MK2 kinase activity does not alter caspase 3 activity, we sought to identify if MK2 kinase activity plays a role in nuclear translocation of caspase 3. Nuclear and cytosolic cellular fractions from H23 cells infected Ad-eGFP shows a significantly reduced nuclear caspase 3 activity compared to cytosolic caspase 3 activity, **Figure 8A**. Infection with Ad-WT MK2 resulted in increased nuclear compared to cytosolic caspase 3 activity, **Figure 8A**, suggesting nuclear translocation of caspase 3 with MK2 expression. To determine if this increased nuclear caspase 3 activity was functional, we measured cleavage of PARP1, a well identified nuclear substrate of caspase 3 critical in the execution of apoptosis(2, 34–36). As shown in **Figure 8B**, there was a marked increase in cleaved PARP1 in cell lysates from H23 cells infected with Ad-WT MK2 as compared to H23 cells infected with Ad-eGFP. To specifically test the role of MK2 kinase activity on nuclear translocation of caspase 3, H23 cells were infected with Ad-Active-MK2 and Ad-Dom-Neg-MK2 and nuclear and cytosolic cellular fractions were assessed for caspase 3 activity. As shown in **Figure 8C**, there was a significant increase in nuclear compared to cytosolic caspase 3 activity with both Ad-Active-MK2 and Ad-Dom-Neg- MK2 conditions; which resulted in increased PARP1 cleavage, **Figure 8D** and **Supplemental Figure 9**. Similar to our previous observations, increased nuclear caspase 3 activity also directly associated with increased cell death; specifically, H23 cells infected with Ad-WT MK2, Ad-Active-MK2 or Ad-Dom-Neg-MK2 all resulted in significantly higher cell death when compared to cells infected with Ad-eGFP, **Figure 8E**. Collectively, these data suggest phosphorylation of caspase 3 by MK2 and MK2’s kinase activity in general is dispensable with regards to MK2-mediated caspase 3 nuclear translocation and resultant cell death.

**Figure 8:**
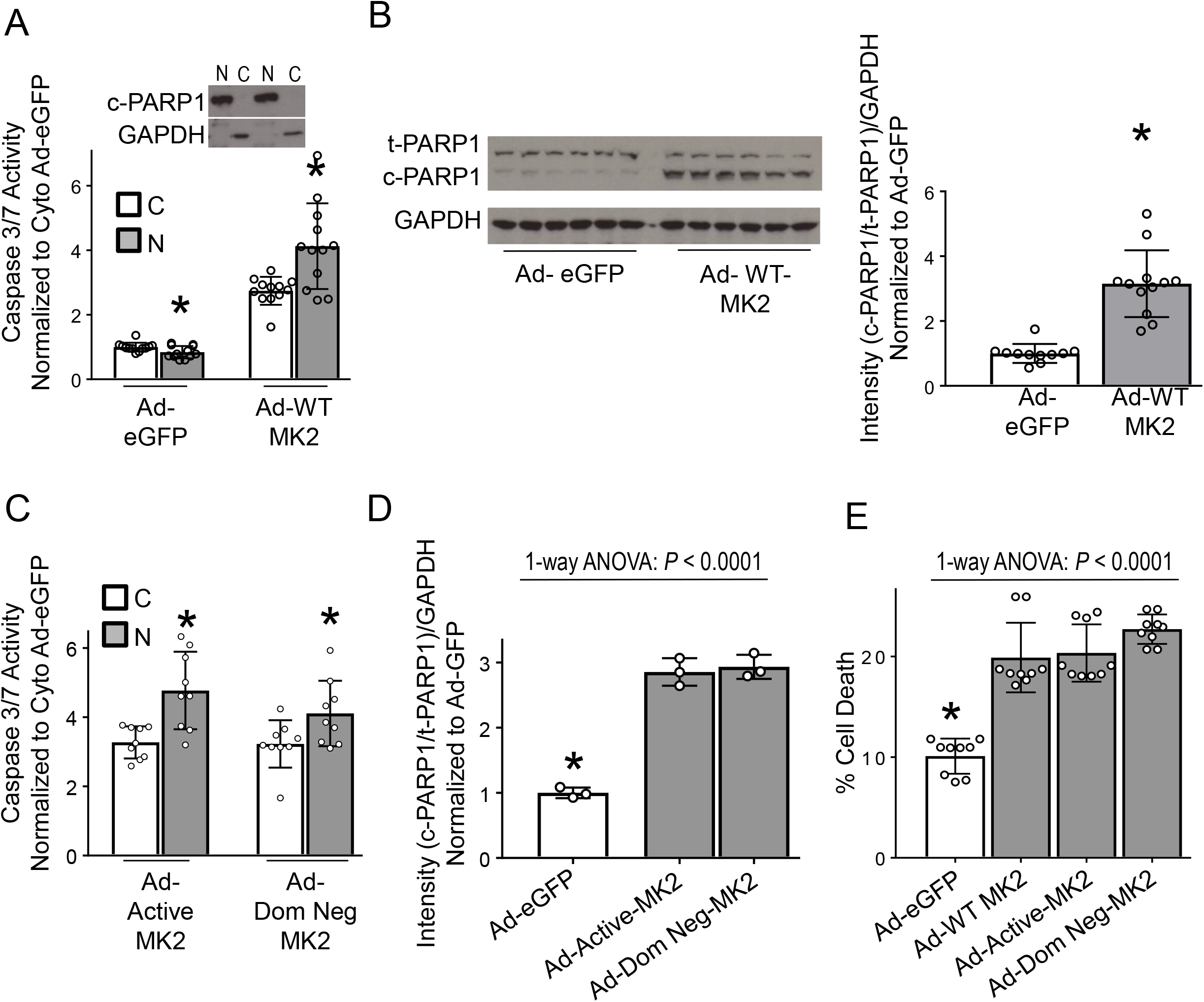
Enzymatic function of MK2 is dispensable in increasing nuclear caspase 3 activity and resultant apoptosis. **A.** H23 NSCLC cells were infected with Ad-eGFP or Ad-WT MK2 and 48hrs afterwards nuclear and cytosolic fractions were separated for analyses. H23 NSCLC cells infected with Ad-eGFP show a reduced nuclear caspase 3/7 activity compared to the cytosolic compartment. Whereas, H23 NSCLC cells infected with Ad-WT MK2 exhibited significantly increased nuclear caspase 3/7 activity compared to the cytosolic compartment. Purity of nuclear and cytosolic fractions was assessed by exclusion of GAPDH and cleaved PARP1 immunoblotting, respectively. N, nuclear. C, cytosolic. *, *P*< 0.05 vs cytosolic compartment by unpaired t test. N= 12 per group. **B.** Representative immunoblot of lysates from H23 cells infected with either Ad- eGFP2 or Ad-WT MK2 for 48hrs indicates an increase in cleavage of PARP1 in Ad-WT MK2 infected H23 cells compared to Ad-eGFP infected H23 cells, confirmed by densitometric analysis. *, *P*< 0.05 by unpaired t test. N= 11-12 per group. **C.** H23 NSCLC cells were infected with Ad-Active-MK2 or Ad-Dom Neg-MK2 and 48hrs afterwards nuclear and cytosolic fractions were separated for analyses. H23 NSCLC cells infected with either Ad-Active-MK2 or Ad-Dom Neg-MK2 exhibited significantly increased nuclear caspase 3/7 activity compared to its cytosolic compartment. N, nuclear. C, cytosolic. *, *P*< 0.05 vs cytosolic compartment by unpaired t test. N= 9 per group. **D.** H23 NSCLC cells were infected with Ad-eGFP, Ad-Active-MK2 or Ad-Dom Neg-MK2 and 48hrs afterwards lysates were immunoblotted for PARP1. H23 NSCLC cells infected with either Ad-Active-MK2 or Ad-Dom Neg-MK2 exhibited significantly increased cleavage of PARP1 compared to cells infected with Ad-eGFP. *, *P*< 0.05 vs all others in post hoc testing using Tukey’s multiple comparisons test. N= 3 per group. **E.** H23 NSCLC cells were infected with Ad-eGFP, Ad-WT-MK2, Ad-Active-MK2 or Ad- Dom Neg-MK2 for 48hrs and were then analyzed for cell death using flow cytometry. H23 NSCLC cells were infected with Ad-WT-MK2, Ad-Active-MK2 or Ad-Dom Neg-MK2 resulted in significantly higher amount of cell death compared to H23 cells infected with Ad-eGFP. *, *P*< 0.05 vs all others in post hoc testing using Tukey’s multiple comparisons test. N= 9 per group.

### MK2 Associates with Caspase 3 and is Required for Increased Nuclear Caspase 3 Activation and Resultant Apoptosis

We hypothesized that the effects of MK2 on caspase 3 nuclear translocation and cell death may be due to a non-enzymatic relationship between MK2 and caspase 3. MK2 is unique in that it has both a nuclear localization sequence (NLS) and a nuclear export sequence (NES)(13). Since caspase 3 lacks an NLS, we reasoned that caspase 3 may associate with MK2 in order to gain access to the nucleus. Normally, MK2 resides in the nucleus as it has a continuously active NLS. However, when activated, MK2 reveals its NES thereby enabling shuttling back and forth between the nucleus and cytoplasm(13, 37). Therefore, we hypothesized that MK2 may be serving as an escort protein for caspase 3 to translocate to the nucleus (via NLS). We first sought to determine if caspase 3 came in close proximity to MK2 using BioID based assays. Immunoblotting of H23 cells infected with Ad-WT MK2-BioID-C or Ad-WT MK2-BioID-N demonstrates MK2 at an appropriately increased molecular weight and with its kinase function intact, as evidenced by phosphorylation of HSP27 (**Supplemental** **Figures 10** **A-B**). As shown in **Figure 9A**, caspase 3 is preferentially detected following pulldown with streptavidin when the biotin ligase is fused to the c-terminus of MK2, suggesting caspase 3 comes in proximity to MK2. We next generated MK2 constructs with a mutated nuclear localization sequence MK2, Ad-Mut-NLS-MK2, and a mutated nuclear export sequence MK2, Ad-Mut-NES-MK2, as previously described(13–15). As shown in **Figure 9B**, there was no MK2 expression in the nuclear fraction of H23 cells infected with Ad-Mut-NLS-

**Figure 9:**
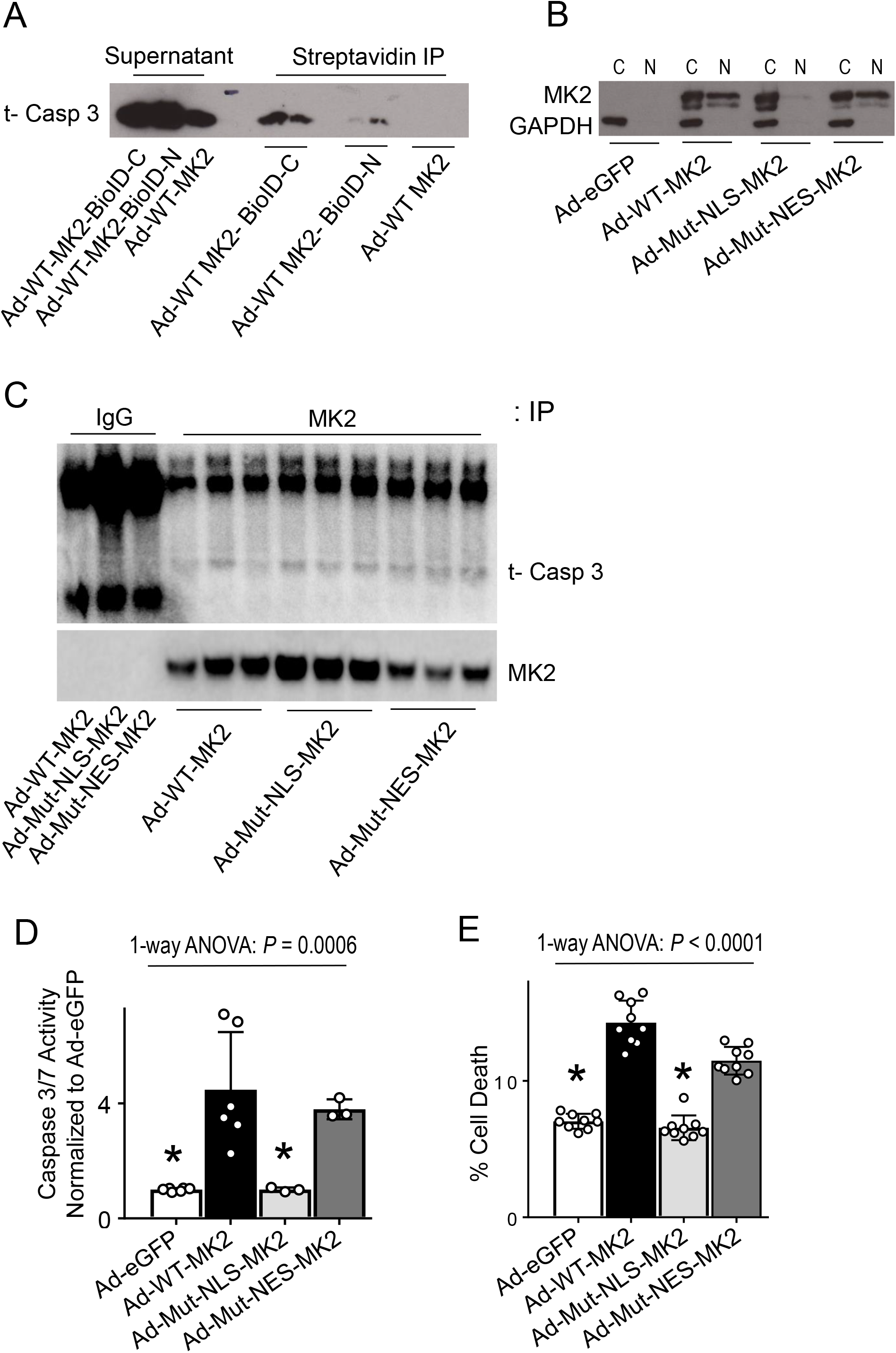
MK2 associates with caspase 3 and MK2’s nuclear translocation is required for increased caspase 3 activation and resultant apoptosis. **A.** H23 cells were infected with Ad-WT-MK2, Ad-WT MK2-BioID-C (MK2 fused to biotin ligase on the c-terminus) or Ad-WT MK2-BioID-N (MK2 fused to biotin ligase on the n-terminus) and following incubation with biotin, cell lysates were precipitated with beads conjugated to streptavidin and then immunoblotted using antibodies directed against caspase 3. As shown, there is increased biotinylated caspase 3 with biotin ligase fused to MK2 on the c-terminus. **B.** H23 cells were infected with Ad-eGFP, Ad-WT-MK2, Ad-Mut-NLS-MK2 (mutated nuclear localization sequence, rendering cytosolic sequestration), Ad-Mut- NES-MK2 (mutated nuclear export sequence, limiting nuclear export) and 48hrs afterwards nuclear and cytosolic fractions were separated for analyses. Representative immunoblot shows wild-type and mutant NES-MK2 constructs translocate to the nuclear compartment, whereas the mutant NLS-MK2 construct is sequestered in the cytosolic fraction and unable to translocate to the nucleus. **C.** H23 cells were infected with Ad- WT-MK2, Ad-Mut-NLS-MK2 (mutated nuclear localization sequence, rendering cytosolic sequestration), Ad-Mut-NES-MK2 (mutated nuclear export sequence, limiting nuclear export) and 48hrs afterwards cell lysates were immunoprecipitated with antibodies against MK2 and then immunoblotted using antibodies directed against caspase 3. As shown, caspase 3 is associated with MK2. There is no difference of association based on localization mutations. **D.** H23 cells were infected with Ad-eGFP, Ad-WT-MK2, Ad- Mut-NLS-MK2 and Ad-Mut-NES-MK2 and 48hrs afterwards cell lysates were analyzed for caspase 3 activity. Cells infected with wild-type and mutant NES-MK2 constructs exhibited a significant increase in caspase 3/7 activity compared with H23 cells infected with Ad-eGFP or Ad-Mut-NLS-MK2. *, *P*< 0.05 vs Ad-WT-MK2 and Ad-Mut-NES-MK2 in post hoc testing using Tukey’s multiple comparisons test. N= 3-6 per group. **E.** H23 cells were infected with Ad-eGFP, Ad-WT-MK2, Ad-Mut-NLS-MK2 and Ad-Mut-NES-MK2 and 48hrs afterwards cells were analyzed for cell death using flow cytometry. Cells infected with wild-type and mutant NES-MK2 constructs exhibited a significant increase in cell death compared with H23 cells infected with Ad-eGFP or Ad-Mut-NLS-MK2*, *P*<0.05 vs Ad-WT-MK2 and Ad-Mut-NES-MK2 in post hoc testing using Tukey’s multiple comparisons test. N= 9 per group.

MK2, compared to Ad-WT-MK2 and Ad-Mut-NES-MK2 infection, where MK2 was seen in both the nuclear and cytosolic fractions. While MK2 and caspase 3 are in close proximity to each other, in order for MK2 to serve as an escort for caspase 3, MK2 would have to complex with caspase 3; thus, we sought to determine if these two proteins are in fact associated, using co-immunoprecipitation assays. As shown in **Figure 9C**, caspase 3 could be detected following pulldown with MK2. Interestingly, we observed no notable difference in amount of caspase 3 detected between cells infected with Ad-WT MK2, Ad-Mut-NLS-MK2 or Ad-Mut-NES-MK2, suggesting these mutations do not impact MK2’s association with caspase 3 (**Figure 9C**). We next sought to determine if prevention of nuclear trafficking of MK2 would protect against caspase 3 mediated apoptosis. As shown in **Figure 9D**, infection of H23 cells with Ad-WT MK2 led to a significant increase in caspase 3 activity, which is prevented when the nuclear localization sequence of MK2 is mutated, as with Ad-Mut-NLS-MK2. While MK2 kinase activity is not required for caspase 3 activation (**Figure 6B**), we noted that mutating the NLS of MK2 also did not impact its kinase activity, as evidenced by its ability to phosphorylate HSP27 (**Supplemental Figure 10C**). Additionally, preventing nuclear translocation of MK2, as with Ad-Mut-NLS-MK2, completely protected against MK2 expression induced apoptosis in H23 cells, **Figure 9E**. These data suggest that MK2 functions as a molecular chaperone for caspase 3 during nuclear translocation of both proteins and resultant caspase 3 mediated apoptosis.

### High Tumor MK2 Expression is Associated with Survival Only in Lung Adenocarcinoma

To determine whether the observed effects of MK2 on cell death and patient survival were specific to lung adenocarcinoma, we examined the effect of MK2 on overall survival across multiple TCGA datasets. For this analysis, we pre-specified the multivariate Cox PH model as follows: MK2 was specified as high/low (as in our previous analyses), and we included cancer stage (defined as early vs. late), gender, smoking and age as relevant covariates. Similar to our prior analysis, we again censored the data at 2 years. Since we were applying a Cox PH model across multiple datasets, we also extracted metrics of model fit, including overall goodness of fit (i.e. the p value associated with the Wald test for the Cox PH model) and PH assumption testing (i.e. p value associated proportionality hazard assumption testing, where p < 0.05 signifies violation of hazard assumption). In **Figure 10 A-B**, we show not only the HR for high MK2 in the multivariate model across datasets, but also the relative performance of our pre-specified model in each dataset. As shown in **Figure 10A**, the LUAD dataset was unique in that MK2 was associated with a significant lower HR for death. We note that the model fit for LUAD was good, as evidenced by the fact that a) the MK2 variable did not violate PH assumption and b) a significant goodness of fit (wald p value < 0.05), **Figure 10B**. Interestingly, in kidney renal clear cell cancer (KIRC), high MK2 transcript expression appeared to be associated with worse survival. However, when comparing MK2 mRNA and MK2 protein expression in renal cell carcinoma cell lines, we observed a negative correlation, which was not statistically significant, **Figure 10C**.

**Figure 10:**
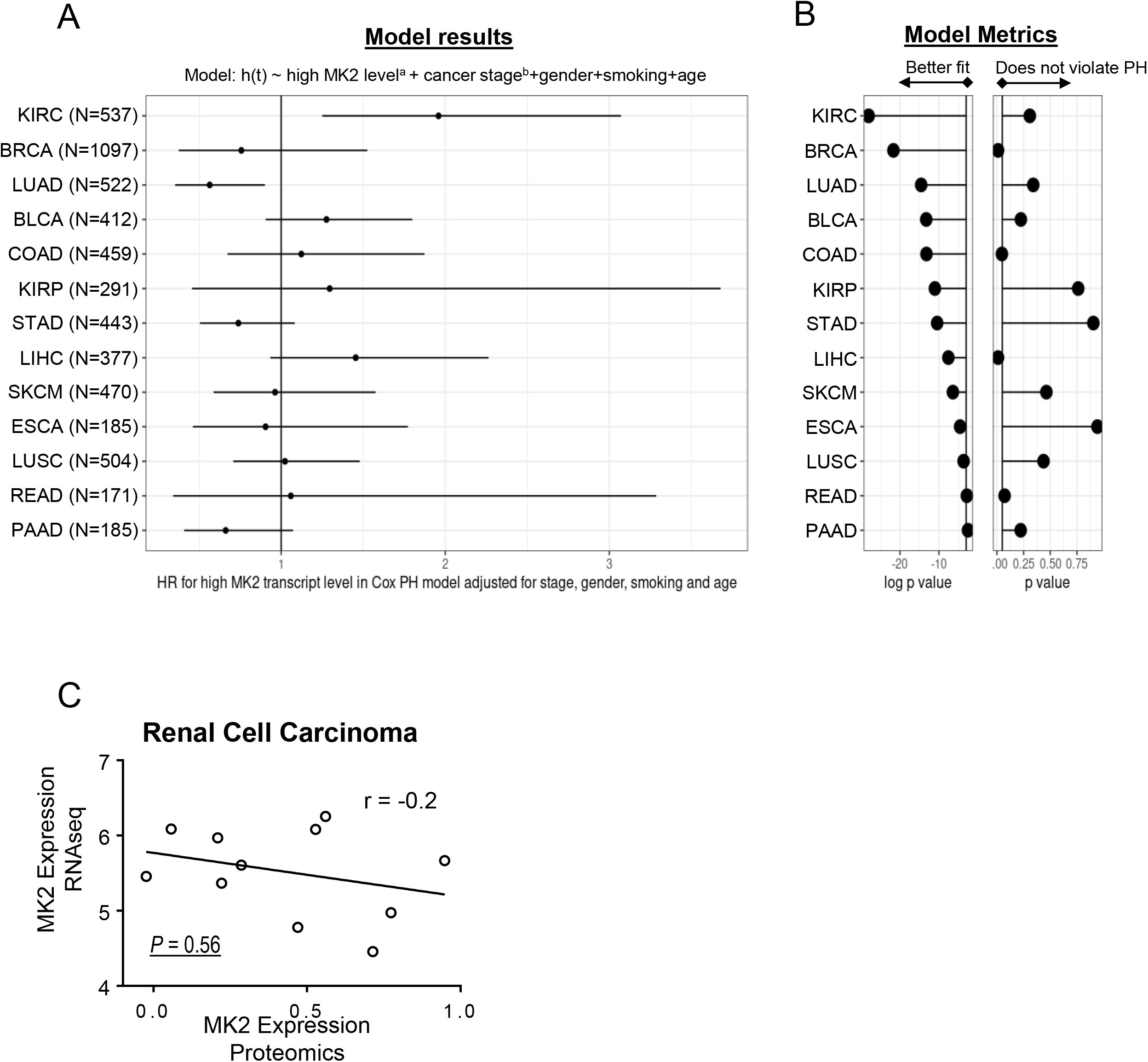
High tumor MK2 expression is associated with improved survival only in lung adenocarcinoma. **A.** Plot showing HR estimates and 95% CI results for a cox proportional hazards model assessing the effect of high MK2 expression^a^ on overall survival at 2 years. Vertical line represents HR of 1. Model: h(time to death | covariates) ∼ high MK2 level^a^ + cancer stage^b^+gender+smoking+age. ^a^ MK2_level defined dichotomously as 0: Bottom 2/3rd of log normalized MK2 transcript levels for that cancer type; 1: Top 1/3. ^b^ Cancer stage defined as: Early Stage (Stage I, Stage II) vs. late stage (Stage III, Stage IV). **B.** Lollipop plots showing Cox PH model metrics by cancer type. Vertical line represents p value of 0.05. *Left:* Dots represent the log p value for goodness of fit (Wald test) for each Cox PH model (p < 0.05 represents significant). *Right:* Dots represents results of proportionality hazard testing p value (p > 0.05 represents no violation of PH assumption). **C.** Renal cell carcinoma cell lines examined by the Cancer Dependency Map project (www.depmap.org) shows a non- significant negative correlation between MK2 mRNA and MK2 protein expression. KIRC, kidney renal cell carcinoma. BRCA, breast invasive carcinoma. LUAD, lung adenocarcinoma. BLCA, bladder urothelial carcinoma. COAD, colon adenocarcinoma. KIRP, kidney renal papillary cell carcinoma. STAD, stomach adenocarcinoma. LIHC, liver hepatocellular carcinoma. SKCM, skin cutaneous melanoma. ESCA, esophageal carcinoma. LUSC, lung squamous cell carcinoma. READ, rectum adenocarcinoma. PAAD, pancreatic adenocarcinoma.

## DISCUSSION

A specific role for nuclear translocation of caspase 3 in apoptosis has been postulated based on transfection of active/cleaved subunits of caspase 3 fused to a NLS moiety resulting in massive apoptosis of transfected cells(38). Furthermore, apoptosis induced by microinjection of active/cleaved caspase 3 was prevented by co-injection with wheat germ agglutinin, a compound known to inhibit all nuclear transport, suggesting that interfering with caspase 3 nuclear translocation prevents apoptosis, despite activation of the apoptotic pathway(39). The classic paradigm for active nuclear transport involves the binding of a NLS region within a target protein to a specific family of soluble karyopherin α(40)carrier proteins, which in turn facilitate nuclear transport of the target protein via the nuclear pore complexes(41). Since caspase 3 does not have an identifiable NLS, an alternative mechanism for nuclear translocation of caspase 3 must exist. Our data show an association between MK2 (which has an NLS) and caspase 3, suggesting MK2 functions as a molecular chaperone for caspase 3 during nuclear translocation and resultant caspase 3 mediated apoptosis. While our data clearly show an association between MK2 and caspase 3, it is not clear if MK2 directly binds to caspase 3 or requires an intermediary scaffolding protein. Being a serine/threonine kinase, MK2 is primarily thought to phosphorylate substrates in order to initiate signaling cascades or alter mRNA stability(42–45); however, enzymes, and kinases in particular, can also function as chaperones(46–49). To date, there have been mixed results regarding the chaperoning function of MK2. Specifically, MK2 complexes with p38 and co-localizes intracellularly with p38 during nuclear export following stimulation with arsenite(15), suggesting a potential chaperoning effect of MK2. In contrast, in other experiments, leptomycin B prevented the nuclear export of MK2 but not p38(50), suggesting separate pathways. HSP27, a downstream substrate for MK2(12, 37, 42, 51), is a well described molecular chaperone (52) and binds directly to caspase 3(53). Further, the phosphorylation of HSP27 alters its cellular localization, with phospho- HSP27 translocating to the nucleus (54, 55) and non-phosphorylatable mutants of HSP27 remaining in the cytosol even after stimulation(55). That said, it seems unlikely phosphorylation of HSP27 is critical for caspase 3 nuclear translocation in NSCLC models, because our data show that when the NLS of MK2 is mutated, there is no increase in caspase 3 activation or cell death despite ample HSP27 phosphorylation (**Supplemental Figure 8A** and **Figure 9C** & **D**). To our knowledge, a direct chaperone function for MK2 has not previously been described.

It is interesting that expressing WT MK2 in H23 cells increased caspase 3 activity, especially given the fact that MK2 is not required for activation of caspase 3, as we have shown previously (8) and again in **Figure 6C**. It is unlikely that MK2 expression alone is cytotoxic as infection with Ad-Mut-NLS-MK2 does not increase caspase 3 activity. The most likely explanation is that potential inflammatory insult following adenoviral infection, or basal endogenous stressors commonly present in cancers(56, 57), coupled with a chaperoning function of MK2 via its NLS, leads to nuclear translocation of caspase 3 and resultant DNA damage, further propagating the apoptotic signaling cascade(58), i.e. increased caspase 3 activation. This hypothesis is supported by our data showing infection with Ad-WT MK2 results in phosphorylation of HSP27 (**Figure 7C** & **Supplemental Figure 8A**), indicating activation of stress-induced pathways(59, 60). Similar stress is noted following infection with Ad-Mut-NLS-MK2 (**Supplemental Figure 8A**) but does not result in increased caspase 3 activity (**Figure 9C**).

Overall, these mechanistic data provide biological rationale for the finding of reduced MK2 expression in late stage compared to early stage NSCLCs (**Figure 4A**). The logical extension of this mechanistic finding of reduced MK2 expression leading to resistance to cell death, resulting in more advanced stage NSCLC is intriguing but may not be transferable to other cancer types(61–63). For example, Henriques et al, demonstrate that intestinal carcinogenesis was related to MK2 kinase activity as chemical inhibition of MK2 led to decreased epithelial proliferation and reduced tumor growth(63). Interestingly, the majority of data showing a role for MK2 in cancers focus on MK2 kinase activity and its role in tumor development(61–64). In a novel murine model of NSCLC that generates MK2-proficient and MK2-deficient tumors within the same animal, Morandell et al show that MK2-proficient tumor area is consistently greater than MK2-deficient tumor area(65). One possible explanation for our data diverging from previously published work is that many of these utilized different cancer types (i.e. intestinal tumors) or murine models(61–65). However, in patient derived data from TCGA, we note a significant association with improved 2-year survival in lung adenocarcinoma with high MK2 expression, **Figure 5** & **10**. The quality of model fit was variable across cancer datasets within TCGA, **Figure 10** **C**; thus, it is possible better model fitting within those datasets, while also accounting for differences in the correlation between MK2 transcript and protein expression, may reveal additional signals linking MK2 to survival. It is also possible that MK2 may have divergent roles as it relates to tumor development and mechanisms of apoptosis resistance, the focus of our studies.

Our data clearly show that a non-kinase function of MK2 is responsible for the observed sensitization to cell death, **Figure 9**. These findings suggest that the association between increased MK2 mRNA transcript levels and better clinical outcomes in patients with early stage NSCLCs is likely driven by increased MK2 protein expression, supported by the fact that in NSCLC cell lines, MK2 protein expression is directly correlated to MK2 mRNA transcript levels, **Supplemental Figure 4**. The gene encoding MK2 is located on chromosome 1 with a chromosomal location of 1q32.1. We sought to determine if methylation was a potential regulator of MK2 transcript levels. However within the TCGA-LUAD methylation data, there was poor probe coverage surrounding the MK2 gene. Additionally, in our validation cohort(33) associated methylation data was not readily available. The mechanisms by which MK2 mRNA expression is regulated are not clear.

Our studies have several important limitations that need to be considered. First, one would predict that increased MK2 expression would result in improved clinical response to chemotherapy. It is possible that different chemotherapy regimens may have differential responses based on MK2 expression. Furthermore, the interaction between different oncogenic driver mutations and level of MK2 on chemotherapy responses may be different. Unfortunately, we were unable to discern these clinical variables using data from TCGA-LUAD or our validation cohort. However, we note our *in vitro* data suggests increased sensitivity to etoposide in cells expressing MK2. Specifically, H23 cells infected with Ad-WT MK2 and exposed to etoposide resulted in a marked increase in cell death compared to H23 cells infected with Ad-eGFP and exposed to etoposide,

**Supplemental Figure 9.** Additionally, we note that the clinical association of higher MK2 expression with improved two-year survival in lung adenocarcinoma was observed in two separate clinical cohorts that have distinct demographic and likely distinct oncogenic tumor driver mutation profiles, suggesting an independent role for MK2.

In summary, this study provides mechanistic insight into how MK2 promotes nuclear translocation of caspase 3 leading to PARP1 cleavage and execution of cell death. Specifically, the non-kinase function of MK2, nuclear trafficking via its NLS, is required for caspase 3 mediated cell death. This study suggests that NSCLCs may evade cell death by down regulating MK2 expression thereby reducing nuclear caspase 3 activity and apoptosis.

---

**Supplemental Figure 1.**
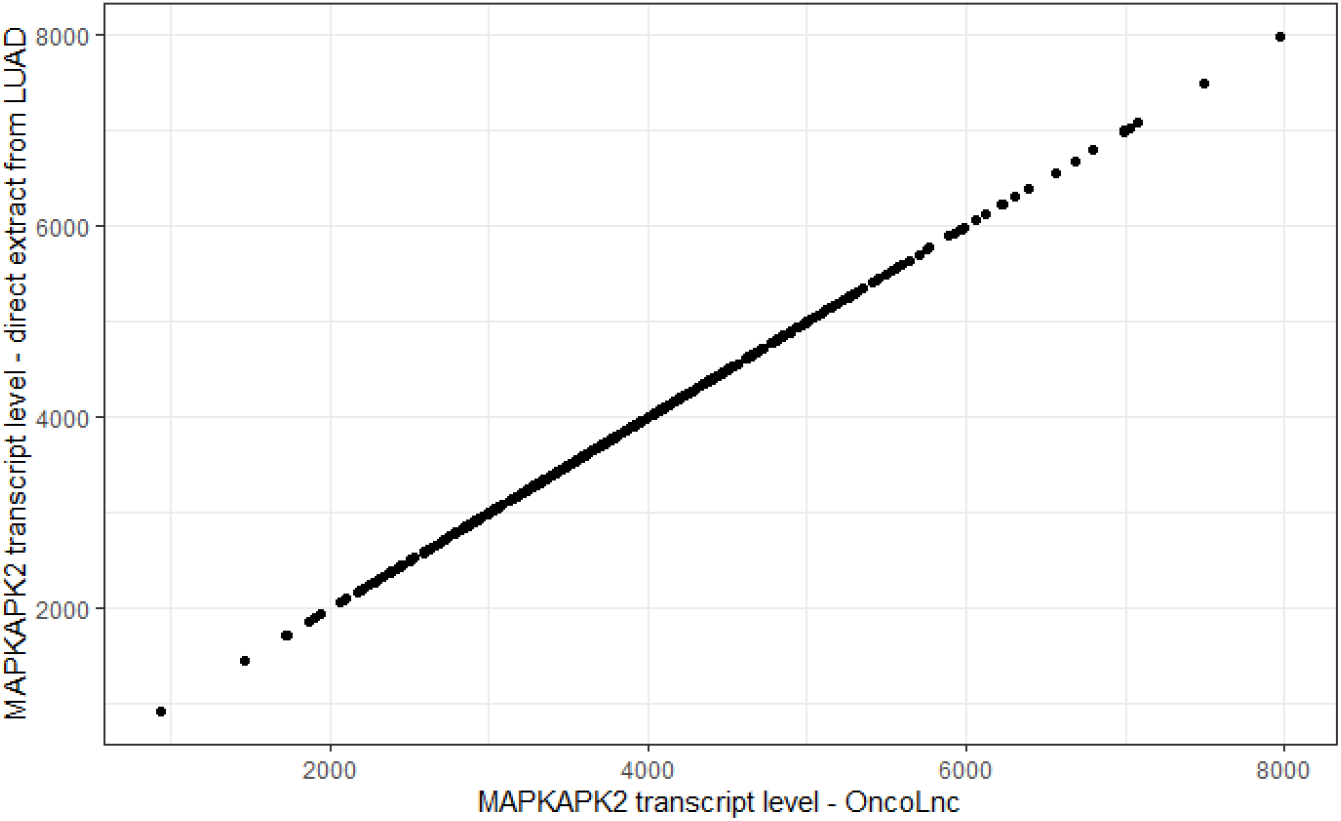

**Supplemental Figure 2.**
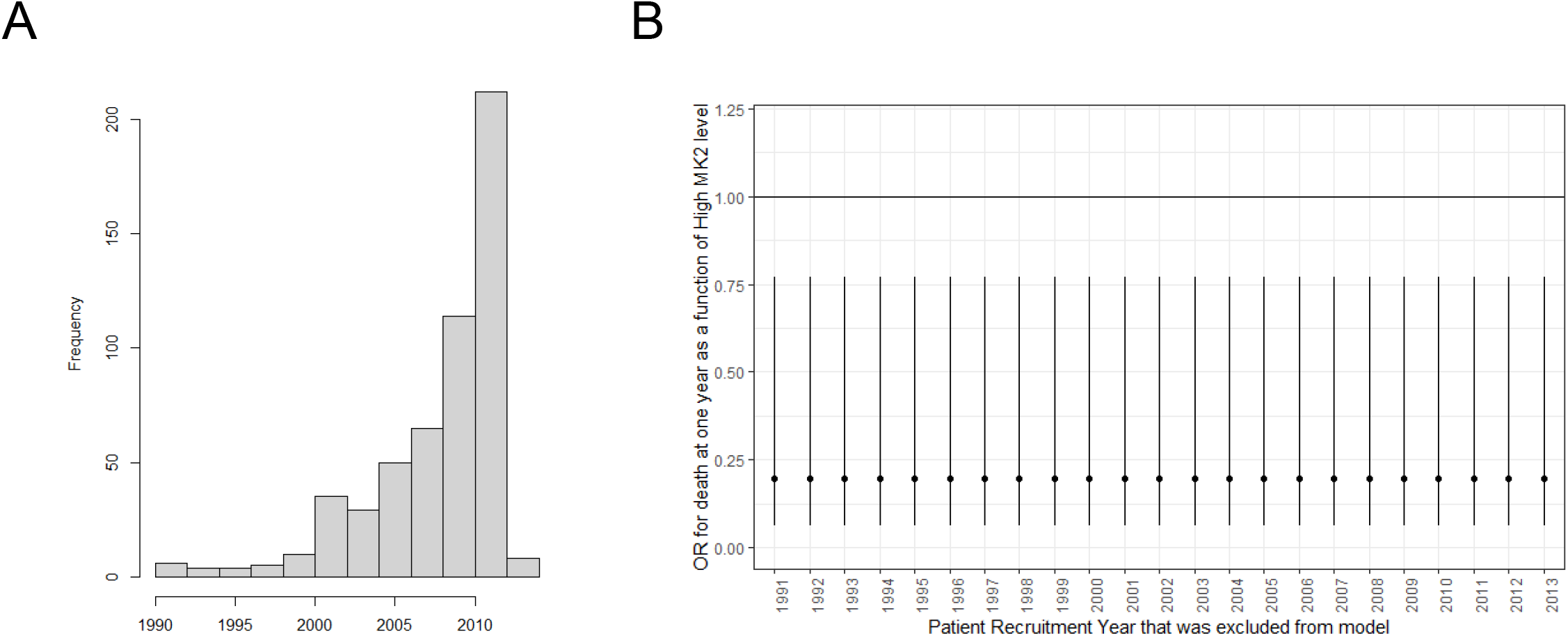

**Supplemental Figure 3.**
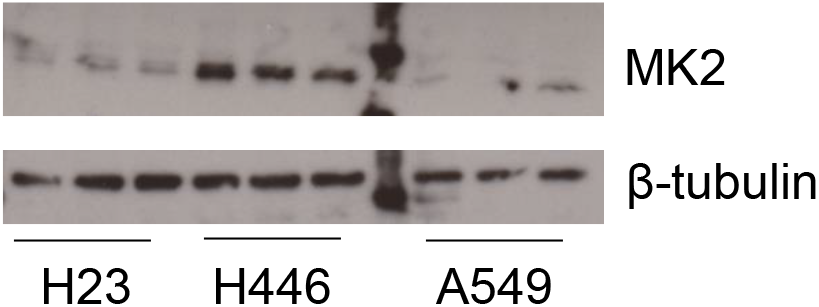

**Supplemental Figure 4.**
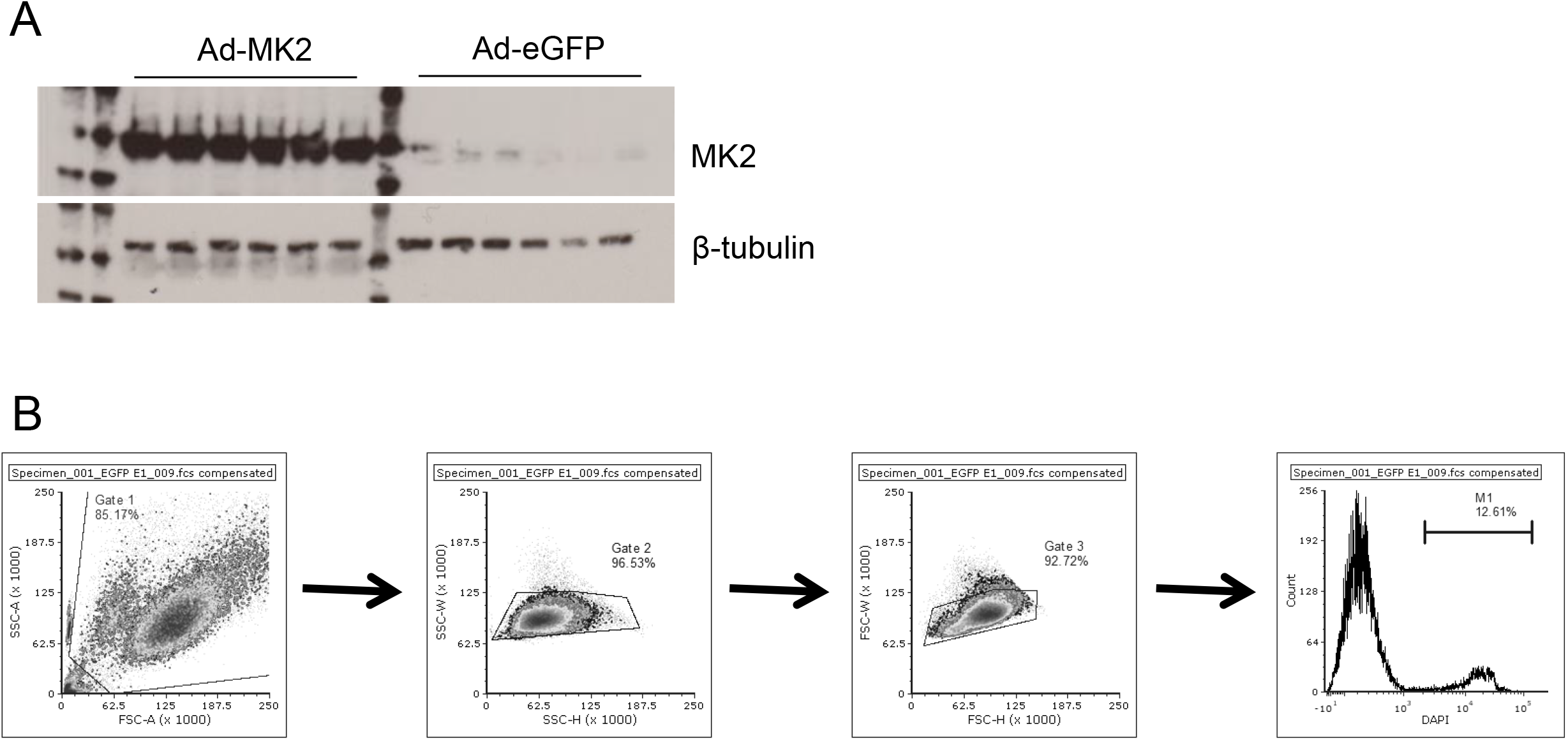

**Supplemental Figure 5.**
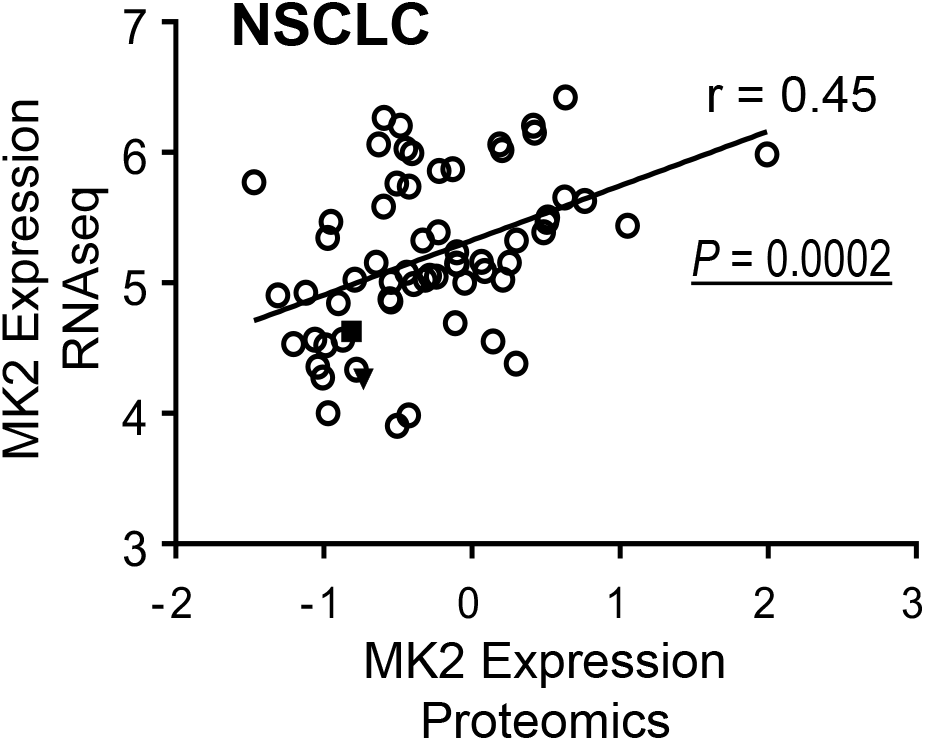

**Supplemental Figure 6.**
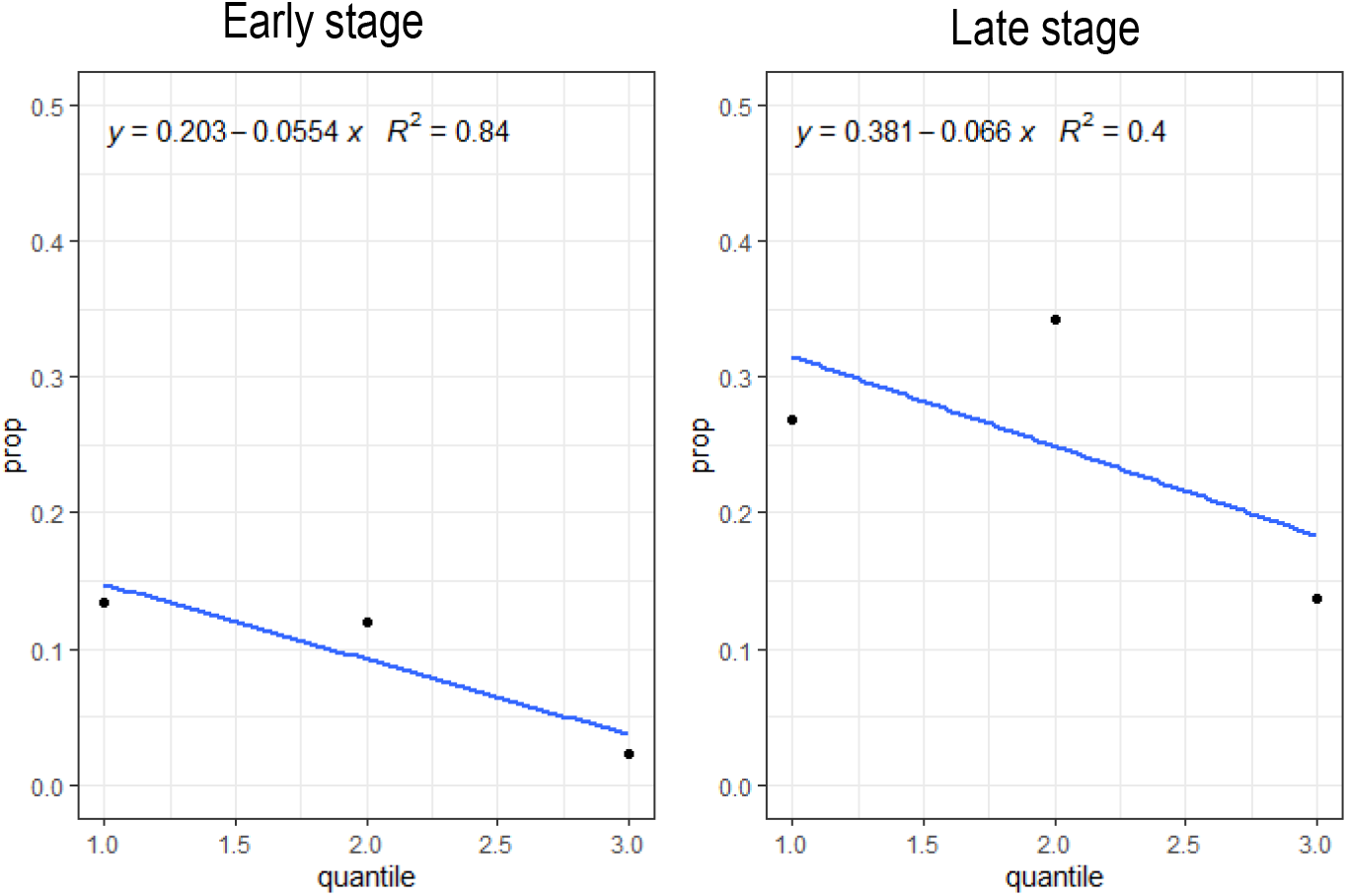

**Supplemental Figure 7.**
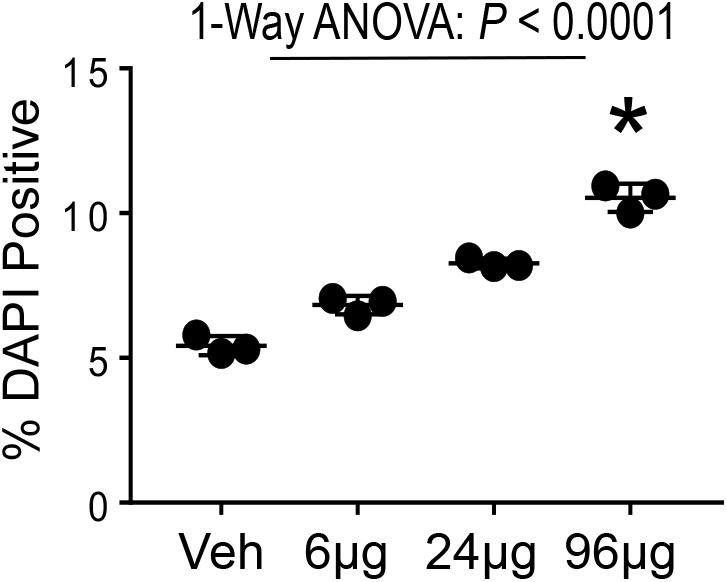

**Supplemental Figure 8.**
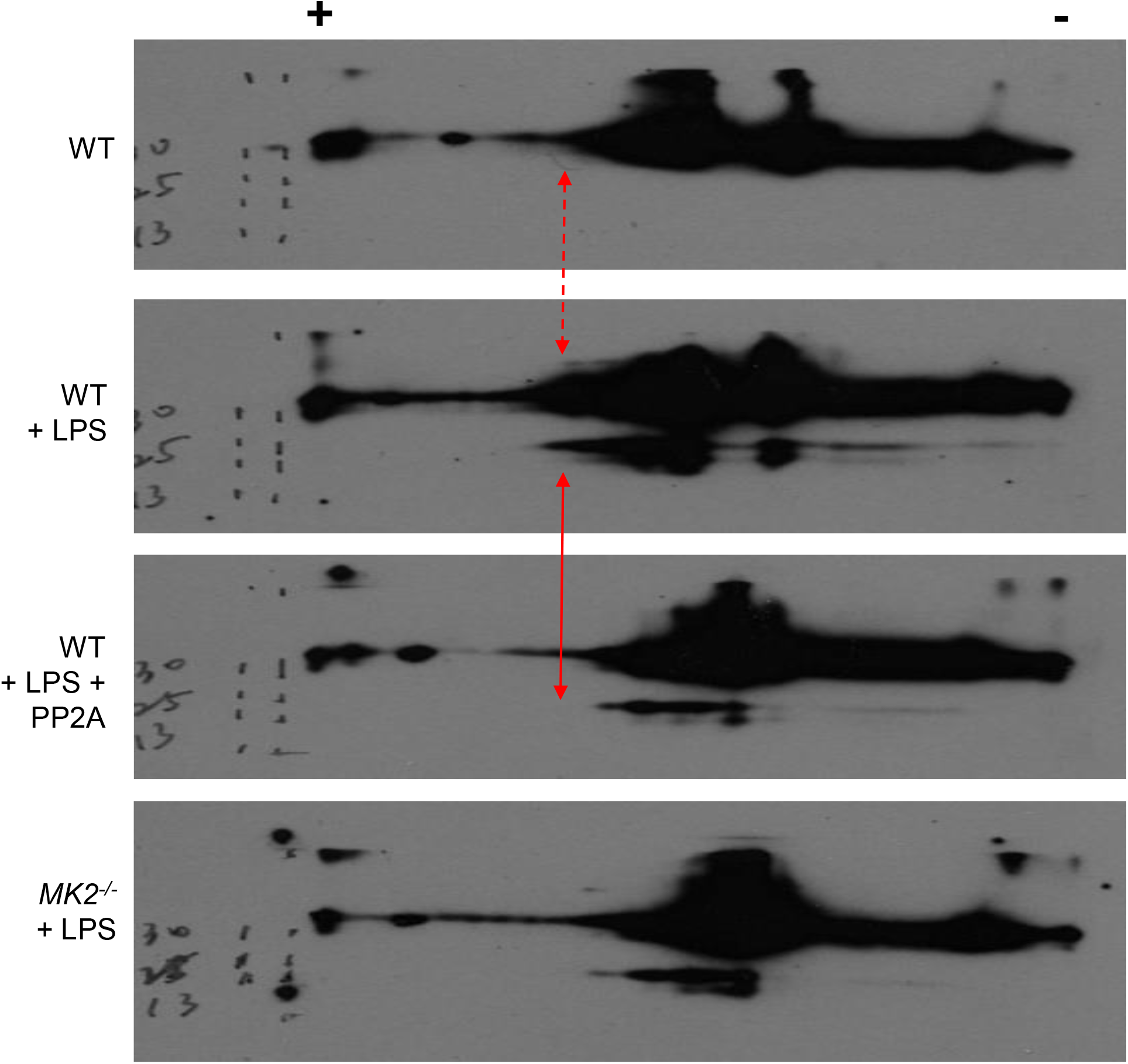

**Supplemental Figure 9.**
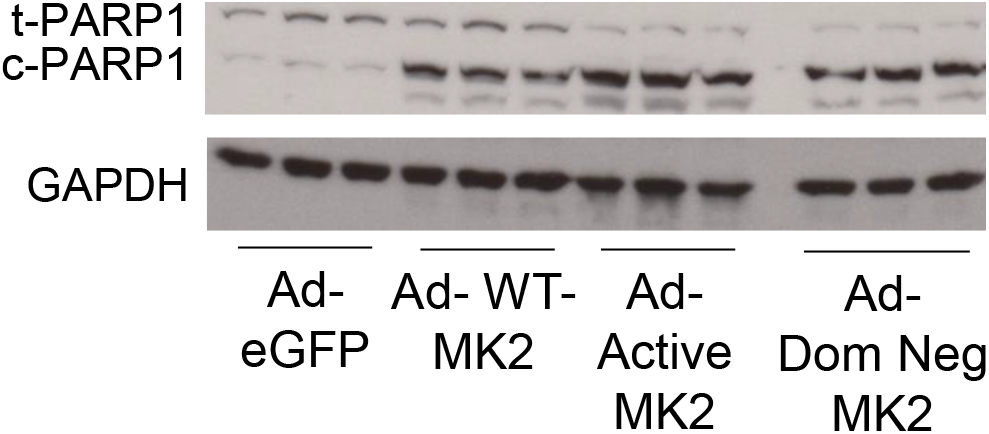

**Supplemental Figure 10.**
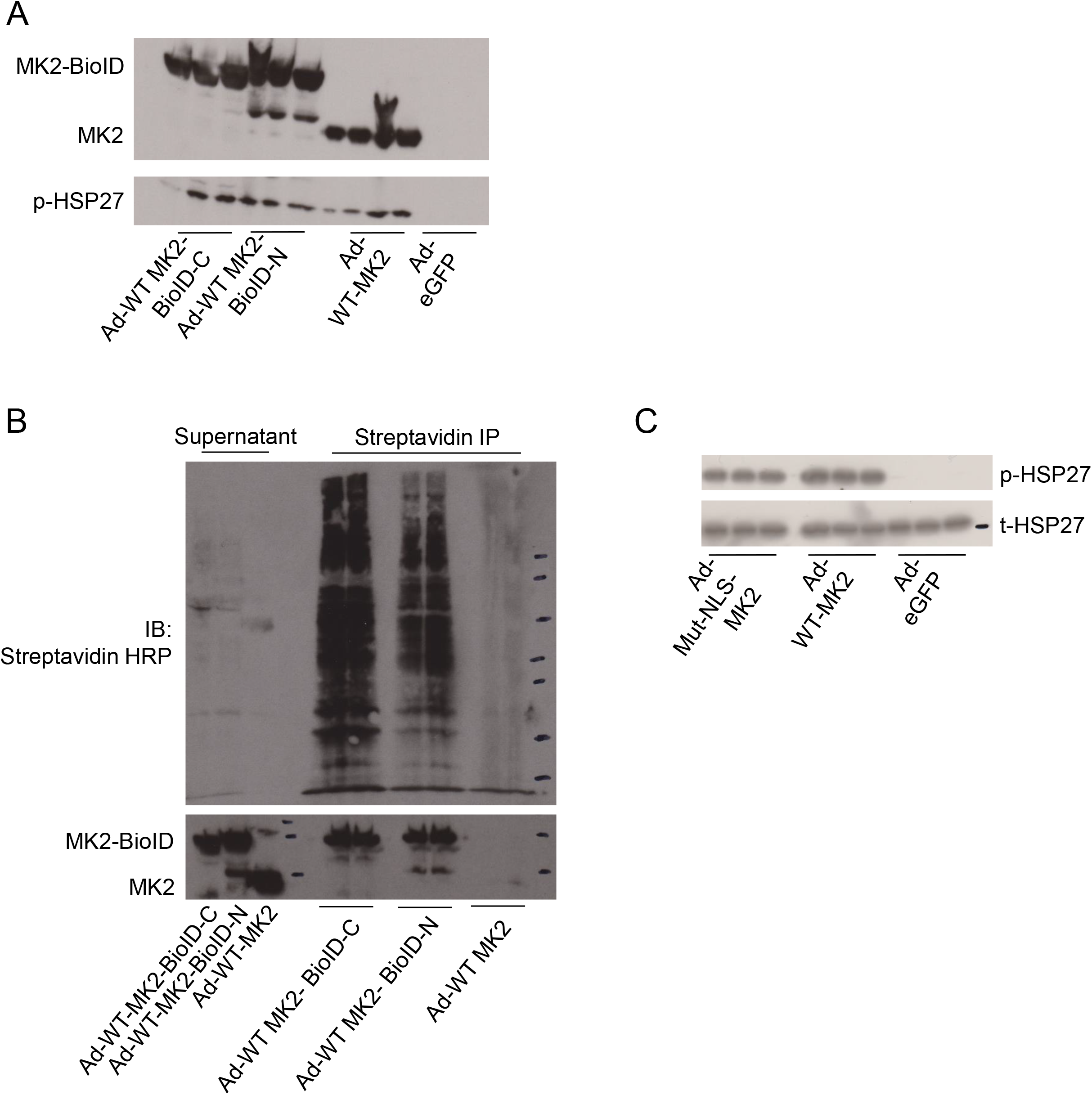

**Supplemental Figure 11.**
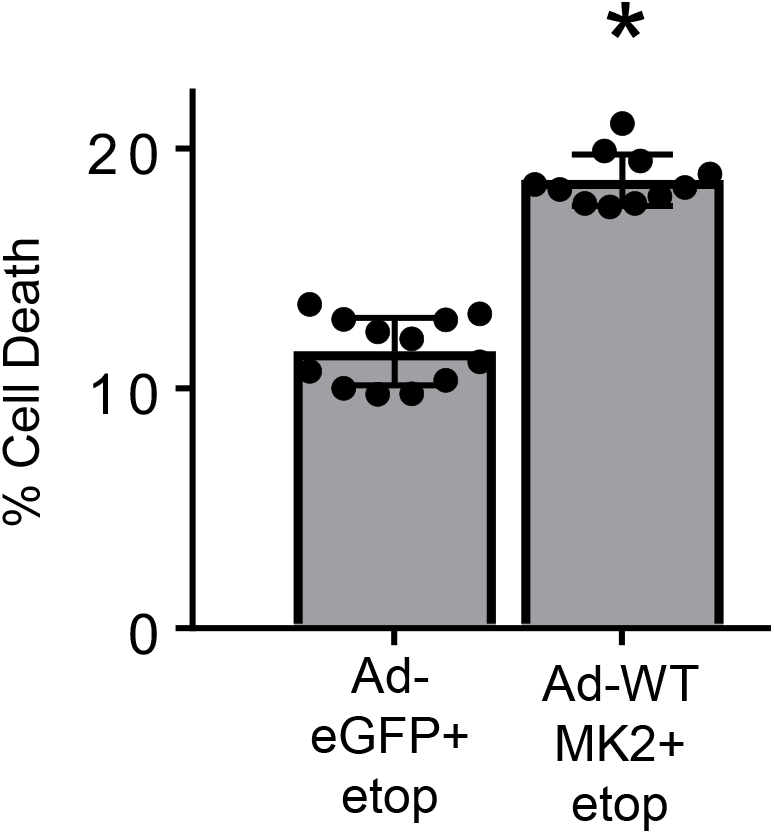

