## Supplemental Methods for "MK2 Expression Promotes Non-Small Cell Lung Cancer Cell Death and Predicts Survival"

* Contribute equally to this manuscript.

The Johns Hopkins University Institutional Animal Care and Use Committee approved all animal protocols.

Male C57BL/6J (wild type, WT) mice aged 10-12 weeks (Jackson Laboratory, Bar Harbor, ME) and *MK2*$^{-/-}$ mice, C57BL/6J background (1) were exposed to intravenous (IV) PBS or lipopolysaccharide (LPS, 0127:B8, product # L3129, Sigma) via retro-orbital injection (2) for up to 6hrs. After exposure to the experimental conditions, lungs were flushed free of blood, removed and homogenized in cell lysis buffer. Lung lysates were frozen by immersion into liquid nitrogen and subsequently stored for later analyses.

**Cell Lines**

Non-small cell lung carcinoma (NSCLC) cell lines- H23 and A549 cell lines were purchased from ATCC (Manassas, VA). Small cell lung carcinoma (SCLC) cell line- H446 was a kind gift from Dr. Phil Dennis. H23 and H446 cells were cultured in RPMI 1640 media (ThermoFisher A1049101) supplemented with 10% (v/v) FBS (Hyclone). A549 cells were cultured in FK12 medium (ThermoFisher 21127022) supplemented with 10% (v/v) FBS (Hyclone). Cells were maintained in full growth media in 75-cm$^2$ flasks. Cells were grown at 37 C with 5% $CO_2$.

Adenoviral vectors: adenoviral vectors encoding wild-type MK2 (Ad-WT MK2), constitutively active MK2 (Ad-Active-MK2; T222E, T334E)(3), dominant negative MK2 (Ad-Dom Neg-MK2; K93R)(4), a mutated nuclear export sequence MK2 (Ad-Mut-NES; L360A) (5), a mutated nuclear localization sequence MK2 (Ad-Mut-NLS-MK2; K372A,

K373A, K388A, K389A) (6, 7), a wild-type MK2 fused to a biotin protein ligase on the c-terminus (Ad-WT-MK2-BioID-C) or the n-terminus (Ad-WT-MK2-BioID-N) were directly purchased from Vector Builder (Chicago, IL). The sequences of the plasmids encoding these vector were verified by Sanger sequencing (The Genetics Resources Core Facility, Johns Hopkins University).

Infections: Cells were seeded in 6-well plates at a cell density of 5x10^5 cells per well. Cells were grown at 37 C with 5% $CO_2$ for approximately 6 hours to allow adherence. After adherence, media was replaced with media plus viral vector. Cells were incubated with viral vector to have a final plaque-forming unit number of up to 100. Cells were left to incubate with the viral media at 37 C with 5% $CO_2$ for approximately 24 hours. Viral media was then replaced with the appropriate subsequent media, depending on experimental conditions.

Immunoblot analyses: Cell cultures were lysed using cell lysis buffer (CST 9803s, Cell Signaling, Boston, MA) supplemented with protease inhibitors cocktail (Sigma P8340), PMSF 1mM, Thermo Fischer 36978), NaF (1mM, Sigma, 201154) and NaOV (1mM, Sigma, S6508). Protein lysates were denatured using Laemmli Sample buffer (BioRad 1610747), 2-Mercaptoethanol (Millipore Sigma M6250), and 100 C heat (5-minute exposure). Proteins were separated by SDS-PAGE (Thermo Fisher XP00122BOX), and transferred to PVDF membranes (BioRad 1620177). Membranes were blocked in 5% non-fat dry milk (BioRad 1706404) in TBS (Quality Biological 50983267) with 0.5%

Tween-20 (Thermo Fisher BP337-500). Membranes were incubated with primary antibodies at 1:1000 dilution overnight in 2.5% non-fat dry milk. PVDF membranes were then incubated with horse radish peroxidase-linked secondary antibodies, anti-mouse (CST 7076) or anti-rabbit (CST 7074), at 1:5000 dilution for 1 hour in 1% non-fat dry milk. Immunoblots for Streptavidin conjugated to horse radish peroxidase utilized bovine serum albumin instead of nonfat dry milk. Protein bands were visualized using chemiluminescent detection methods. Band intensities were quantified using ImageJ software.

Phospho-specific anti-total antibodies directed at HSP27(p-HSP27- CST-2401; t-HSP27- CST-2402) and anti-total antibodies directed at MK2 (CST-3042), caspase 3 (CST-9662), β-tubulin (CST-5346), GAPDH (CST-3683), PARP1 (CST-9542) (Cell Signaling, Boston, MA) were used. Streptavidin conjugated to horse radish peroxidase was also used (CST-3999).

Quantitative-PCR: For quantitative PCR, we isolated total RNA from cells with Trizol (ThermoFisher Scientific; Waltham, MA) and purified with a commercially available kit (QIAgen; Germantown, MD). Subsequently prepared cDNA (GE First-Strand cDNA Synthesis Kit; Niskayuna, NY) was used as the template for quantitative PCR using the BIORAD iQ5 thermal cycler. Primer sets for human Mapkapk2 (#PPH01783A) and β-Actin (#PPH00073G) were purchased from QIAgen. Background subtracted amplification data were analyzed using open-source software(8) to estimate Ct values and amplification efficiency. Target gene expression was normalized to a reference gene using the comparative Ct method(9).

Nuclear and cytosolic fractionation: Cells were trypsinized and re-suspended in PBS. Nuclear and cytosolic fractions of the resulting cell suspensions were generated using NE-PER Nuclear and Cytoplasmic Extraction Reagents (Thermo Fisher 78833, Rockford, IL). The purity of cytosolic fractions was assessed by lack of PARP1 immuno-reactivity and purity of the nuclear fraction was assessed by lack of GAPDH immuno-reactivity (Cell Signaling, Boston, MA) using standard immuno-blotting techniques.

Brightfield microscopy: Six hours after sub-confluent plating, H23 cells were infected with Ad-eGFP or Ad-WT MK2 and imaged over time. A mark was placed under side each dish to allow for sequential imaging of the same region. All images were anonymized and number of cells per image were counted by blinded investigators (AS, GS, MK). Cell counts obtained by each investigator were subsequently averaged together.

In vitro kinase assay: Reactions were performed in 50 mM sodium β-glycerophosphate containing 10 mM Mg-acetate, 0.1 mM ATP, 2 uCi $\gamma$-$^{32}$P-ATP (New England Nuclear, PerkinElmer, Waltham, MA) and 170 ng (6 U) of recombinant active MK2 (Millipore, Dundee, UK). Substrates were recombinant active caspase-3 (250 ng, Enzo, Farmingdale, NY) or recombinant HSP27 (14 ng, Enzo, Farmingdale, NY). Reactions were incubated at 30ºC for 30 or 60 min, and resolved on 4-12% NUPAGE gels (Life Technologies, Carlsbad, CA). After staining with Coomassie Blue, gels were dried and phosphorylated proteins were detected on a Molecular Imager FX phosphorimager (BioRad, Hercules, CA) using Quantity One software.

Bio-ID assay: H23 cells were infected with Ad-WT MK2, Ad-WT-MK2-BioID-C or Ad-WT-MK2-BioID-N for 24 hours after which the media was changed to include biotin (50 uM) (ThermoFisher, 29129) for an additional 18 hours, after which cell lysates were prepared. Cell lysates were incubated with 50 uL of Streptavidin Sepharose High Performance Beads (GE Healthcare, 17511301) overnight at 4˚C with gentle shaking. Beads were washed four times with buffer to elute off non-specific proteins bound to the sepharose beads. The buffer was aspirated and beads were re-suspended in Laemmli buffer for Western blotting.

### TCGA Data access and analysis

*Clinical and mRNA expression accession:* Clinical data for TCGA datasets was accessed using the *TCGABiolinks* package (11-13) in R/Bioconductor (14, 15). mRNA expression data and censored survival data was obtained using (16). Patient level clinical and mRNA data were merged using individual TCGA sample identifiers. The code used to analyze the TCGA data is presented in markdown format in **Supplemental Data**. All raw data and code pertaining to TCGA analyses performed herein are available at: github.com/suresh-lab/TCGA-casp3/

*Model construction and testing*: Cox proportional hazards models were constructed using the *survminer* (REF XXX) and *survival* packages in R (19). Proportional hazard assumption testing was performed by examining Schoenfield residuals and Q-Q plots. Logistic regression model testing was performed using the Hosmer-Lemeshow test. During initial exploration of the data, we observed that patients were entered into the TCGA dataset over a long period of time (1991-2013). The number of patients enrolled (per year) was skewed, with the majority of patients being enrolled after 2005 (**Supplemental Figure 2A**). To determine whether time of enrollment could be confounding our results, we performed a jackknife analysis whereby we examined the effect of sequential exclusion of one particular year of data (i.e. excluding all patients diagnosed in that year) on the point estimate results for Model 1 2005 (**Supplemental Figure 2B**). We observed that the MK2 hazard ratio and confidence intervals were very stable even with specific years were excluded, suggesting that imbalances in enrollment time were likely not playing a big role. Some survival models (e.g. modeling survival across the entire span of survival time in the validation cohort) failed to meet proportionality hazard assumption testing. Thus, these models were not used.

*Model results:* For survival analysis, Kaplan-Meier curves and Hazard ratios were calculated. For logistic regression, odds ratios as well as predicted probabilities were calculated. Point estimates are provided with 95% confidence intervals. A $P$ value < 0.05 was accepted as statistically significant.

19. Terry M. Therneau PMG. *Modeling Survival Data: Extending the Cox Model.* New York, NY: Springer; 2000.

**Supplemental Figure Legends:**

**Supplemental Figure 1:** Scatterplot comparing MK2 transcript levels obtained via two different methods: a) download of data using the OncoLnc web interface and b) extraction of MK2 transcript reads from the TCGA directly, via GDC.

**Supplemental Figure 2: A.** Bar graph showing histogram of number of patients recruited per year in the TCGA-LUAD dataset. **B.** Jackknife analysis showing point estimate and 95% CI for odds ratio for high MK2 transcript level and death at one year following removal (with replacement) of patients recruited at one particular year within the LUAD dataset.

**Supplemental Figure 3:** Cell lysates from the NSCLC cell lines, H23, A549 and the SCLC cell line, H446 were immunoblotted with antibodies recognizing total MK2. As shown, there is markedly reduced MK2 expression in H23 and A549 cells compared to H446 cells.

**Supplemental Figure 4: A.** H23 NSCLC cells were infected with adenovirus encoding MK2 or eGFP (Ad-MK2 and Ad-eGFP, respectively) for 48hrs and then harvested for protein analyses. H23 cells infected with Ad-MK2 leads to marked increase in MK2 protein expression. **B.** Gating strategy to identify apoptotic H23 cells. Forward and side scatter (FSC and SSC) gating on area (A), height (H), and width (W) excluded debris and non-single cell events. Then relative fluorescence of DAPI staining was measured. The amount of positive DAPI fluorescent staining was quantified.

**Supplemental Figure 5: MK2 mRNA and protein expression is directly correlated in NSCLC.** Non-small cell lung cancer cell lines examined by the Cancer Dependency Map project (www.https://depmap.org) shows direct correlation between MK2 mRNA and MK2 protein expression. Solid square- A549. Solid triangle- H23.

**Supplemental Figure 6:** Scatterplots with linear regression line showing proportion of people who died within each quantile of MK2 expression in early- and late-stage NSCLC.

**Supplemental Figure 7:** H23 cells were exposed to etoposide at varying concentrations for 24hrs followed by staining with DAPI and cell death was assessed using flow cytometry. There is a dose dependent increase in cell death in response to etoposide with 96ug/mL resulting in the highest amount of death. *, $P < 0.05$ vs all other conditions using Dunnett's multiple comparison test.

**Supplemental Figure 8:** Lung homogenates from wild-type and $MK^{-/-}$ mice exposed to PBS or LPS (IV 7.5mg/kg, 6hrs) were treated with or without the serine/threonine phosphatase PP2A and underwent 2-Dimensional immunoblotting for caspase 3. As shown, there is a shift toward the positive electrode on an isoelectric gradient in lung homogenates from WT animals exposed to LPS (Red Dash Arrow). Treatment of lung homogenates from WT animals exposed to LPS with active recombinant serine/threonine phosphatase, PP2A, reverses these charge based shifts (red arrow), suggestive of a LPS-induced phosphorylation event. Additionally, 2-Dimensional immunoblotting for caspase 3 of lung homogenates from *MK$^{-/-}$* mice exposed to LPS appears similar to that of lung homogenates from WT animals exposed to LPS and treated with active recombinant PP2A, suggestive of a MK2-dependent phosphorylation event on caspase 3. Representative immunoblots from 3 mice/group.

**Supplemental Figure 9:** H23 NSCLC cells were infected with Ad-eGFP, Ad-WT-MK2, Ad-Active-MK2 or Ad-Dom Neg-MK2 for 48hrs and then harvested for protein analyses. Representative indicates an increase in cleavage of PARP1 in Ad-WT MK2, Ad-Active-MK2 and Ad-Dom Neg-MK2 infected H23 cells compared to Ad-eGFP infected H23 cells.

**Supplemental Figure 10:** H23 cells were infected with Ad-WT-MK2, Ad-WT MK2-BioID-C (MK2 fused to biotin ligase on the c-terminus) or Ad-WT MK2-BioID-N (MK2 fused to biotin ligase on the n-terminus) and following incubation with biotin, cell lysates were precipitated with beads conjugated to streptavidin and then subjected to immunoblotting. **A:** There is an increase in molecular weight of MK2 when conjugated to biotin ligase, as shown by the shift in the immunoreactive band. There is also phosphorylation of HSP27 in cells infected with Ad-WT-MK2, Ad-WT MK2-BioID-C or Ad-WT MK2-BioID-N, demonstrating the addition of the biotin ligase did not impact its kinase function. **B.** Purity of streptavidin pull downs is demonstrated by minimal immunoreactivity in the supernatants and in lysates infected with Ad-WT-MK2. **C.** Representative immunoblot of lysates from H23 cells infected with Ad-eGFP, Ad-WT MK2 or Ad-Mut-NLS-MK2 for 48 hours demonstrates phosphorylation of HSP27 in cells infected with Ad-WT MK2 or Ad-

Mut-NLS-MK2, demonstrating retention of MK2's kinase activity with the mutation within the NLS.

**Supplemental Figure 11:** H23 cells were infected with Ad-eGFP or Ad-WT MK2 for 24 hours and then treated with etoposide 96ug/mL for 24hrs. Cells were then analyzed for cell death using flow cytometry. As shown, there is significant increase in cell death with Ad-WT MK2. *, $P < 0.05$ vs Ad-eGFP.
