## Supplemental Data for "MK2 Expression Promotes Non-Small Cell Lung Cancer Cell Death and Predicts Survival"

### Analysis of MK2 transcript levels in TCGA-LUAD (NSCLC) dataset

Karthik Suresh

November 18, 2021 - Revision 4

**Additional models and analysis** 44
Continuous MK2 expression, no R censoring: Survival ~ expression (continuous) + clinical stage (stage 1, stage 2,. . . ) . . . . . . . . . . . . . . . . . . . . . . . . . . . . . . . . . . . . . . . . 44
Logistic regression for death at 1 year, stratified by specific stage . . . . . . . . . . . . . . . . . . 46
Jackknife-type analysis for effects due to long period of patient recruitment . . . . . . . . . . . . . 52

**Hsp27 and MK2** 57

Revisions (since rev2):

1. We've added the concept of validating RNASeq counts using a secondary extraction process. Using TCGAbiolinks, we extracted normalized RSEM reads and compared these reads to those obtained via OncoLnc to ensure that the read counts were largely similar at a patient-to-patient level

2. The original scatter plot showing differences in MK2 is restricted to only "Adenocarcinomas" within LUAD. We've applied that restriction to all the regression analyses as well
3. Some small clean-ups to the code
4. We've re-analyzed the proportion dead by tertiles of MK2 (instead of quantiles)
5. One advantage of downloading TCGA data is that we also happened to grab MK2 transcript levels from 50 samples with normal tissue controls, thus allowing us to calculate fold change in MK2 transcript levels between normal and tumor in matched samples.
6. R censoring is now performed at one year and two year. This is in part to harmonize analyses with the validation cohort, where censoring could only be done at 2 years.

### Data retrieval and cleaning

First, we are downloading clinical data from TCGA-LUAD dataset using the TCGAbiolinks API. For the MK2 transcript data, I grabbed these data from oncoLnc.org using "MAPKAPK2" as the gene name and TCGA-LUAD as the cohort.

THe code used to generate the TCGA manual reads is provided elsewhere.

#### Retrieve data from TCGA dataset, __join it with the MK2 transcript data, with additional joins for the manually extracted reads from TCGA and for Casp3 (from OncoLnc)

We retrieve the clinical data directly using GDC_query. For the MK2 transcript data, this was downloaded from OncoLnc.org using the following parameters: Gene-MAPKAPK2, and percentile: 50/50. We did this because this allowed us to get transcript levels on all available patients (since upper and lower 50% would include all patients). We then calculated the quantiles manually, so that in the future, if we needed to change the threshold, we could do so without re-downloading the OncoLnc data.

In this version, there is an additional step. Using TCGAbiolinks() we have manually extracted normalized RSEM reads from the LUAD dataset and compared it to what was given to us by OncoLnc

```
library(knitr)
opts_chunk$set(tidy.opts=list(width.cutoff=70), tidy=TRUE)
```

```
knitr::opts_chunk$set(echo = TRUE)
rm(list = ls())
library("SummarizedExperiment")
```

```
## Loading required package: MatrixGenerics

## Loading required package: matrixStats

##
## Attaching package: 'MatrixGenerics'

## The following objects are masked from 'package:matrixStats':
##
##     colAlls, colAnyNAs, colAnys, colAvgsPerRowSet, colCollapse,
##     colCounts, colCummaxs, colCummins, colCumprods, colCumsums,
##     colDiffs, colIQRDiffs, colIQRs, colLogSumExps, colMadDiffs,
##     colMads, colMaxs, colMeans2, colMedians, colMins, colOrderStats,
##     colProds, colQuantiles, colRanges, colRanks, colSdDiffs, colSds,
##     colSums2, colTabulates, colVarDiffs, colVars, colWeightedMads,
##     colWeightedMeans, colWeightedMedians, colWeightedSds,
##     colWeightedVars, rowAlls, rowAnyNAs, rowAnys, rowAvgsPerColSet,
```

```
##     rowCollapse, rowCounts, rowCummaxs, rowCummins, rowCumprods,
##     rowCumsums, rowDiffs, rowIQRDiffs, rowIQRs, rowLogSumExps,
##     rowMadDiffs, rowMads, rowMaxs, rowMeans2, rowMedians, rowMins,
##     rowOrderStats, rowProds, rowQuantiles, rowRanges, rowRanks,
##     rowSdDiffs, rowSds, rowSums2, rowTabulates, rowVarDiffs, rowVars,
##     rowWeightedMads, rowWeightedMeans, rowWeightedMedians,
##     rowWeightedSds, rowWeightedVars

## Loading required package: GenomicRanges

## Loading required package: stats4

## Loading required package: BiocGenerics

## Loading required package: parallel

##
## Attaching package: 'BiocGenerics'

## The following objects are masked from 'package:parallel':
##
##     clusterApply, clusterApplyLB, clusterCall, clusterEvalQ,
##     clusterExport, clusterMap, parApply, parCapply, parLapply,
##     parLapplyLB, parRapply, parSapply, parSapplyLB

## The following objects are masked from 'package:stats':
##
##     IQR, mad, sd, var, xtabs

## The following objects are masked from 'package:base':
##
##     anyDuplicated, append, as.data.frame, basename, cbind, colnames,
##     dirname, do.call, duplicated, eval, evalq, Filter, Find, get, grep,
##     grepl, intersect, is.unsorted, lapply, Map, mapply, match, mget,
##     order, paste, pmax, pmax.int, pmin, pmin.int, Position, rank,
##     rbind, Reduce, rownames, sapply, setdiff, sort, table, tapply,
##     union, unique, unsplit, which.max, which.min

## Loading required package: S4Vectors

##
## Attaching package: 'S4Vectors'

## The following objects are masked from 'package:base':
##
##     expand.grid, I, unname

## Loading required package: IRanges

## Loading required package: GenomeInfoDb

## Loading required package: Biobase

## Welcome to Bioconductor
##
##     Vignettes contain introductory material; view with
##     'browseVignettes()'. To cite Bioconductor, see
##     'citation("Biobase")', and for packages 'citation("pkgname")'.

##
## Attaching package: 'Biobase'
```

```
## The following object is masked from 'package:MatrixGenerics':
##
##     rowMedians

## The following objects are masked from 'package:matrixStats':
##
##     anyMissing, rowMedians
```

```r
library("dplyr")
```

```
##
## Attaching package: 'dplyr'

## The following object is masked from 'package:Biobase':
##
##     combine

## The following objects are masked from 'package:GenomicRanges':
##
##     intersect, setdiff, union

## The following object is masked from 'package:GenomeInfoDb':
##
##     intersect

## The following objects are masked from 'package:IRanges':
##
##     collapse, desc, intersect, setdiff, slice, union

## The following objects are masked from 'package:S4Vectors':
##
##     first, intersect, rename, setdiff, setequal, union

## The following objects are masked from 'package:BiocGenerics':
##
##     combine, intersect, setdiff, union

## The following object is masked from 'package:matrixStats':
##
##     count

## The following objects are masked from 'package:stats':
##
##     filter, lag

## The following objects are masked from 'package:base':
##
##     intersect, setdiff, setequal, union
```

```r
library("DT")
library("TCGAbiolinks")
library(survival)
library(ggplot2)
library(ggpubr)
library(survminer)
library(knitr)
library("tidyr")
```

```
##
## Attaching package: 'tidyr'
```

```
## The following object is masked from 'package:S4Vectors':
##
##     expand
```

```
library("ggpmisc")
```

```
## Loading required package: ggpp
```

```
##
## Attaching package: 'ggpp'
```

```
## The following object is masked from 'package:ggplot2':
##
##     annotate
```

```
clinical <- GDCquery_clinic(project = "TCGA-LUAD", type = "clinical")
expr <- read.csv("../rawdata/LUAD.MAPKAPK2Exp.csv", header = TRUE)
colnames(expr) <- c("Patient", "MK2.Expression")
expr2 <- read.csv("../rawdata/LUAD.MAPKAPK2Exp.manual.csv", header = FALSE)
Casp3exp <- read.csv("../rawdata/LUAD.casp3Exp.2.csv")

colnames(expr2) <- c("MK2.Expression.Manual", "Stage", "Patient")
expr2_normalcontrols <- expr2[grepl("11A", expr2$Patient), ]
# take out normal controls
expr2 <- expr2[!grepl("11A", expr2$Patient), ]

# remove non '01A' samples
expr3 <- expr2[grepl("01A", expr2$Patient), ]
expr3$Patient <- substring(expr3$Patient, 1, 12)
colnames(Casp3exp) <- c("Patient", "Casp3.Expression")
# the left_join. we need the clinical data, so we left_joined by TCGA
# patient id to the clinical dataset. This means that if there was a
# patient with clinical data without transcript levels, that would be
# a 'NA'.
clinical <- left_join(clinical, expr, c(submitter_id = "Patient"))
clinical <- left_join(clinical, expr3, c(submitter_id = "Patient"))
clinical <- left_join(clinical, Casp3exp, c(submitter_id = "Patient"))

ggplot(clinical, aes(x = MK2.Expression, y = MK2.Expression.Manual)) +
    geom_point() + theme_bw() + ylab("MAPKAPK2 transcript level - direct extract from LUAD") +
    xlab("MAPKAPK2 transcript level - OncoLnc")
```

```
## Warning: Removed 104 rows containing missing values (geom_point).
```

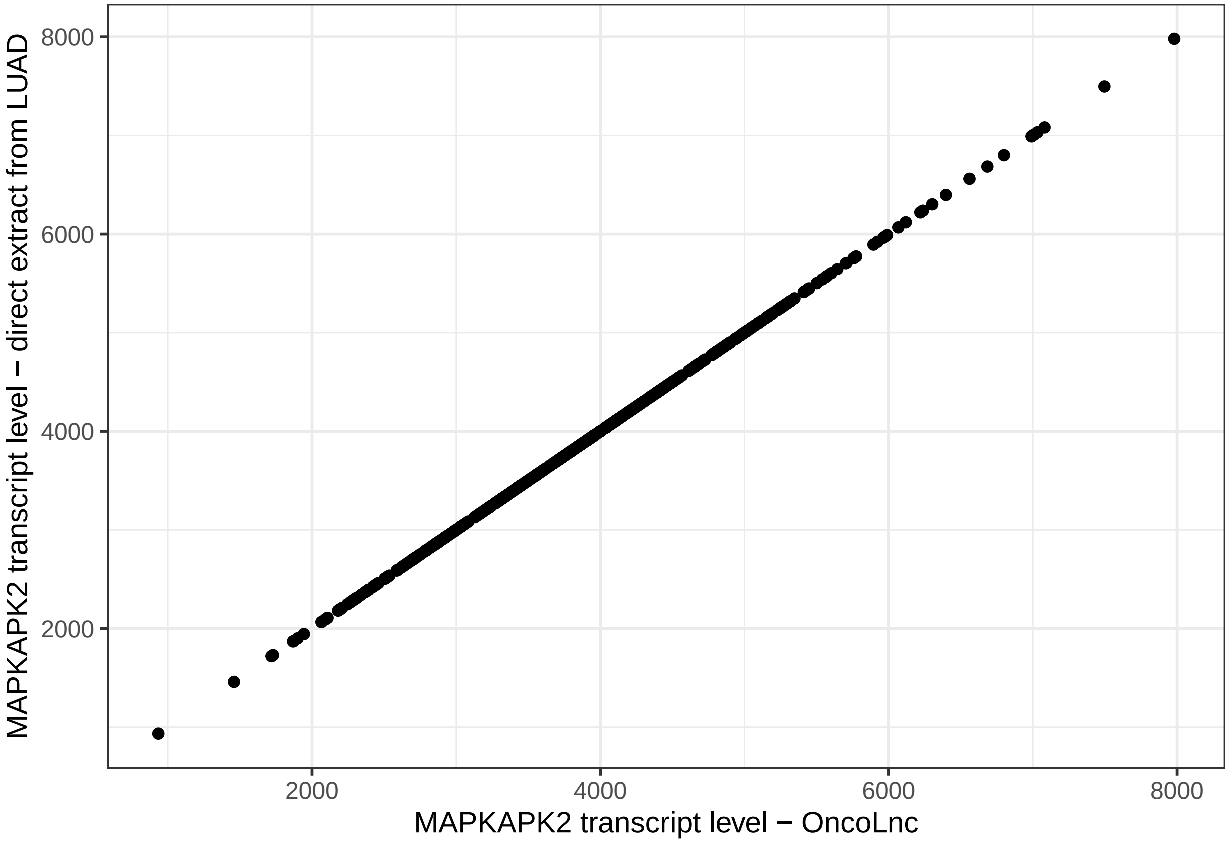

```
clinical <- clinical[!is.na(clinical$vital_status), ]
```

This shows that the correlation between MK2 Expression derived from OncoLnc and from TCGA is excellent. THus, we feel confident that the MK2 reads data from OncoLnc can be used going forward. This makes it easier to grab mRNA transcript levels for other genes directly from OncoLnc rather than having to extract it from TCGA via GDC.

#### Clean the data, create a df for an LR and Cox PH analyses later

Now that we have left-joined the data, and checked to make sure that the join was done properly, we move on to prep the data for Cox PH work.

```
clinical$time <- mapply(function(x, y) if (is.na(x)) return(y) else return(x),
    clinical$days_to_death, clinical$days_to_last_follow_up)
clinical$age_at_diagnosis <- clinical$age_at_diagnosis/365
clinical$race <- as.factor(clinical$race)
clinical$smoking <- lapply(clinical$cigarettes_per_day, function(x) if (!is.na(x)) return(1) else retur
clinical$tumor_grade <- as.factor(clinical$tumor_grade)
clinical$vital_status <- as.factor(clinical$vital_status)
clinical$status <- lapply(clinical$vital_status, function(x) if (x == "alive") return(0) else return(1)
clinical$smoking <- as.numeric(clinical$smoking)

# combine clinical stage

combineStage <- function(x) {
    if (!is.na(x)) {
        if (x == "Stage I" | x == "Stage IB" | x == "Stage IA")
```

```
            return("Early Stage")
        if (x == "Stage II" | x == "Stage IIA" | x == "Stage IIB")
            return("Early Stage")
        if (x == "Stage IIIA" | x == "Stage IIIB")
            return("Late Stage")
        if (x == "Stage IV")
            return("Late Stage")
    }
}

# combined histologic types

combinePath <- function(x) {
    if (!is.na(x)) {

        if (x == "Adenocarcinoma, NOS")
            return("Adenocarcinoma")
        if (x == "Mucinous adenocarcinoma")
            return("Adenocarcinoma")
        if (x == "Adenocarcinoma with mixed subtypes")
            return("Adenocarcinoma")
        if (x == "Papillary adenocarcinoma")
            return("Adenocarcinoma") else return("Non-Adenocarcinoma")

    }

}

# combine sub-stages into 1,2,3
clinical$ajcc_pathologic_stage_combined <- lapply(clinical$ajcc_pathologic_stage,
    combineStage)

clinical$ajcc_pathologic_stage_combined <- as.factor(as.character(clinical$ajcc_pathologic_stage_combin

# combine histology types
clinical$primary_diagnosis_combined <- lapply(clinical$primary_diagnosis,
    combinePath)
clinical$primary_diagnosis_combined <- as.factor(unlist(clinical$primary_diagnosis_combined))

# get rid of NULL entries
clinical <- clinical[clinical$ajcc_pathologic_stage_combined != "NULL",
    ]

# We already got rid of NULLs for ajcc_pathologic_stage. However, the
# factor levels still list 3 levels - early, late and NULL. To
# re-level, convert to character, re-convert to factor, and now only
# 2 levels.
clinical$ajcc_pathologic_stage_combined <- as.factor(as.character(clinical$ajcc_pathologic_stage_combin

clinical <- clinical[!is.na(clinical$MK2.Expression), ]

# create censor variable for CoxPH
```

```r
clinical$censor <- lapply(clinical$vital_status, function(x) if (x == "Alive") return(0) else return(1))
clinical$censor <- as.numeric(clinical$censor)

# scale the MK2 variable
clinical$MK2.Expression <- clinical$MK2.Expression/1000
clinical$MK2.Expression.Manual <- clinical$MK2.Expression.Manual/1000
clinical$Casp3.Expression <- clinical$Casp3.Expression/1000
# create a simplified stage

combineStage.Simp <- function(x) {
    if (!is.na(x)) {
        if (x == "Stage I" | x == "Stage IB" | x == "Stage IA")
            return("Stage 1")
        if (x == "Stage II" | x == "Stage IIA" | x == "Stage IIB")
            return("Stage 2")
        if (x == "Stage IIIA" | x == "Stage IIIB")
            return("Stage 3")
        if (x == "Stage IV")
            return("Stage 4")
    }
}
clinical$ajcc_pathologic_stage_combined.2 <- lapply(clinical$ajcc_pathologic_stage,
    combineStage.Simp)
clinical$ajcc_pathologic_stage_combined.2 <- as.factor(unlist(clinical$ajcc_pathologic_stage_combined.2))

write.table(as.matrix(clinical), file = "../rawdata/tcga-luad-clinical.processed.csv",
    sep = ",", quote = FALSE, col.names = TRUE)
```

### Exploratory graphs

#### Plot differences in transcript level by tumor histology and cancer stage

```r
# plot MK2 differences for adenocarcinoma only
ggplot(clinical[clinical$primary_diagnosis_combined == "Adenocarcinoma",
    ], aes(x = ajcc_pathologic_stage_combined, y = MK2.Expression, )) +
    geom_boxplot() + geom_point(position = position_jitter()) + stat_compare_means() +
    theme_bw() + ylab("MK2 transcript level")
```

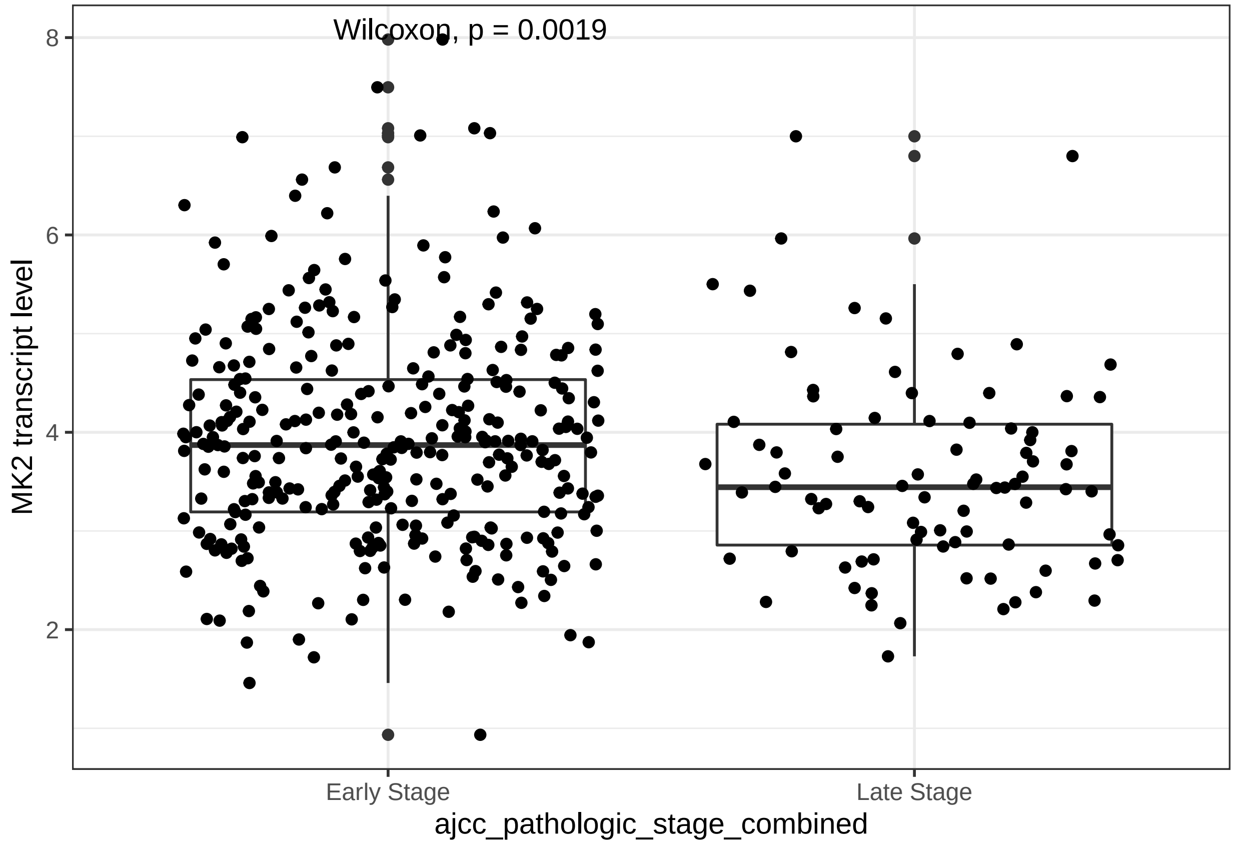

```
ggplot(clinical[clinical$primary_diagnosis_combined == "Adenocarcinoma",
    ], aes(x = ajcc_pathologic_stage_combined, y = MK2.Expression.Manual,
    )) + geom_boxplot() + geom_point(position = position_jitter()) + stat_compare_means() +
    theme_bw() + ylab("MK2 transcript level")
```

```
## Warning: Removed 1 rows containing non-finite values (stat_boxplot).

## Warning: Removed 1 rows containing non-finite values (stat_compare_means).

## Warning: Removed 1 rows containing missing values (geom_point).
```

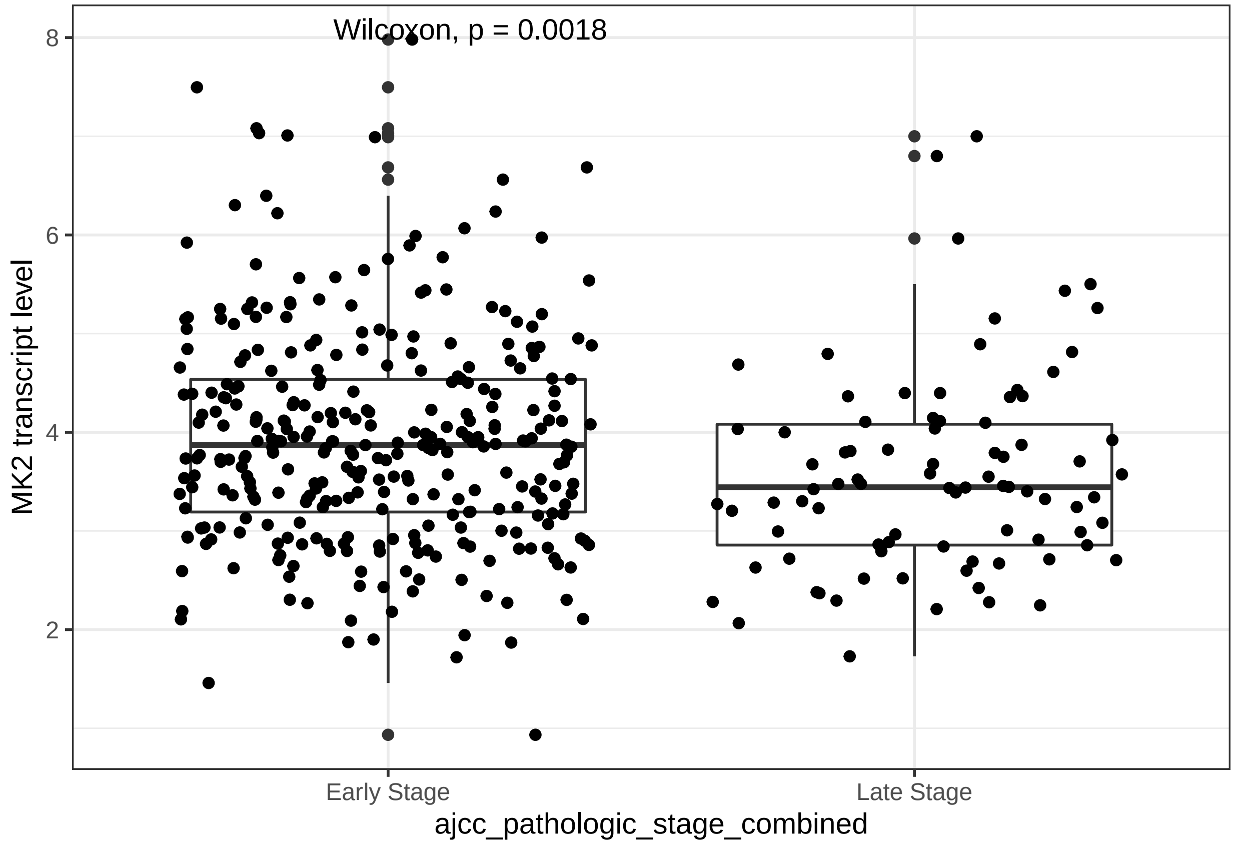

This is a key difference: MK2 is clearly associated with tumor stage. Early stage tumors have more MK2 transcript than late stage tumors. Thus, going forward, we will stratify our analyses by Tumor stage.

Here, we look at tumor differences more granularly (stage 1 vs. 2 vs. 3 vs. 4) and see a linear relationship - as stage increases, MK2 exp decreases. We also see that the LUAD cases that aren't really "AD" (non-Adenocarcinoma) don't share this relationship between Early/Late stage and MK2 transcript levels. So, when we prep the data for model work (below) we will restrict to Adenocarcinoma cases only

```
ggplot(clinical[clinical$primary_diagnosis_combined == "Adenocarcinoma",
    ], aes(x = ajcc_pathologic_stage_combined.2, y = MK2.Expression, )) +
    geom_boxplot() + geom_point(position = position_jitter()) + stat_compare_means() +
    theme_bw() + ylab("MK2 transcript level")
```

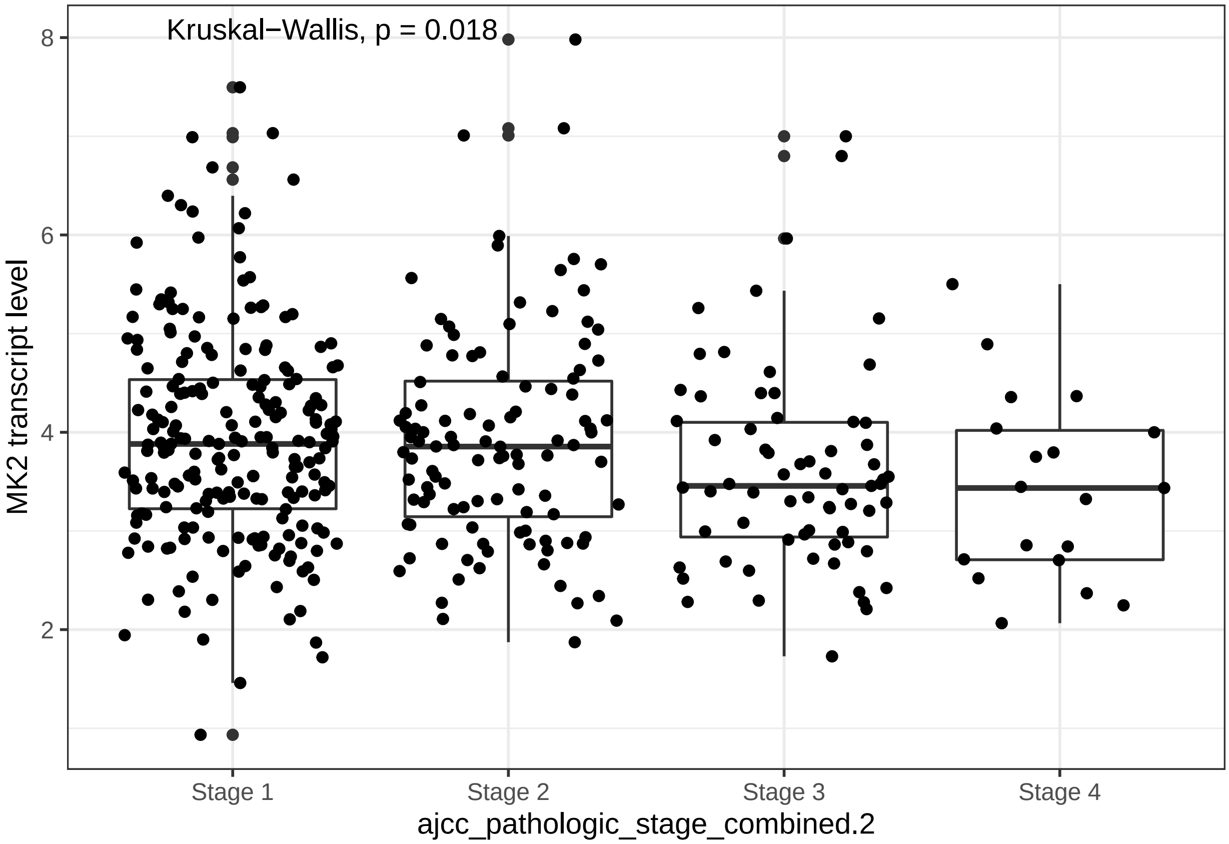

```
ggplot(clinical, aes(x = ajcc_pathologic_stage_combined, y = MK2.Expression,
    fill = primary_diagnosis_combined)) + geom_boxplot() + geom_point(position = position_jitterdodge()
    theme_bw()
```

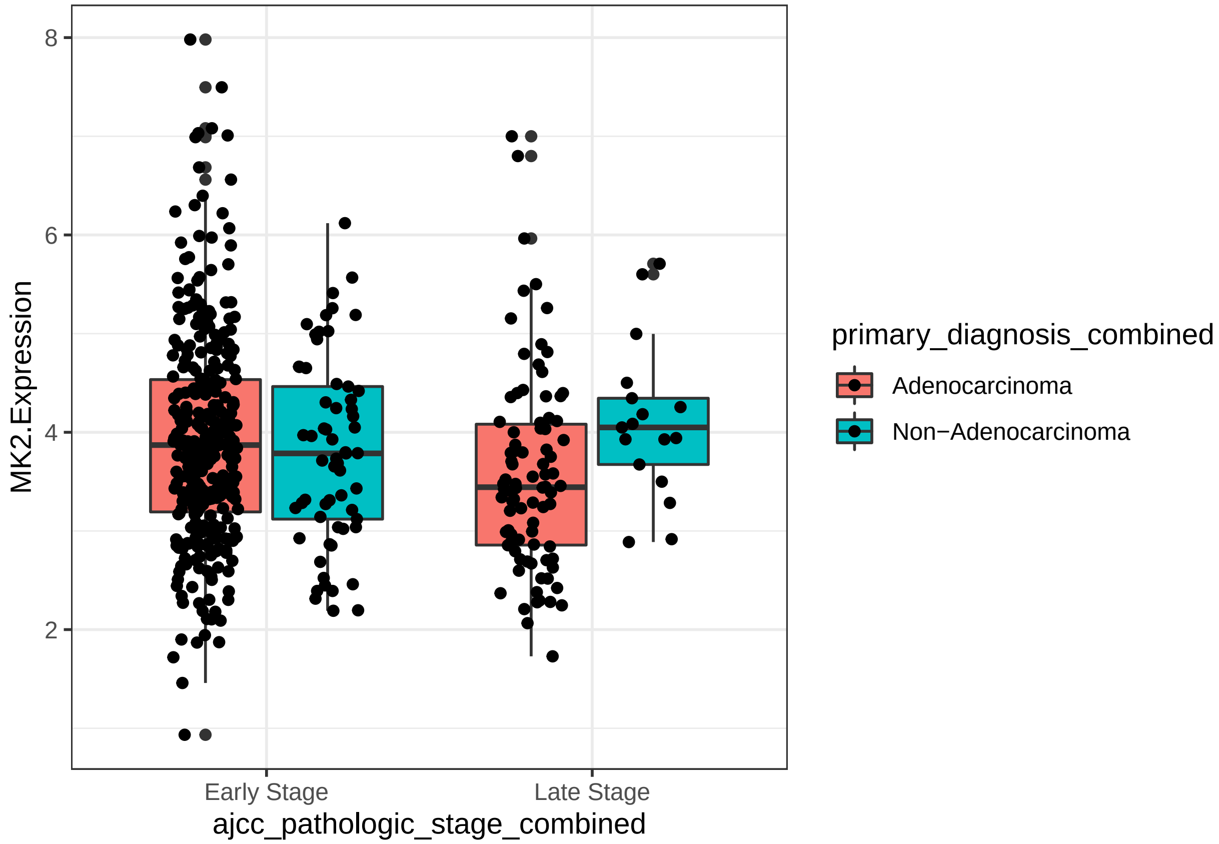

##### Explore the relationship between MK2 and Casp3

Here we look at the correlation between MK2 and Casp3 transcript levels.

```
ggplot(clinical, aes(x = MK2.Expression, y = Casp3.Expression)) + geom_point() +
    theme_bw() + geom_smooth(method = "lm")
```

```
## `geom_smooth()` using formula 'y ~ x'
```

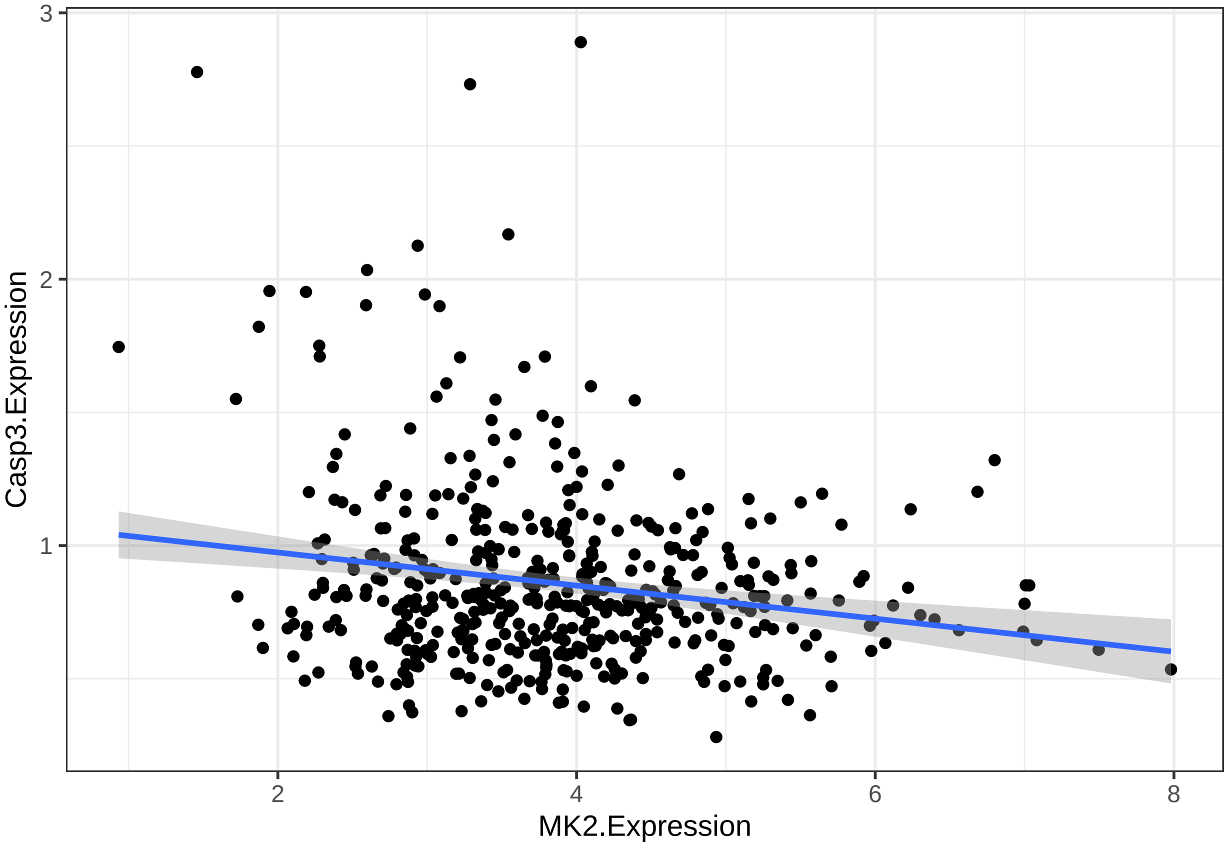

```
ggplot(clinical, aes(x = MK2.Expression.Manual, y = Casp3.Expression)) +
    geom_point() + theme_bw()
```

```
## Warning: Removed 2 rows containing missing values (geom_point).
```

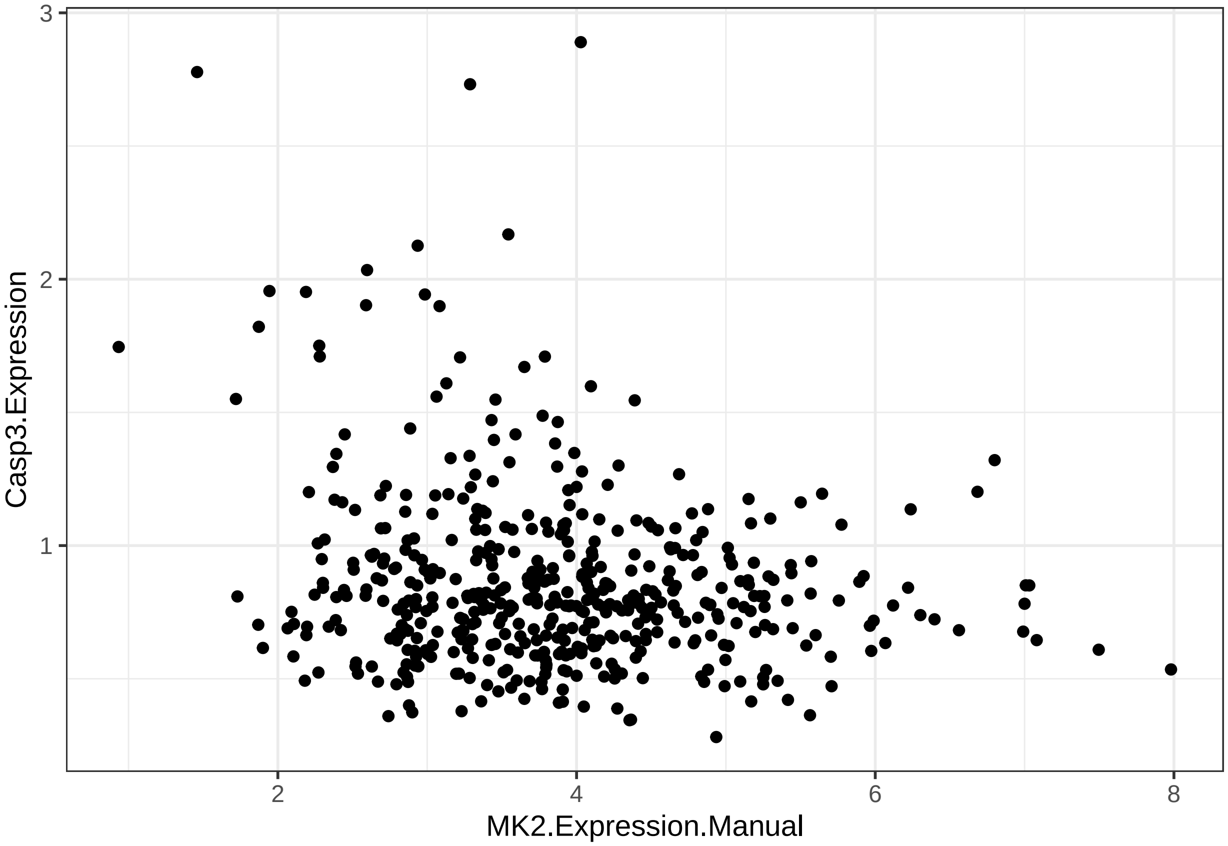

We are technically able to "fit" a line through this data, but the relationship (i.e slope) is not particularly impressive. The raw data does not suggest any sort of linear relationship between Casp3 mRNA and MK2 mRNA levels.

##### Defining MK2 thresholds for "high" and "low" - Top 1/3 vs Bottom 2/3

Mechanistically, we're more interested in whether high vs. low MK2 transcript levels impact outcome. For this, we first categorize MK2 transcript levels based on quantiles and only take the lower and upper third in our groups for comparison. The MK2_Expression_topQ variable essentially captures this. If the transcript is in the top 33% then we return 1, everthing else (i.e the bottom 66%) returns 0.

```
# classify the MK2 expression variable as either top or bottom
# quantile, based on quantile definitions

bottom_quant <<- 0.33
top_quant <<- 0.66

clinical$MK2_Expression_bottomQ <- lapply(clinical$MK2.Expression, function(x) if (x <
    quantile(clinical$MK2.Expression, bottom_quant)) return(1) else return(0))

clinical$MK2_Expression_topQ <- lapply(clinical$MK2.Expression, function(x) if (x >
    quantile(clinical$MK2.Expression, top_quant)) return(1) else return(0))
```

#### Changes in proportion of people who died across quantiles of MK2

##### Prep data for using MK2 quantiles

The (poorly named) MK2_CoxPH_quantile df is the df we're using for all of our modeling purposes. This was originally used just for CoxPH, hence the name. We restrict this dataset to only those with follow-up available, and Adenocarcinomas only within LUAD. Additionally, we perform some more cleanup/create new variables as shown below.

```
# quantile and logistic regression

# this is the new data frame for this new round of calculations.
MK2_CoxPH_quantile <- clinical[!is.na(clinical$time) & clinical$primary_diagnosis_combined ==
    "Adenocarcinoma", ]

# this provides the variable for R censoring at one year and the
# outcome variable for logit regression.
MK2_CoxPH_quantile$death_at_1year <- mapply(function(dead, time) if (dead ==
    1 & time < 366) return(1) else if (dead == 0 & time > 366) return(0) else if (dead ==
    0 & time < 366) return(0) else if (dead == 1 & time > 366) return(0),
    MK2_CoxPH_quantile$censor, as.numeric(MK2_CoxPH_quantile$time))

MK2_CoxPH_quantile$death_at_2year <- mapply(function(dead, time) if (dead ==
    1 & time < 366 * 2) return(1) else if (dead == 0 & time > 366 * 2) return(0) else if (dead ==
    0 & time < 366 * 2) return(0) else if (dead == 1 & time > 366 * 2) return(0),
    MK2_CoxPH_quantile$censor, as.numeric(MK2_CoxPH_quantile$time))

MK2_CoxPH_quantile$death_at_3year <- mapply(function(dead, time) if (dead ==
    1 & time < 366 * 3) return(1) else if (dead == 0 & time > 366 * 3) return(0) else if (dead ==
    0 & time < 366 * 3) return(0) else if (dead == 1 & time > 366 * 3) return(0),
    MK2_CoxPH_quantile$censor, as.numeric(MK2_CoxPH_quantile$time))

# this piece of code above is critical - so let's double check and
# make sure everything got done right

ggplot(MK2_CoxPH_quantile, aes(x = submitter_id, y = time, color = vital_status)) +
    geom_point() + facet_wrap(~death_at_1year) + ylim(0, 500) + xlab("Patient ID") +
    ylab("time to death") + geom_hline(yintercept = 365)
```

```
## Warning: Removed 258 rows containing missing values (geom_point).
```

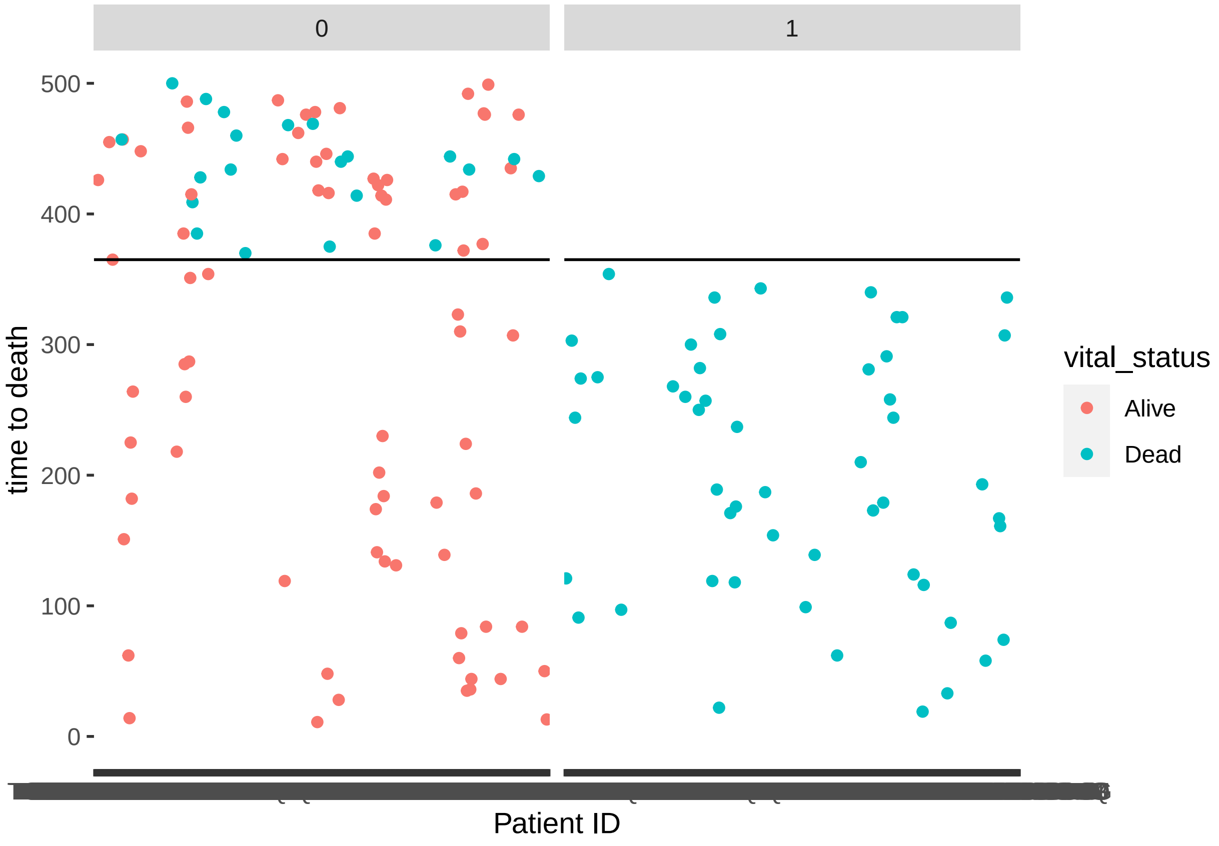

Interpretation off the plot above. So, we have colored each patient's vital status by alive or dead, and we have faceted by whether they were assigned as "Death in one year" or not. So, if our code worked, all patients who died in less than one year (blue color; y < 365) should be classified as dead at one year ("1"). All patients who were alive in that time period should be classified as red and in the "0" column. After 365 days, everyone should have gotten a classification of "0" for death at one year, since they are all by definition still alive at 1 year, regardless of subsequent alive or dead status (color should mix, all colors in the "0" column). And that's what we see. . .

```
# lastly, going forward, we wish to show time in months rather than
# days, so we'll divide by 30
MK2_CoxPH_quantile$time <- MK2_CoxPH_quantile$time/30
```

##### Proportion of deaths by MK2 quantiles

```
# Now, moving on.

MK2_CoxPH_quantile$MK2_quantile <- ntile(MK2_CoxPH_quantile$MK2.Expression,
    3)
MK2_CoxPH_quantile$Casp3_quantile <- ntile(MK2_CoxPH_quantile$Casp3.Expression,
    3)
a <- aggregate(death_at_1year ~ MK2_quantile + ajcc_pathologic_stage_combined,
    FUN = sum, data = MK2_CoxPH_quantile)
b <- aggregate(death_at_1year ~ MK2_quantile + ajcc_pathologic_stage_combined,
    FUN = length, data = MK2_CoxPH_quantile)

quantile_df <- as.data.frame(a$MK2_quantile)
quantile_df$stage <- as.data.frame(a$ajcc_pathologic_stage_combined)
```

```
quantile_df$prop <- as.data.frame(unlist(a$death_at_1year/b$death_at_1year))
quantile_df$prop <- unlist(quantile_df$prop)
colnames(quantile_df) <- c("quantile", "stage", "prop")
p1 <- ggplot(quantile_df[quantile_df$stage == "Early Stage", ], aes(x = quantile,
    y = prop)) + geom_point() + geom_smooth(method = "lm", se = FALSE) +
    stat_poly_eq(formula = "y~x", aes(label = paste(..eq.label.., ..rr.label..,
        sep = "~~~")), parse = TRUE) + theme_bw() + ylim(0, 0.5)

p2 <- ggplot(quantile_df[quantile_df$stage == "Late Stage", ], aes(x = quantile,
    y = prop)) + geom_point() + geom_smooth(method = "lm", se = FALSE) +
    stat_poly_eq(formula = "y~x", aes(label = paste(..eq.label.., ..rr.label..,
        sep = "~~~")), parse = TRUE) + theme_bw() + ylim(0, 0.5)

# This graph shows the relationship between MK2 quantile and
# proportion of patients who died in Early and Late Stage cancers.
# The relationship between MK2 quantile and Proportion of those
# patients who died (denominator: all patients in that quantile)
gridExtra::grid.arrange(p1, p2, ncol = 2)
```

```
## `geom_smooth()` using formula 'y ~ x'
## `geom_smooth()` using formula 'y ~ x'
```

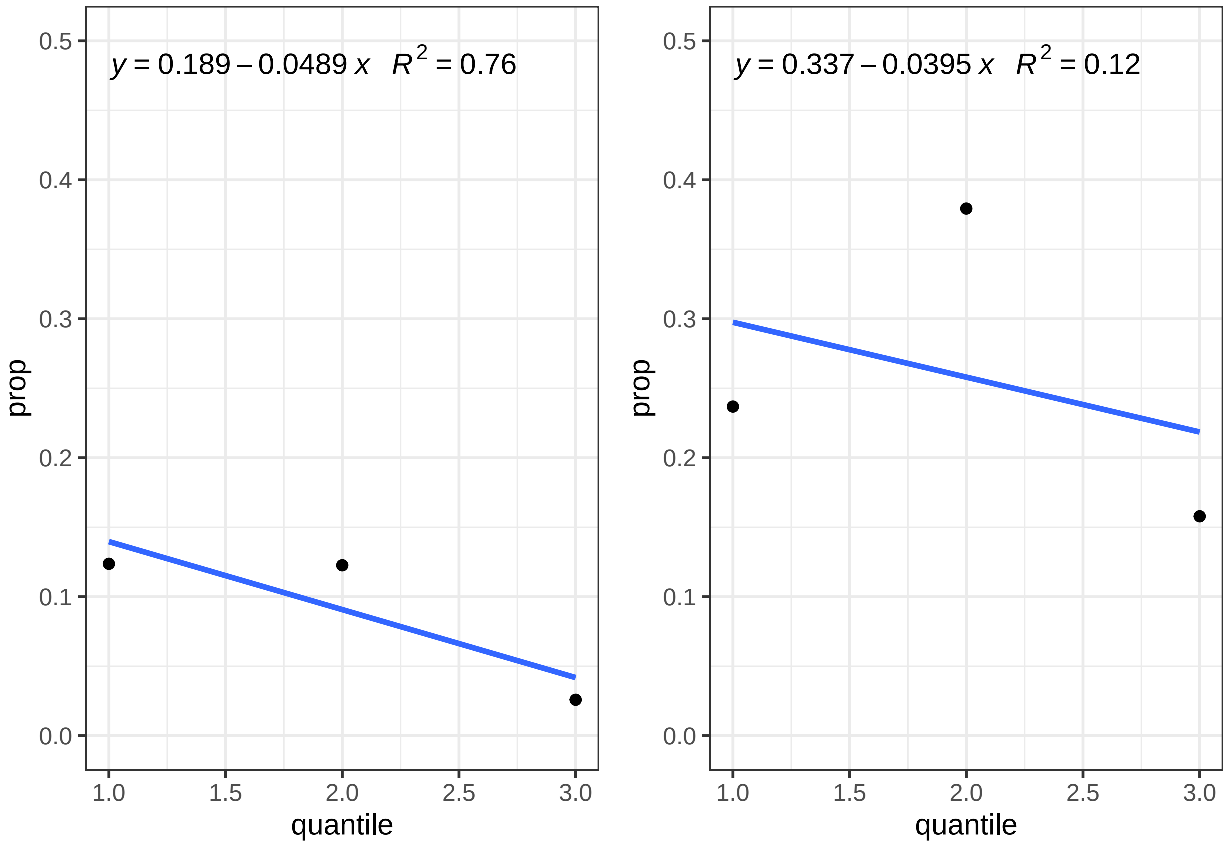

### Logistic regression modeling death at 1 year ~ MK2+covariates

#### Data clean-up

```
MK2_CoxPH_quantile$death_at_1year <- as.factor(MK2_CoxPH_quantile$death_at_1year)
# logistic regression and ROC
MK2_CoxPH_quantile$ajcc_pathologic_stage_early_stage <- unlist(MK2_CoxPH_quantile$ajcc_pathologic_stage_
MK2_CoxPH_quantile$MK2_quantile <- as.factor(MK2_CoxPH_quantile$MK2_quantile)
MK2_CoxPH_quantile$MK2_Expression_topQ <- as.factor(unlist(MK2_CoxPH_quantile$MK2_Expression_topQ))
```

#### Logistic regression for death at 1 year, stratified by early/late, MK2 as a continuous variable

##### Univariate regression

So, here we run a univariate analysis, with MK2 as a continuous variable, stratified by stage. We see a signal in early but not late stage disease.

```
# LR stratified by Early/Late
summary(glm(death_at_1year ~ MK2.Expression, data = MK2_CoxPH_quantile[MK2_CoxPH_quantile$ajcc_patholog
    "Early Stage", ], family = binomial(link = "logit")))
```

```
##
## Call:
## glm(formula = death_at_1year ~ MK2.Expression, family = binomial(link = "logit"),
##     data = MK2_CoxPH_quantile[MK2_CoxPH_quantile$ajcc_pathologic_stage_combined ==
##         "Early Stage", ])
##
## Deviance Residuals:
##     Min       1Q   Median       3Q      Max
## -0.7132  -0.4651  -0.4104  -0.3454   2.4882
##
## Coefficients:
##                Estimate Std. Error z value Pr(>|z|)
## (Intercept)     -0.8740     0.7590  -1.152   0.2495
## MK2.Expression  -0.3910     0.2041  -1.916   0.0554 .
## ---
## Signif. codes:  0 '***' 0.001 '**' 0.01 '*' 0.05 '.' 0.1 ' ' 1
##
## (Dispersion parameter for binomial family taken to be 1)
##
##     Null deviance: 189.71  on 318  degrees of freedom
## Residual deviance: 185.76  on 317  degrees of freedom
## AIC: 189.76
##
## Number of Fisher Scoring iterations: 5
```

```
summary(glm(death_at_1year ~ MK2.Expression, data = MK2_CoxPH_quantile[MK2_CoxPH_quantile$ajcc_patholog
    "Late Stage", ], family = binomial(link = "logit")))
```

```
##
## Call:
## glm(formula = death_at_1year ~ MK2.Expression, family = binomial(link = "logit"),
##     data = MK2_CoxPH_quantile[MK2_CoxPH_quantile$ajcc_pathologic_stage_combined ==
##         "Late Stage", ])
##
```

```
## Deviance Residuals:
##     Min       1Q   Median       3Q      Max
## -0.8449  -0.8022  -0.7821   1.5805   1.6744
##
## Coefficients:
##                Estimate Std. Error z value Pr(>|z|)
## (Intercept)    -0.69350    0.91543  -0.758    0.449
## MK2.Expression -0.08846    0.24994  -0.354    0.723
##
## (Dispersion parameter for binomial family taken to be 1)
##
##     Null deviance: 99.880  on 85  degrees of freedom
## Residual deviance: 99.753  on 84  degrees of freedom
## AIC: 103.75
##
## Number of Fisher Scoring iterations: 4
```

In this univariate regression, there is a suggestion of a negative association between MK2 (treated as a continuous variable) and death at one year, in the early stage patients. That is, higher MK2 = lower odds of death at one year. the CIs are imprecise, and thus the p values is 0.06. We do not see this in the late-stage patients.

##### Multivariate regression

Now, we add covariates to the univariate model above. Again, because there is a baseline imbalance in transcript level between early and late stage, we are stratifying rather than adjusting. MK2 here remains a continuous variable, for now.

```
summary(glm(death_at_1year ~ MK2.Expression + smoking + age_at_diagnosis +
    gender, data = MK2_CoxPH_quantile[MK2_CoxPH_quantile$ajcc_pathologic_stage_combined ==
    "Early Stage", ], family = binomial(link = "logit")))
```

```
##
## Call:
## glm(formula = death_at_1year ~ MK2.Expression + smoking + age_at_diagnosis +
##     gender, family = binomial(link = "logit"), data = MK2_CoxPH_quantile[MK2_CoxPH_quantile$ajcc_pat
##     "Early Stage", ])
##
## Deviance Residuals:
##     Min       1Q   Median       3Q      Max
## -0.7519  -0.4579  -0.3556  -0.2933   2.5860
##
## Coefficients:
##                  Estimate Std. Error z value Pr(>|z|)
## (Intercept)      -1.01796    1.63391  -0.623   0.5333
## MK2.Expression   -0.32422    0.21042  -1.541   0.1234
## smoking           0.13949    0.49344   0.283   0.7774
## age_at_diagnosis -0.01141    0.02121  -0.538   0.5906
## gendermale        0.82992    0.43657   1.901   0.0573 .
## ---
## Signif. codes:  0 '***' 0.001 '**' 0.01 '*' 0.05 '.' 0.1 ' ' 1
##
## (Dispersion parameter for binomial family taken to be 1)
##
##     Null deviance: 174.15  on 311  degrees of freedom
```

```
## Residual deviance: 166.81  on 307  degrees of freedom
##   (7 observations deleted due to missingness)
## AIC: 176.81
##
## Number of Fisher Scoring iterations: 5
```

```
summary(glm(death_at_1year ~ MK2.Expression + smoking + age_at_diagnosis +
    gender, data = MK2_CoxPH_quantile[MK2_CoxPH_quantile$ajcc_pathologic_stage_combined ==
    "Late Stage", ], family = binomial(link = "logit")))
```

```
##
## Call:
## glm(formula = death_at_1year ~ MK2.Expression + smoking + age_at_diagnosis +
##     gender, family = binomial(link = "logit"), data = MK2_CoxPH_quantile[MK2_CoxPH_quantile$ajcc_path
##     "Late Stage", ])
##
## Deviance Residuals:
##     Min       1Q   Median       3Q      Max
## -1.1177  -0.8219  -0.7066   1.4406   1.9247
##
## Coefficients:
##                  Estimate Std. Error z value Pr(>|z|)
## (Intercept)      -2.64865    1.98157  -1.337    0.181
## MK2.Expression   -0.06162    0.26183  -0.235    0.814
## smoking          -0.42478    0.51632  -0.823    0.411
## age_at_diagnosis  0.03072    0.02594   1.184    0.236
## gendermale        0.33048    0.49610   0.666    0.505
##
## (Dispersion parameter for binomial family taken to be 1)
##
##     Null deviance: 99.880  on 85  degrees of freedom
## Residual deviance: 97.263  on 81  degrees of freedom
## AIC: 107.26
##
## Number of Fisher Scoring iterations: 4
```

Even with adjustment, there is again a suggestion of a negative association between MK2 transcript level and death at one year, with a p value of 0.1. There is a strong positive association between male gender and death at one year.

#### Model 2 in paper: Logistic regression for death at 1 year, stratified by early/late, but using Top 1/3 MK2 expression (dichotomous MK2 variable)

Now, we are interested in using MK2 as as dichotomous variable - high (top 1/3) or low (bottom 2/3). Clinically, with an appropriate reference cohort, this might be more translatable. Thus, we modeled death at one year, as a logistic regression, but now with MK2 as a dichotomous variable. The univariate analysis was significant, so here we run the multi-variate regression.

##### Model 2 - Univariate Regression

```
# LR stratified by Early/Late
summary(glm(death_at_1year ~ MK2_Expression_topQ, data = MK2_CoxPH_quantile[MK2_CoxPH_quantile$ajcc_path
    "Early Stage", ], family = binomial(link = "logit")))
```

```
##
```

```
## Call:
## glm(formula = death_at_1year ~ MK2_Expression_topQ, family = binomial(link = "logit"),
##     data = MK2_CoxPH_quantile[MK2_CoxPH_quantile$ajcc_pathologic_stage_combined ==
##         "Early Stage", ])
##
## Deviance Residuals:
##     Min       1Q   Median       3Q      Max
## -0.5127  -0.5127  -0.5127  -0.2289   2.7037
##
## Coefficients:
##                       Estimate Std. Error z value Pr(>|z|)
## (Intercept)            -1.9629     0.2136  -9.190  < 2e-16 ***
## MK2_Expression_topQ1  -1.6659     0.6227  -2.675  0.00747 **
## ---
## Signif. codes:  0 '***' 0.001 '**' 0.01 '*' 0.05 '.' 0.1 ' ' 1
##
## (Dispersion parameter for binomial family taken to be 1)
##
##     Null deviance: 189.71  on 318  degrees of freedom
## Residual deviance: 179.35  on 317  degrees of freedom
## AIC: 183.35
##
## Number of Fisher Scoring iterations: 6
```

```
summary(glm(death_at_1year ~ MK2_Expression_topQ, data = MK2_CoxPH_quantile[MK2_CoxPH_quantile$ajcc_pat
    "Late Stage", ], family = binomial(link = "logit")))
```

```
##
## Call:
## glm(formula = death_at_1year ~ MK2_Expression_topQ, family = binomial(link = "logit"),
##     data = MK2_CoxPH_quantile[MK2_CoxPH_quantile$ajcc_pathologic_stage_combined ==
##         "Late Stage", ])
##
## Deviance Residuals:
##     Min       1Q   Median       3Q      Max
## -0.8421  -0.8421  -0.8421   1.5550   1.9214
##
## Coefficients:
##                       Estimate Std. Error z value Pr(>|z|)
## (Intercept)            -0.8544     0.2670  -3.200  0.00137 **
## MK2_Expression_topQ1  -0.8196     0.6835  -1.199  0.23047
## ---
## Signif. codes:  0 '***' 0.001 '**' 0.01 '*' 0.05 '.' 0.1 ' ' 1
##
## (Dispersion parameter for binomial family taken to be 1)
##
##     Null deviance: 99.88  on 85  degrees of freedom
## Residual deviance: 98.26  on 84  degrees of freedom
## AIC: 102.26
##
## Number of Fisher Scoring iterations: 4
```

We see highly significant association between high MK2 expression (as a dichotomous variable) and death at 1 year, in early but not late stage cancer.

##### Model 2: Multivariate regression

So, now we will run 2 models. MK2 as dichotomous variable, with covariates, stratified by stage with the outcome of death at 1 year, as well as death at 2 years.

Here we first look at Early Stage only

```
logreg_topq1_early <- glm(death_at_1year ~ MK2_Expression_topQ + smoking +
    age_at_diagnosis + gender, data = MK2_CoxPH_quantile[MK2_CoxPH_quantile$ajcc_pathologic_stage_combi
    "Early Stage", ], family = binomial(link = "logit"))

summary(logreg_topq1_early)
```

```
##
## Call:
## glm(formula = death_at_1year ~ MK2_Expression_topQ + smoking +
##     age_at_diagnosis + gender, family = binomial(link = "logit"),
##     data = MK2_CoxPH_quantile[MK2_CoxPH_quantile$ajcc_pathologic_stage_combined ==
##         "Early Stage", ])
##
## Deviance Residuals:
##     Min       1Q   Median       3Q      Max
## -0.7059  -0.4897  -0.3689  -0.2019   2.8548
##
## Coefficients:
##                      Estimate Std. Error z value Pr(>|z|)
## (Intercept)          -1.62809    1.47184  -1.106   0.2687
## MK2_Expression_topQ1 -1.50091    0.63011  -2.382   0.0172 *
## smoking               0.16785    0.49734   0.337   0.7357
## age_at_diagnosis     -0.01533    0.02089  -0.734   0.4629
## gendermale            0.78866    0.43959   1.794   0.0728 .
## ---
## Signif. codes:  0 '***' 0.001 '**' 0.01 '*' 0.05 '.' 0.1 ' ' 1
##
## (Dispersion parameter for binomial family taken to be 1)
##
##     Null deviance: 174.15  on 311  degrees of freedom
## Residual deviance: 161.58  on 307  degrees of freedom
##   (7 observations deleted due to missingness)
## AIC: 171.58
##
## Number of Fisher Scoring iterations: 6
```

```
exp(logreg_topq1_early$coefficients)
```

```
##          (Intercept) MK2_Expression_topQ1              smoking
##            0.1963034            0.2229281            1.1827597
##     age_at_diagnosis           gendermale
##            0.9847829            2.2004382
```

```
exp(confint.lm(logreg_topq1_early))
```

```
##                           2.5 %    97.5 %
## (Intercept)          0.01084259 3.5540422
## MK2_Expression_topQ1 0.06451920 0.7702661
## smoking              0.44451696 3.1470578
```

```
## age_at_diagnosis     0.94512249 1.0261075
## gendermale           0.92651066 5.2259825
```

```
logreg_topq1_early_2y <- glm(death_at_2year ~ MK2_Expression_topQ + smoking +
    age_at_diagnosis + gender, data = MK2_CoxPH_quantile[MK2_CoxPH_quantile$ajcc_pathologic_stage_combi
    "Early Stage", ], family = binomial(link = "logit"))

summary(logreg_topq1_early_2y)
```

```
##
## Call:
## glm(formula = death_at_2year ~ MK2_Expression_topQ + smoking +
##     age_at_diagnosis + gender, family = binomial(link = "logit"),
##     data = MK2_CoxPH_quantile[MK2_CoxPH_quantile$ajcc_pathologic_stage_combined ==
##         "Early Stage", ])
##
## Deviance Residuals:
##     Min       1Q   Median       3Q      Max
## -0.8105  -0.6614  -0.5588  -0.4594   2.2780
##
## Coefficients:
##                      Estimate Std. Error z value Pr(>|z|)
## (Intercept)         -2.184387   1.112186  -1.964   0.0495 *
## MK2_Expression_topQ1 -0.695725   0.345540  -2.013   0.0441 *
## smoking              0.373370   0.360243   1.036   0.3000
## age_at_diagnosis     0.005672   0.015531   0.365   0.7150
## gendermale           0.389860   0.304645   1.280   0.2006
## ---
## Signif. codes:  0 '***' 0.001 '**' 0.01 '*' 0.05 '.' 0.1 ' ' 1
##
## (Dispersion parameter for binomial family taken to be 1)
##
##     Null deviance: 287.50  on 311  degrees of freedom
## Residual deviance: 279.64  on 307  degrees of freedom
##   (7 observations deleted due to missingness)
## AIC: 289.64
##
## Number of Fisher Scoring iterations: 4
```

```
exp(logreg_topq1_early_2y$coefficients)
```

```
##          (Intercept) MK2_Expression_topQ1              smoking
##            0.1125467            0.4987126            1.4526216
##     age_at_diagnosis           gendermale
##            1.0056877            1.4767739
```

```
exp(confint.lm(logreg_topq1_early_2y))
```

```
##                           2.5 %    97.5 %
## (Intercept)          0.01261513 1.0040923
## MK2_Expression_topQ1 0.25267507 0.9843244
## smoking              0.71498909 2.9512471
## age_at_diagnosis     0.97541719 1.0368977
## gendermale           0.81091195 2.6893934
```

Now we look at late stage

```r
logreg_topq1_late <- glm(death_at_1year ~ MK2_Expression_topQ + smoking +
    age_at_diagnosis + gender, data = MK2_CoxPH_quantile[MK2_CoxPH_quantile$ajcc_pathologic_stage_combi
    "Late Stage", ], family = binomial(link = "logit"))

summary(logreg_topq1_late)
```

```
##
## Call:
## glm(formula = death_at_1year ~ MK2_Expression_topQ + smoking +
##     age_at_diagnosis + gender, family = binomial(link = "logit"),
##     data = MK2_CoxPH_quantile[MK2_CoxPH_quantile$ajcc_pathologic_stage_combined ==
##         "Late Stage", ])
##
## Deviance Residuals:
##     Min       1Q   Median       3Q      Max
## -1.1548  -0.8278  -0.6714   1.3897   2.1483
##
## Coefficients:
##                      Estimate Std. Error z value Pr(>|z|)
## (Intercept)          -2.63173    1.80098  -1.461    0.144
## MK2_Expression_topQ1 -0.72657    0.69371  -1.047    0.295
## smoking              -0.38921    0.51949  -0.749    0.454
## age_at_diagnosis      0.02873    0.02663   1.079    0.281
## gendermale            0.33859    0.49884   0.679    0.497
##
## (Dispersion parameter for binomial family taken to be 1)
##
##     Null deviance: 99.880  on 85  degrees of freedom
## Residual deviance: 96.109  on 81  degrees of freedom
## AIC: 106.11
##
## Number of Fisher Scoring iterations: 4
```

```r
exp(logreg_topq1_late$coefficients)
```

```
##          (Intercept) MK2_Expression_topQ1              smoking
##           0.07195393           0.48356425           0.67759368
##     age_at_diagnosis           gendermale
##           1.02914235           1.40296293
```

```r
exp(confint.lm(logreg_topq1_late))
```

```
##                            2.5 %   97.5 %
## (Intercept)          0.001998996 2.589984
## MK2_Expression_topQ1 0.121623019 1.922616
## smoking              0.241030494 1.904876
## age_at_diagnosis     0.976038424 1.085136
## gendermale           0.519985525 3.785307
```

```r
logreg_topq1_late_2y <- glm(death_at_2year ~ MK2_Expression_topQ + smoking +
    age_at_diagnosis + gender, data = MK2_CoxPH_quantile[MK2_CoxPH_quantile$ajcc_pathologic_stage_combi
    "Late Stage", ], family = binomial(link = "logit"))

summary(logreg_topq1_late_2y)
```

```
##
## Call:
## glm(formula = death_at_2year ~ MK2_Expression_topQ + smoking +
##     age_at_diagnosis + gender, family = binomial(link = "logit"),
##     data = MK2_CoxPH_quantile[MK2_CoxPH_quantile$ajcc_pathologic_stage_combined ==
##         "Late Stage", ])
##
## Deviance Residuals:
##     Min       1Q   Median       3Q      Max
## -1.3149  -1.0925  -0.9229   1.2055   1.5937
##
## Coefficients:
##                      Estimate Std. Error z value Pr(>|z|)
## (Intercept)          -1.09315    1.47675  -0.740    0.459
## MK2_Expression_topQ1 -0.37233    0.54314  -0.686    0.493
## smoking              -0.25976    0.46684  -0.556    0.578
## age_at_diagnosis      0.01838    0.02209   0.832    0.405
## gendermale           -0.02267    0.43926  -0.052    0.959
##
## (Dispersion parameter for binomial family taken to be 1)
##
##     Null deviance: 118.48  on 85  degrees of freedom
## Residual deviance: 116.76  on 81  degrees of freedom
## AIC: 126.76
##
## Number of Fisher Scoring iterations: 4
```

```
exp(logreg_topq1_late_2y$coefficients)
```

```
##          (Intercept) MK2_Expression_topQ1              smoking
##            0.3351602            0.6891295            0.7712360
##     age_at_diagnosis           gendermale
##            1.0185536            0.9775869
```

```
exp(confint.lm(logreg_topq1_late_2y))
```

```
##                           2.5 %   97.5 %
## (Intercept)          0.01774918 6.328876
## MK2_Expression_topQ1 0.23386708 2.030639
## smoking              0.30464012 1.952484
## age_at_diagnosis     0.97476408 1.064310
## gendermale           0.40793462 2.342719
```

##### Model 2: MK2 multivariate model with Casp3

Now we run the multi-variate model again but with casp3 expression

```
summary(glm(death_at_2year ~ MK2_Expression_topQ + smoking + age_at_diagnosis +
    gender + Casp3.Expression, data = MK2_CoxPH_quantile[MK2_CoxPH_quantile$ajcc_pathologic_stage_combi
    "Early Stage", ], family = binomial(link = "logit")))
```

```
##
## Call:
## glm(formula = death_at_2year ~ MK2_Expression_topQ + smoking +
##     age_at_diagnosis + gender + Casp3.Expression, family = binomial(link = "logit"),
##     data = MK2_CoxPH_quantile[MK2_CoxPH_quantile$ajcc_pathologic_stage_combined ==
```

```
##         "Early Stage", ])
##
## Deviance Residuals:
##     Min       1Q   Median       3Q      Max
## -0.8350  -0.6598  -0.5608  -0.4615   2.2744
##
## Coefficients:
##                       Estimate Std. Error z value Pr(>|z|)
## (Intercept)          -2.255784   1.179108  -1.913   0.0557 .
## MK2_Expression_topQ1 -0.688504   0.347914  -1.979   0.0478 *
## smoking               0.373941   0.360311   1.038   0.2994
## age_at_diagnosis      0.005639   0.015532   0.363   0.7166
## gendermale            0.392415   0.305005   1.287   0.1982
## Casp3.Expression      0.080910   0.443139   0.183   0.8551
## ---
## Signif. codes:  0 '***' 0.001 '**' 0.01 '*' 0.05 '.' 0.1 ' ' 1
##
## (Dispersion parameter for binomial family taken to be 1)
##
##     Null deviance: 287.50  on 311  degrees of freedom
## Residual deviance: 279.61  on 306  degrees of freedom
##   (7 observations deleted due to missingness)
## AIC: 291.61
##
## Number of Fisher Scoring iterations: 4
```

Addition of Casp3 does not change the effect of MK2_expression on death at 2 year in this model.

### Cox Proportional Hazards modeling, with right censoring at 1 year.

#### Data Cleaning

```
# data cleaning prior to Cox PH model
MK2_CoxPH_quantile$MK2_Expression_topQ <- as.factor(unlist(MK2_CoxPH_quantile$MK2_Expression_topQ))
MK2_CoxPH_quantile$gender <- as.factor(MK2_CoxPH_quantile$gender)

# be careful here. as.numeric alone will return the factor level, not
# the value!
MK2_CoxPH_quantile$censor <- as.numeric(as.character(MK2_CoxPH_quantile$death_at_1year))
MK2_CoxPH_quantile$censor.2y <- as.numeric(as.character(MK2_CoxPH_quantile$death_at_2year))
MK2_CoxPH_quantile$censor.3y <- as.numeric(as.character(MK2_CoxPH_quantile$death_at_3year))

# there should be no discrepancy - only two dots should show up
plot(MK2_CoxPH_quantile$censor, MK2_CoxPH_quantile$death_at_1year)
```

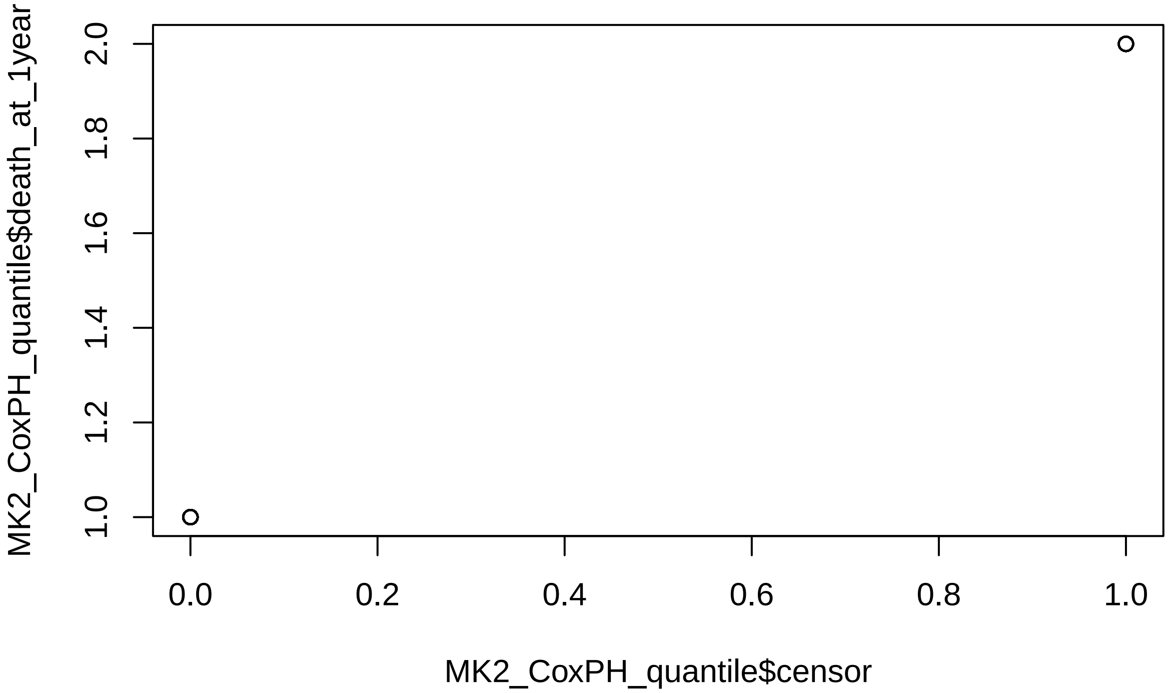

##### Cox PH model R censored at one year, stratified by early/late, MK2 as a continuous variable

###### Model 1: MK2 only

```
# univariate Cox PH
summary(coxph(Surv(time, censor) ~ MK2.Expression, id = submitter_id, data = MK2_CoxPH_quantile[MK2_CoxI
    "Early Stage", ]))
```

```
## Call:
## coxph(formula = Surv(time, censor) ~ MK2.Expression, data = MK2_CoxPH_quantile[MK2_CoxPH_quantile$aj
##     "Early Stage", ], id = submitter_id)
##
##   n= 319, number of events= 28
##
##                   coef exp(coef) se(coef)      z Pr(>|z|)
## MK2.Expression -0.3488    0.7055   0.1930 -1.808   0.0707 .
## ---
## Signif. codes:  0 '***' 0.001 '**' 0.01 '*' 0.05 '.' 0.1 ' ' 1
##
##                exp(coef) exp(-coef) lower .95 upper .95
## MK2.Expression    0.7055      1.417    0.4833      1.03
##
## Concordance= 0.606  (se = 0.046 )
## Likelihood ratio test= 3.45  on 1 df,   p=0.06
## Wald test            = 3.27  on 1 df,   p=0.07
## Score (logrank) test = 3.25  on 1 df,   p=0.07
```

```
# Multi-variate Cox PH
MK2_coxph_model_tb_1y_early <- coxph(Surv(time, censor) ~ MK2.Expression +
    gender + smoking + age_at_diagnosis, id = submitter_id, data = MK2_CoxPH_quantile[MK2_CoxPH_quantil
    "Early Stage", ])
```

```r
MK2_coxph_model_tb_1y_late <- coxph(Surv(time, censor) ~ MK2.Expression +
    gender + smoking + age_at_diagnosis, id = submitter_id, data = MK2_CoxPH_quantile[MK2_CoxPH_quantil
    "Late Stage", ])

# PH testing
cox.zph(MK2_coxph_model_tb_1y_early)
```

```
##                  chisq df    p
## MK2.Expression   1.522  1 0.22
## gender           0.419  1 0.52
## smoking          3.276  1 0.07
## age_at_diagnosis 1.616  1 0.20
## GLOBAL           6.371  4 0.17
```

```r
cox.zph(MK2_coxph_model_tb_1y_late)
```

```
##                   chisq df    p
## MK2.Expression   0.3655  1 0.55
## gender           1.2832  1 0.26
## smoking          0.2681  1 0.60
## age_at_diagnosis 0.0349  1 0.85
## GLOBAL           1.9921  4 0.74
```

```r
# results
summary(MK2_coxph_model_tb_1y_early)
```

```
## Call:
## coxph(formula = Surv(time, censor) ~ MK2.Expression + gender +
##     smoking + age_at_diagnosis, data = MK2_CoxPH_quantile[MK2_CoxPH_quantile$ajcc_pathologic_stage_c
##     "Early Stage", ], id = submitter_id)
##
##   n= 312, number of events= 25
##    (7 observations deleted due to missingness)
##
##                      coef exp(coef) se(coef)      z Pr(>|z|)
## MK2.Expression   -0.27510   0.75949  0.19775 -1.391   0.1642
## gendermale        0.83411   2.30276  0.41878  1.992   0.0464 *
## smoking           0.06909   1.07153  0.47106  0.147   0.8834
## age_at_diagnosis -0.01332   0.98676  0.02050 -0.650   0.5158
## ---
## Signif. codes:  0 '***' 0.001 '**' 0.01 '*' 0.05 '.' 0.1 ' ' 1
##
##                  exp(coef) exp(-coef) lower .95 upper .95
## MK2.Expression      0.7595     1.3167    0.5155     1.119
## gendermale          2.3028     0.4343    1.0134     5.233
## smoking             1.0715     0.9332    0.4256     2.698
## age_at_diagnosis    0.9868     1.0134    0.9479     1.027
##
## Concordance= 0.644  (se = 0.058 )
## Likelihood ratio test= 7.33  on 4 df,   p=0.1
## Wald test            = 7.09  on 4 df,   p=0.1
## Score (logrank) test = 7.36  on 4 df,   p=0.1
```

```r
summary(MK2_coxph_model_tb_1y_late)
```

```
## Call:
## coxph(formula = Surv(time, censor) ~ MK2.Expression + gender +
##     smoking + age_at_diagnosis, data = MK2_CoxPH_quantile[MK2_CoxPH_quantile$ajcc_pathologic_stage_c
##     "Late Stage", ], id = submitter_id)
##
##   n= 86, number of events= 23
##
##                       coef exp(coef) se(coef)      z Pr(>|z|)
## MK2.Expression   -0.07991   0.92320  0.21887 -0.365    0.715
## gendermale        0.20038   1.22187  0.41969  0.477    0.633
## smoking          -0.24635   0.78165  0.43026 -0.573    0.567
## age_at_diagnosis  0.02883   1.02925  0.02252  1.280    0.201
##
##                  exp(coef) exp(-coef) lower .95 upper .95
## MK2.Expression      0.9232     1.0832    0.6012     1.418
## gendermale          1.2219     0.8184    0.5368     2.781
## smoking             0.7817     1.2793    0.3363     1.817
## age_at_diagnosis    1.0293     0.9716    0.9848     1.076
##
## Concordance= 0.601  (se = 0.058 )
## Likelihood ratio test= 2.58  on 4 df,   p=0.6
## Wald test            = 2.39  on 4 df,   p=0.7
## Score (logrank) test = 2.43  on 4 df,   p=0.7
```

So here, we can see that in multi-variate Cox PH model, we see a signal for MK2 and improved survival, but only in the early stage tumors.

##### MK2 with casp 3

```r
summary(coxph(Surv(time, censor) ~ MK2.Expression + gender + smoking +
    age_at_diagnosis + Casp3.Expression, id = submitter_id, data = MK2_CoxPH_quantile[MK2_CoxPH_quantil
    "Early Stage", ]))
```

```
## Call:
## coxph(formula = Surv(time, censor) ~ MK2.Expression + gender +
##     smoking + age_at_diagnosis + Casp3.Expression, data = MK2_CoxPH_quantile[MK2_CoxPH_quantile$ajcc_
##     "Early Stage", ], id = submitter_id)
##
##   n= 312, number of events= 25
##    (7 observations deleted due to missingness)
##
##                       coef exp(coef) se(coef)      z Pr(>|z|)
## MK2.Expression   -0.25541   0.77460  0.20031 -1.275   0.2023
## gendermale        0.84606   2.33045  0.41925  2.018   0.0436 *
## smoking           0.07316   1.07591  0.47159  0.155   0.8767
## age_at_diagnosis -0.01301   0.98707  0.02050 -0.635   0.5256
## Casp3.Expression  0.30076   1.35089  0.58307  0.516   0.6060
## ---
## Signif. codes:  0 '***' 0.001 '**' 0.01 '*' 0.05 '.' 0.1 ' ' 1
##
##                  exp(coef) exp(-coef) lower .95 upper .95
## MK2.Expression      0.7746     1.2910    0.5231     1.147
```

```
## gendermale          2.3305     0.4291    1.0247     5.300
## smoking             1.0759     0.9294    0.4269     2.711
## age_at_diagnosis    0.9871     1.0131    0.9482     1.028
## Casp3.Expression    1.3509     0.7403    0.4308     4.236
##
## Concordance= 0.646  (se = 0.059 )
## Likelihood ratio test= 7.58  on 5 df,   p=0.2
## Wald test            = 7.38  on 5 df,   p=0.2
## Score (logrank) test = 7.67  on 5 df,   p=0.2
```

Similar to our logistic reg model, no real difference with Casp3 addition to the model.

##### Model 1 in paper: Cox PH model R censored at one year and two year, stratified by early late, MK2 as dichotomous (top 1/3 vs. bottom 2/3)

###### Model 1: MK2+other covariates, at 1 year and 2 year

```
summary(coxph(Surv(time, censor) ~ MK2_Expression_topQ + gender + smoking +
    age_at_diagnosis, id = submitter_id, data = MK2_CoxPH_quantile[MK2_CoxPH_quantile$ajcc_pathologic_s
    "Early Stage", ]))
```

```
## Call:
## coxph(formula = Surv(time, censor) ~ MK2_Expression_topQ + gender +
##     smoking + age_at_diagnosis, data = MK2_CoxPH_quantile[MK2_CoxPH_quantile$ajcc_pathologic_stage_c
##     "Early Stage", ], id = submitter_id)
##
##   n= 312, number of events= 25
##    (7 observations deleted due to missingness)
##
##                          coef exp(coef) se(coef)      z Pr(>|z|)
## MK2_Expression_topQ1 -1.42504   0.24050  0.61622 -2.313   0.0207 *
## gendermale            0.81309   2.25486  0.41812  1.945   0.0518 .
## smoking               0.06489   1.06704  0.47247  0.137   0.8908
## age_at_diagnosis     -0.01696   0.98318  0.02001 -0.848   0.3966
## ---
## Signif. codes:  0 '***' 0.001 '**' 0.01 '*' 0.05 '.' 0.1 ' ' 1
##
##                      exp(coef) exp(-coef) lower .95 upper .95
## MK2_Expression_topQ1    0.2405     4.1580   0.07188    0.8047
## gendermale              2.2549     0.4435   0.99361    5.1171
## smoking                 1.0670     0.9372   0.42268    2.6937
## age_at_diagnosis        0.9832     1.0171   0.94536    1.0225
##
## Concordance= 0.701  (se = 0.053 )
## Likelihood ratio test= 12.76  on 4 df,   p=0.01
## Wald test            = 10.18  on 4 df,   p=0.04
## Score (logrank) test = 11.53  on 4 df,   p=0.02
```

```
summary(coxph(Surv(time, censor) ~ MK2_Expression_topQ + gender + smoking +
    age_at_diagnosis, id = submitter_id, data = MK2_CoxPH_quantile[MK2_CoxPH_quantile$ajcc_pathologic_s
    "Late Stage", ]))
```

```
## Call:
## coxph(formula = Surv(time, censor) ~ MK2_Expression_topQ + gender +
##     smoking + age_at_diagnosis, data = MK2_CoxPH_quantile[MK2_CoxPH_quantile$ajcc_pathologic_stage_c
```

```
##     "Late Stage", ], id = submitter_id)
##
##   n= 86, number of events= 23
##
##                          coef exp(coef) se(coef)      z Pr(>|z|)
## MK2_Expression_topQ1 -0.71417   0.48960  0.62288 -1.147    0.252
## gendermale            0.19689   1.21761  0.41857  0.470    0.638
## smoking              -0.20205   0.81705  0.43065 -0.469    0.639
## age_at_diagnosis      0.02776   1.02815  0.02282  1.217    0.224
##
##                      exp(coef) exp(-coef) lower .95 upper .95
## MK2_Expression_topQ1    0.4896     2.0425    0.1444     1.660
## gendermale              1.2176     0.8213    0.5361     2.766
## smoking                 0.8171     1.2239    0.3513     1.900
## age_at_diagnosis        1.0281     0.9726    0.9832     1.075
##
## Concordance= 0.626  (se = 0.058 )
## Likelihood ratio test= 4  on 4 df,   p=0.4
## Wald test            = 3.51  on 4 df,   p=0.5
## Score (logrank) test = 3.6  on 4 df,   p=0.5
```

```
summary(coxph(Surv(time, censor.2y) ~ MK2_Expression_topQ + gender + smoking +
    age_at_diagnosis, id = submitter_id, data = MK2_CoxPH_quantile[MK2_CoxPH_quantile$ajcc_pathologic_s
    "Early Stage", ]))
```

```
## Call:
## coxph(formula = Surv(time, censor.2y) ~ MK2_Expression_topQ +
##     gender + smoking + age_at_diagnosis, data = MK2_CoxPH_quantile[MK2_CoxPH_quantile$ajcc_pathologi
##     "Early Stage", ], id = submitter_id)
##
##   n= 312, number of events= 54
##    (7 observations deleted due to missingness)
##
##                           coef exp(coef)  se(coef)      z Pr(>|z|)
## MK2_Expression_topQ1 -0.643511  0.525444  0.319120 -2.017   0.0437 *
## gendermale            0.420566  1.522823  0.273765  1.536   0.1245
## smoking               0.309145  1.362260  0.330173  0.936   0.3491
## age_at_diagnosis      0.004461  1.004471  0.014624  0.305   0.7603
## ---
## Signif. codes:  0 '***' 0.001 '**' 0.01 '*' 0.05 '.' 0.1 ' ' 1
##
##                      exp(coef) exp(-coef) lower .95 upper .95
## MK2_Expression_topQ1    0.5254     1.9032    0.2811    0.9821
## gendermale              1.5228     0.6567    0.8905    2.6042
## smoking                 1.3623     0.7341    0.7132    2.6020
## age_at_diagnosis        1.0045     0.9955    0.9761    1.0337
##
## Concordance= 0.624  (se = 0.038 )
## Likelihood ratio test= 8.26  on 4 df,   p=0.08
## Wald test            = 7.82  on 4 df,   p=0.1
## Score (logrank) test = 8.05  on 4 df,   p=0.09
```

```
summary(coxph(Surv(time, censor.2y) ~ MK2_Expression_topQ + gender + smoking +
    age_at_diagnosis, id = submitter_id, data = MK2_CoxPH_quantile[MK2_CoxPH_quantile$ajcc_pathologic_s
    "Late Stage", ]))
```

```
## Call:
## coxph(formula = Surv(time, censor.2y) ~ MK2_Expression_topQ +
##     gender + smoking + age_at_diagnosis, data = MK2_CoxPH_quantile[MK2_CoxPH_quantile$ajcc_pathologi
##     "Late Stage", ], id = submitter_id)
##
##   n= 86, number of events= 39
##
##                          coef exp(coef)  se(coef)      z Pr(>|z|)
## MK2_Expression_topQ1 -0.483550  0.616590  0.420764 -1.149    0.250
## gendermale            0.008068  1.008101  0.322004  0.025    0.980
## smoking              -0.103562  0.901620  0.339616 -0.305    0.760
## age_at_diagnosis      0.018713  1.018889  0.016820  1.113    0.266
##
##                      exp(coef) exp(-coef) lower .95 upper .95
## MK2_Expression_topQ1    0.6166     1.6218    0.2703     1.407
## gendermale              1.0081     0.9920    0.5363     1.895
## smoking                 0.9016     1.1091    0.4634     1.754
## age_at_diagnosis        1.0189     0.9815    0.9858     1.053
##
## Concordance= 0.604  (se = 0.048 )
## Likelihood ratio test= 3.2  on 4 df,   p=0.5
## Wald test            = 2.95  on 4 df,   p=0.6
## Score (logrank) test = 2.99  on 4 df,   p=0.6
```

**Model 1: MK2 + other covariates + casp3, at 1 year and 2 year**

```
summary(coxph(Surv(time, censor) ~ MK2_Expression_topQ + gender + smoking +
    age_at_diagnosis + Casp3.Expression, id = submitter_id, data = MK2_CoxPH_quantile[MK2_CoxPH_quantil
    "Early Stage", ]))
```

```
## Call:
## coxph(formula = Surv(time, censor) ~ MK2_Expression_topQ + gender +
##     smoking + age_at_diagnosis + Casp3.Expression, data = MK2_CoxPH_quantile[MK2_CoxPH_quantile$ajcc
##     "Early Stage", ], id = submitter_id)
##
##   n= 312, number of events= 25
##    (7 observations deleted due to missingness)
##
##                         coef exp(coef) se(coef)      z Pr(>|z|)
## MK2_Expression_topQ1 -1.40534   0.24528  0.61767 -2.275   0.0229 *
## gendermale            0.82233   2.27580  0.41871  1.964   0.0495 *
## smoking               0.07424   1.07707  0.47357  0.157   0.8754
## age_at_diagnosis     -0.01664   0.98350  0.01999 -0.832   0.4051
## Casp3.Expression      0.30803   1.36074  0.56469  0.545   0.5854
## ---
## Signif. codes:  0 '***' 0.001 '**' 0.01 '*' 0.05 '.' 0.1 ' ' 1
##
##                      exp(coef) exp(-coef) lower .95 upper .95
## MK2_Expression_topQ1    0.2453     4.0769    0.0731    0.8231
## gendermale              2.2758     0.4394    1.0017    5.1706
## smoking                 1.0771     0.9284    0.4257    2.7248
## age_at_diagnosis        0.9835     1.0168    0.9457    1.0228
## Casp3.Expression        1.3607     0.7349    0.4499    4.1157
##
```

```
## Concordance= 0.707  (se = 0.054 )
## Likelihood ratio test= 13.04  on 5 df,   p=0.02
## Wald test            = 10.5  on 5 df,   p=0.06
## Score (logrank) test = 11.81  on 5 df,   p=0.04
```

So, the same pattern here - we don't really see an impact of adding casp3 to the multivariable Cox PH model.

#### Model Graphics

##### KM curve - here, we're going to stratify MK2 by "hi" vs. "low" - Model 1

```
library(survminer)
library(survival)
library(ggsci)
km_early <- survfit(Surv(time, censor) ~ MK2_Expression_topQ, data = MK2_CoxPH_quantile[MK2_CoxPH_quant
    "Early Stage", ], type = "kaplan-meier")
ggsurvplot(km_early, xlim = c(0, 12), conf.int = TRUE, break.x.by = 2,
    pval = TRUE, font.y = 14, font.x = 14, font.tickslab = 14, ggtheme = theme_bw())
```

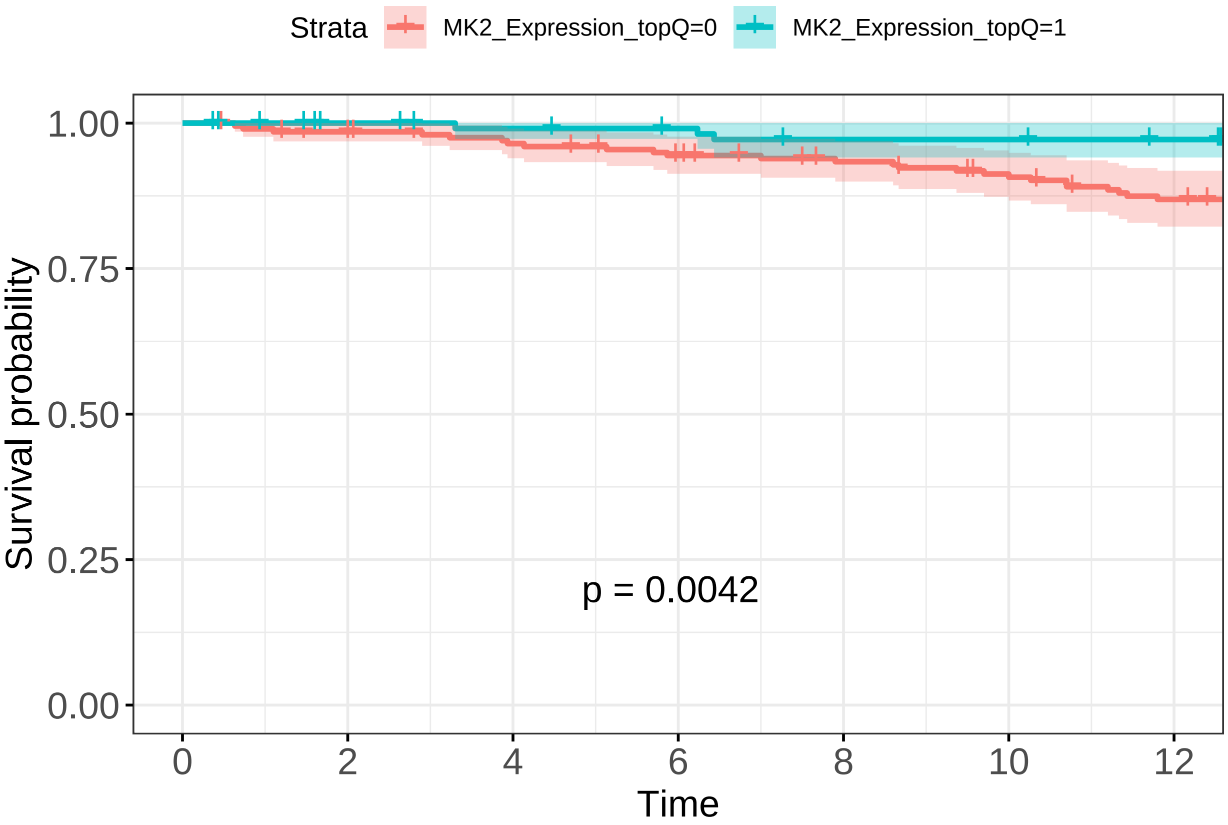

```
km_late <- survfit(Surv(time, censor) ~ MK2_Expression_topQ, data = MK2_CoxPH_quantile[MK2_CoxPH_quanti
    "Late Stage", ], type = "kaplan-meier")
ggsurvplot(km_late, xlim = c(0, 12), conf.int = TRUE, break.x.by = 2, pval = TRUE,
    font.y = 14, font.x = 14, font.tickslab = 14, ggtheme = theme_bw())
```

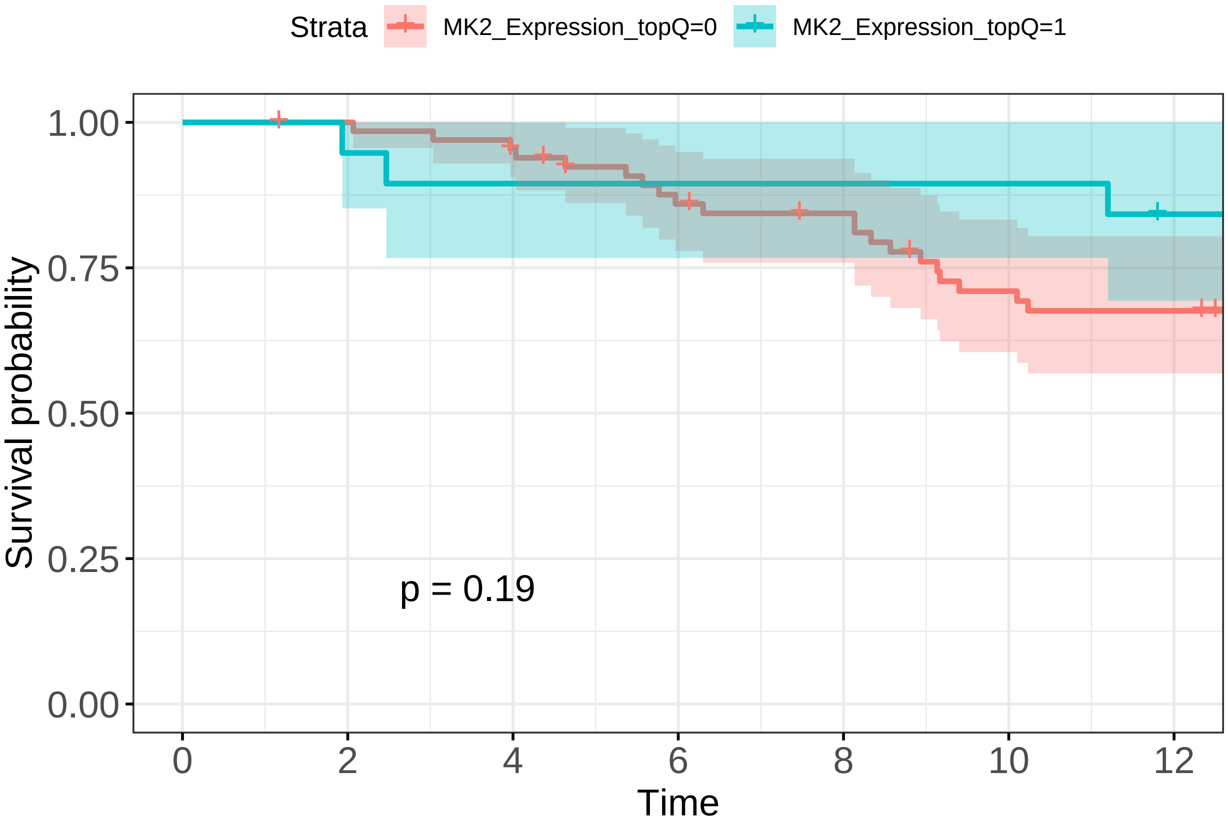

```
km_early.2y <- survfit(Surv(time, censor.2y) ~ MK2_Expression_topQ, data = MK2_CoxPH_quantile[MK2_CoxPH_
    "Early Stage", ], type = "kaplan-meier")
ggsurvplot(km_early.2y, xlim = c(0, 24), conf.int = TRUE, break.x.by = 2,
    pval = TRUE, font.y = 14, font.x = 14, font.tickslab = 14, ggtheme = theme_bw())
```

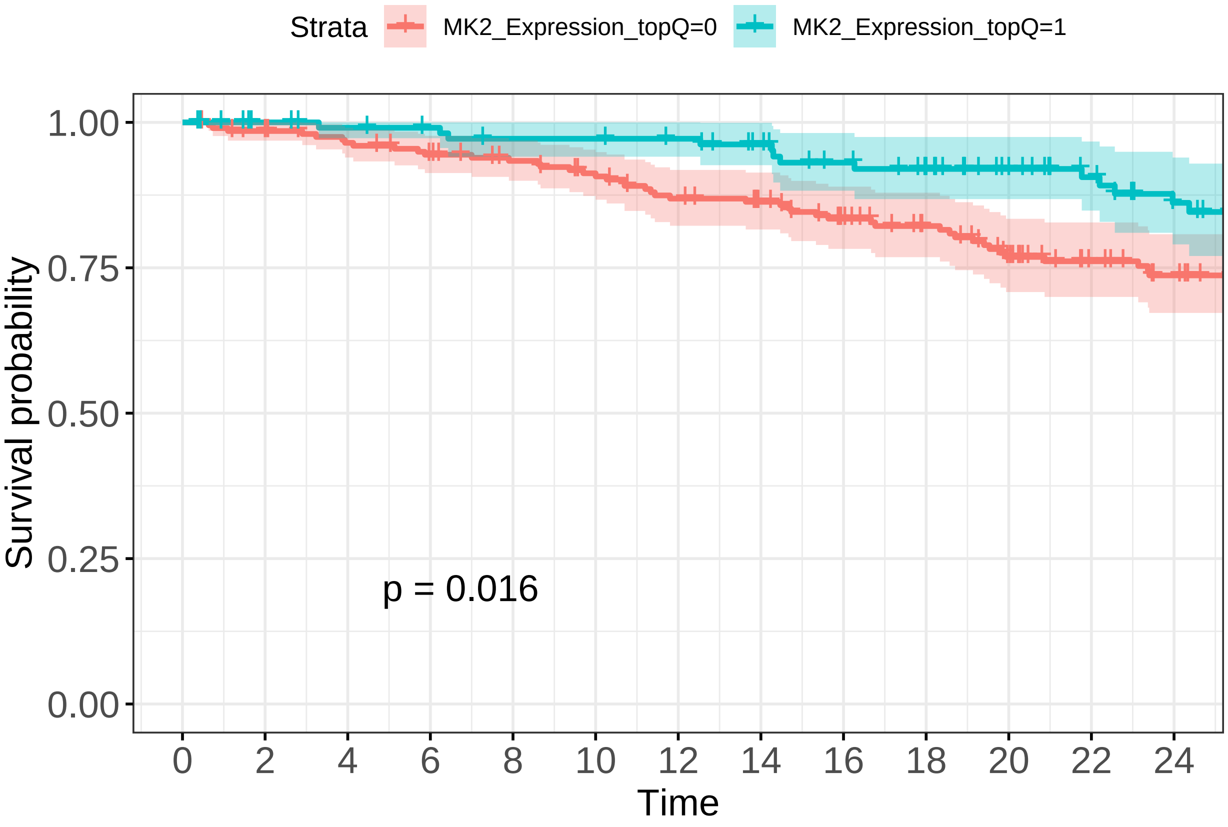

```
km_late.2y <- survfit(Surv(time, censor.2y) ~ MK2_Expression_topQ, data = MK2_CoxPH_quantile[MK2_CoxPH_q
    "Late Stage", ], type = "kaplan-meier")
ggsurvplot(km_late.2y, xlim = c(0, 24), conf.int = TRUE, break.x.by = 2,
    pval = TRUE, font.y = 14, font.x = 14, font.tickslab = 14, ggtheme = theme_bw())
```

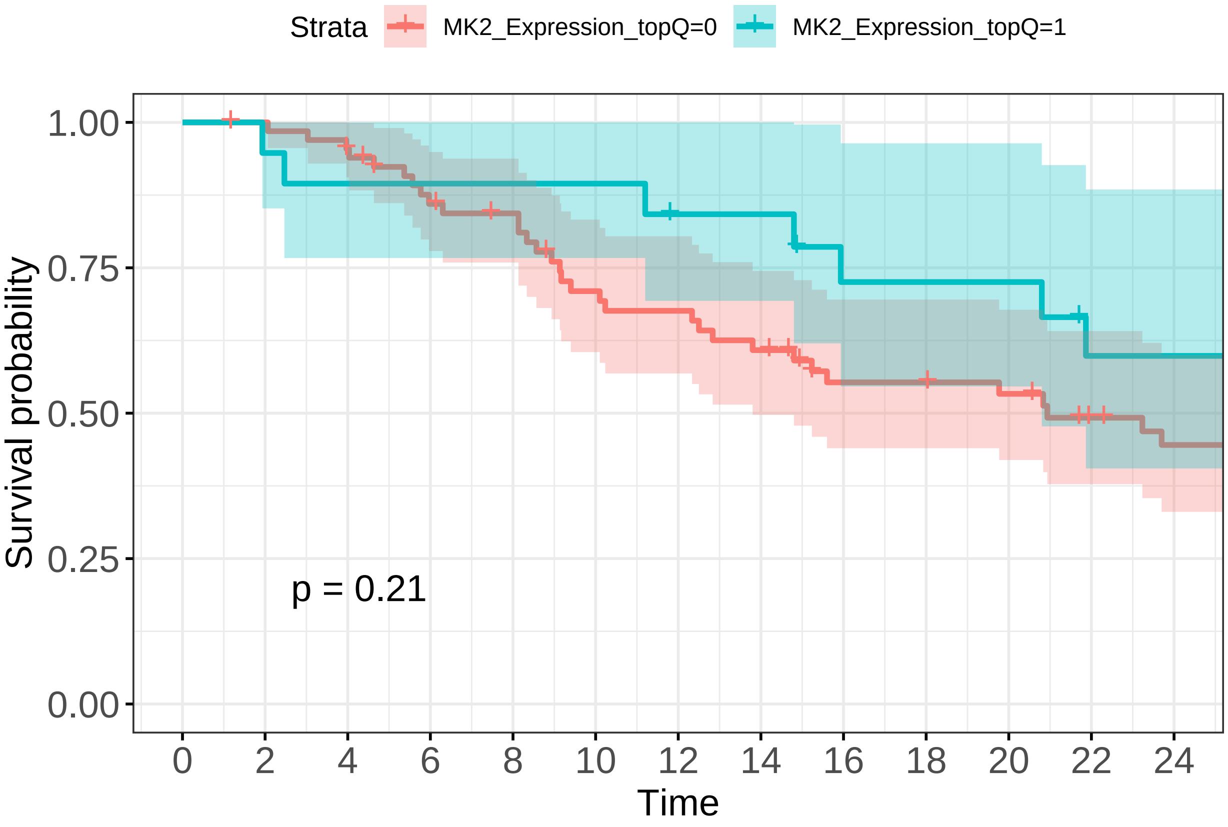

##### Predicted probabilities graphs for Logistic Regression (Model 2)

```
library(effects)
```

```
## Loading required package: carData
```

```
## lattice theme set by effectsTheme()
## See ?effectsTheme for details.
```

```
library(ggthemes)
MK2efx_early <- Effect("MK2_Expression_topQ", logreg_topq1_early_2y)
MK2efx_late <- Effect("MK2_Expression_topQ", logreg_topq1_late_2y)

MK2efx_early <- as.data.frame(MK2efx_early)
MK2efx_late <- as.data.frame(MK2efx_late)
MK2efx_early$stage <- "Early"
MK2efx_late$stage <- "Late"
MK2efx <- rbind(MK2efx_early, MK2efx_late)
MK2efx$MK2_Expression_topQ <- lapply(MK2efx$MK2_Expression_topQ, function(x) if (x ==
    0) return("Low") else return("High"))
MK2efx$MK2_Expression_topQ <- as.factor(unlist(MK2efx$MK2_Expression_topQ))
ggplot(MK2efx, aes(x = stage, y = fit, group = MK2_Expression_topQ, color = MK2_Expression_topQ,
    ymin = lower, ymax = upper)) + geom_point(position = position_dodge(width = 0.2)) +
    geom_errorbar(width = 0, position = position_dodge(width = 0.2)) +
    theme_bw(base_size = 14) + ylab("Predicted  Probability of death at 2 years") +
    coord_flip()
```

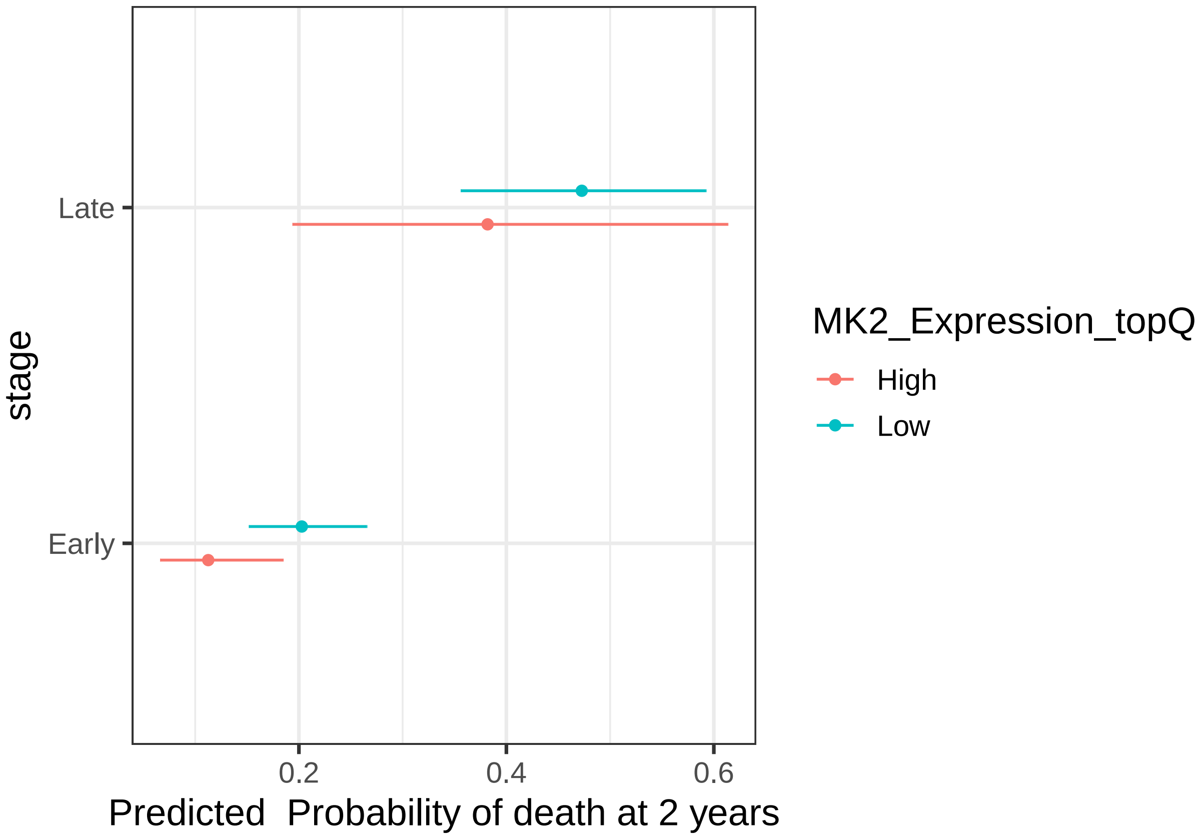

#

Demographics / Table 1

```
library(tableone)
listVars <- c("age_at_diagnosis", "gender", "smoking", "ajcc_pathologic_stage",
    "ajcc_pathologic_n", "prior_malignancy", "primary_diagnosis", "time",
    "race", "MK2.Expression")
catVars <- c("gender", "smoking", "ajcc_pathologic_stage", "ajcc_pathologic_n",
    "prior_malignancy", "primary_diagnosis", "race")

# Table 1 where we stratify by Early/Late Stage
table1 <- CreateTableOne(listVars, clinical, catVars, strata = c("ajcc_pathologic_stage_combined"),
    addOverall = TRUE)
clinical$MK2_Expression_topQ_t1 <- unlist(clinical$MK2_Expression_topQ)
kable(print(table1))
```

```
##                                                 Stratified by ajcc_pathologic_stage_combined
##                                                  Overall
##    n                                                483
##    age_at_diagnosis (mean (SD))                   65.77 (9.97)
##    gender = male (%)                                219 (45.3)
##    smoking = 1 (%)                                  331 (68.5)
##    ajcc_pathologic_stage (%)
##       Stage I                                         5 ( 1.0)
##       Stage IA                                      129 (26.7)
##       Stage IB                                      128 (26.5)
##       Stage II                                        1 ( 0.2)
##       Stage IIA                                      50 (10.4)
##       Stage IIB                                      67 (13.9)
```

```
##     Stage IIIA                                      69 (14.3)
##     Stage IIIB                                      10 ( 2.1)
##     Stage IV                                        24 ( 5.0)
##   ajcc_pathologic_n (%)
##     N0                                             313 (64.8)
##     N1                                              90 (18.6)
##     N2                                              68 (14.1)
##     N3                                               2 ( 0.4)
##     NX                                              10 ( 2.1)
##   prior_malignancy = yes (%)                        79 (16.4)
##   primary_diagnosis (%)
##     Acinar cell carcinoma                           22 ( 4.6)
##     Adenocarcinoma with mixed subtypes             103 (21.3)
##     Adenocarcinoma, NOS                            289 (59.8)
##     Bronchio-alveolar carcinoma, mucinous            5 ( 1.0)
##     Bronchiolo-alveolar adenocarcinoma, NOS          3 ( 0.6)
##     Bronchiolo-alveolar carcinoma, non-mucinous     17 ( 3.5)
##     Clear cell adenocarcinoma, NOS                   2 ( 0.4)
##     Micropapillary carcinoma, NOS                    2 ( 0.4)
##     Mucinous adenocarcinoma                         13 ( 2.7)
##     Papillary adenocarcinoma, NOS                   21 ( 4.3)
##     Signet ring cell carcinoma                       1 ( 0.2)
##     Solid carcinoma, NOS                             5 ( 1.0)
##   time (mean (SD))                              879.55 (873.37)
##   race (%)
##     american indian or alaska native                 1 ( 0.2)
##     asian                                            8 ( 1.7)
##     black or african american                       50 (10.4)
##     not reported                                    44 ( 9.1)
##     white                                          380 (78.7)
##   MK2.Expression (mean (SD))                      3.86 (1.05)
##                                                 Stratified by ajcc_pathologic_stage_combined
##                                                  Early Stage
##   n                                                380
##   age_at_diagnosis (mean (SD))                   66.01 (9.71)
##   gender = male (%)                                171 (45.0)
##   smoking = 1 (%)                                  259 (68.2)
##   ajcc_pathologic_stage (%)
##     Stage I                                          5 ( 1.3)
##     Stage IA                                       129 (33.9)
##     Stage IB                                       128 (33.7)
##     Stage II                                         1 ( 0.3)
##     Stage IIA                                       50 (13.2)
##     Stage IIB                                       67 (17.6)
##     Stage IIIA                                       0 ( 0.0)
##     Stage IIIB                                       0 ( 0.0)
##     Stage IV                                         0 ( 0.0)
##   ajcc_pathologic_n (%)
##     N0                                             297 (78.2)
##     N1                                              77 (20.3)
##     N2                                               0 ( 0.0)
##     N3                                               0 ( 0.0)
##     NX                                               6 ( 1.6)
##   prior_malignancy = yes (%)                        65 (17.1)
```

```
##   primary_diagnosis (%)
##      Acinar cell carcinoma                              16 ( 4.2)
##      Adenocarcinoma with mixed subtypes                 77 (20.3)
##      Adenocarcinoma, NOS                               230 (60.5)
##      Bronchio-alveolar carcinoma, mucinous               5 ( 1.3)
##      Bronchiolo-alveolar adenocarcinoma, NOS             3 ( 0.8)
##      Bronchiolo-alveolar carcinoma, non-mucinous        13 ( 3.4)
##      Clear cell adenocarcinoma, NOS                      1 ( 0.3)
##      Micropapillary carcinoma, NOS                       2 ( 0.5)
##      Mucinous adenocarcinoma                            12 ( 3.2)
##      Papillary adenocarcinoma, NOS                      15 ( 3.9)
##      Signet ring cell carcinoma                          1 ( 0.3)
##      Solid carcinoma, NOS                                5 ( 1.3)
##   time (mean (SD))                                   929.04 (921.78)
##   race (%)
##      american indian or alaska native                    1 ( 0.3)
##      asian                                               6 ( 1.6)
##      black or african american                          41 (10.8)
##      not reported                                       30 ( 7.9)
##      white                                             302 (79.5)
##   MK2.Expression (mean (SD))                           3.91 (1.06)
##                                                     Stratified by ajcc_pathologic_stage_combined
##                                                      Late Stage      p      test
##   n                                                    103
##   age_at_diagnosis (mean (SD))                        64.89 (10.86)   0.314
##   gender = male (%)                                     48 (46.6)    0.859
##   smoking = 1 (%)                                       72 (69.9)    0.827
##   ajcc_pathologic_stage (%)                                          <0.001
##      Stage I                                             0 ( 0.0)
##      Stage IA                                            0 ( 0.0)
##      Stage IB                                            0 ( 0.0)
##      Stage II                                            0 ( 0.0)
##      Stage IIA                                           0 ( 0.0)
##      Stage IIB                                           0 ( 0.0)
##      Stage IIIA                                         69 (67.0)
##      Stage IIIB                                         10 ( 9.7)
##      Stage IV                                           24 (23.3)
##   ajcc_pathologic_n (%)                                              <0.001
##      N0                                                 16 (15.5)
##      N1                                                 13 (12.6)
##      N2                                                 68 (66.0)
##      N3                                                  2 ( 1.9)
##      NX                                                  4 ( 3.9)
##   prior_malignancy = yes (%)                            14 (13.6)    0.481
##   primary_diagnosis (%)                                               0.621
##      Acinar cell carcinoma                               6 ( 5.8)
##      Adenocarcinoma with mixed subtypes                 26 (25.2)
##      Adenocarcinoma, NOS                                59 (57.3)
##      Bronchio-alveolar carcinoma, mucinous               0 ( 0.0)
##      Bronchiolo-alveolar adenocarcinoma, NOS             0 ( 0.0)
##      Bronchiolo-alveolar carcinoma, non-mucinous         4 ( 3.9)
##      Clear cell adenocarcinoma, NOS                      1 ( 1.0)
##      Micropapillary carcinoma, NOS                       0 ( 0.0)
##      Mucinous adenocarcinoma                             1 ( 1.0)
```

```
##      Papillary adenocarcinoma, NOS                 6 ( 5.8)
##      Signet ring cell carcinoma                    0 ( 0.0)
##      Solid carcinoma, NOS                          0 ( 0.0)
##   time (mean (SD))                            696.95 (636.03)  0.017
##   race (%)                                                     0.449
##      american indian or alaska native              0 ( 0.0)
##      asian                                         2 ( 1.9)
##      black or african american                     9 ( 8.7)
##      not reported                                 14 (13.6)
##      white                                        78 (75.7)
##   MK2.Expression (mean (SD))                    3.66 (0.99)    0.032
```

| | Overall | Early Stage | Late Stage | p | test |
|---|---|---|---|---|---|
| n | 483 | 380 | 103 | | |
| age_at_diagnosis (mean (SD)) | 65.77 (9.97) | 66.01 (9.71) | 64.89 (10.86) | 0.314 | |
| gender = male (%) | 219 (45.3) | 171 (45.0) | 48 (46.6) | 0.859 | |
| smoking = 1 (%) | 331 (68.5) | 259 (68.2) | 72 (69.9) | 0.827 | |
| ajcc_pathologic_stage (%) | | | | <0.001 | |
| Stage I | 5 ( 1.0) | 5 ( 1.3) | 0 ( 0.0) | | |
| Stage IA | 129 (26.7) | 129 (33.9) | 0 ( 0.0) | | |
| Stage IB | 128 (26.5) | 128 (33.7) | 0 ( 0.0) | | |
| Stage II | 1 ( 0.2) | 1 ( 0.3) | 0 ( 0.0) | | |
| Stage IIA | 50 (10.4) | 50 (13.2) | 0 ( 0.0) | | |
| Stage IIB | 67 (13.9) | 67 (17.6) | 0 ( 0.0) | | |
| Stage IIIA | 69 (14.3) | 0 ( 0.0) | 69 (67.0) | | |
| Stage IIIB | 10 ( 2.1) | 0 ( 0.0) | 10 ( 9.7) | | |
| Stage IV | 24 ( 5.0) | 0 ( 0.0) | 24 (23.3) | | |
| ajcc_pathologic_n (%) | | | | <0.001 | |
| N0 | 313 (64.8) | 297 (78.2) | 16 (15.5) | | |
| N1 | 90 (18.6) | 77 (20.3) | 13 (12.6) | | |
| N2 | 68 (14.1) | 0 ( 0.0) | 68 (66.0) | | |
| N3 | 2 ( 0.4) | 0 ( 0.0) | 2 ( 1.9) | | |
| NX | 10 ( 2.1) | 6 ( 1.6) | 4 ( 3.9) | | |
| prior_malignancy = yes (%) | 79 (16.4) | 65 (17.1) | 14 (13.6) | 0.481 | |
| primary_diagnosis (%) | | | | 0.621 | |
| Acinar cell carcinoma | 22 ( 4.6) | 16 ( 4.2) | 6 ( 5.8) | | |
| Adenocarcinoma with mixed subtypes | 103 (21.3) | 77 (20.3) | 26 (25.2) | | |
| Adenocarcinoma, NOS | 289 (59.8) | 230 (60.5) | 59 (57.3) | | |
| Bronchio-alveolar carcinoma, mucinous | 5 ( 1.0) | 5 ( 1.3) | 0 ( 0.0) | | |
| Bronchiolo-alveolar adenocarcinoma, NOS | 3 ( 0.6) | 3 ( 0.8) | 0 ( 0.0) | | |
| Bronchiolo-alveolar carcinoma, non-mucinous | 17 ( 3.5) | 13 ( 3.4) | 4 ( 3.9) | | |
| Clear cell adenocarcinoma, NOS | 2 ( 0.4) | 1 ( 0.3) | 1 ( 1.0) | | |
| Micropapillary carcinoma, NOS | 2 ( 0.4) | 2 ( 0.5) | 0 ( 0.0) | | |
| Mucinous adenocarcinoma | 13 ( 2.7) | 12 ( 3.2) | 1 ( 1.0) | | |
| Papillary adenocarcinoma, NOS | 21 ( 4.3) | 15 ( 3.9) | 6 ( 5.8) | | |
| Signet ring cell carcinoma | 1 ( 0.2) | 1 ( 0.3) | 0 ( 0.0) | | |
| Solid carcinoma, NOS | 5 ( 1.0) | 5 ( 1.3) | 0 ( 0.0) | | |
| time (mean (SD)) | 879.55 (873.37) | 929.04 (921.78) | 696.95 (636.03) | 0.017 | |
| race (%) | | | | 0.449 | |
| american indian or alaska native | 1 ( 0.2) | 1 ( 0.3) | 0 ( 0.0) | | |
| asian | 8 ( 1.7) | 6 ( 1.6) | 2 ( 1.9) | | |
| black or african american | 50 (10.4) | 41 (10.8) | 9 ( 8.7) | | |
| not reported | 44 ( 9.1) | 30 ( 7.9) | 14 (13.6) | | |
| white | 380 (78.7) | 302 (79.5) | 78 (75.7) | | |

| | Overall | Early Stage | Late Stage | p | test |
|---|---|---|---|---|---|
| MK2.Expression (mean (SD)) | 3.86 (1.05) | 3.91 (1.06) | 3.66 (0.99) | 0.032 | |

```r
# Table 1 where we stratify by MK2 'hi' vs. 'low'
table1b <- CreateTableOne(listVars, clinical, catVars, strata = c("MK2_Expression_topQ_t1"),
    addOverall = TRUE)
kable(print(table1b))
```

```
##                                                   Stratified by MK2_Expression_topQ_t1
##                                                    Overall
##   n                                                   483
##   age_at_diagnosis (mean (SD))                      65.77 (9.97)
##   gender = male (%)                                   219 (45.3)
##   smoking = 1 (%)                                     331 (68.5)
##   ajcc_pathologic_stage (%)
##      Stage I                                            5 ( 1.0)
##      Stage IA                                         129 (26.7)
##      Stage IB                                         128 (26.5)
##      Stage II                                           1 ( 0.2)
##      Stage IIA                                         50 (10.4)
##      Stage IIB                                         67 (13.9)
##      Stage IIIA                                        69 (14.3)
##      Stage IIIB                                        10 ( 2.1)
##      Stage IV                                          24 ( 5.0)
##   ajcc_pathologic_n (%)
##      N0                                               313 (64.8)
##      N1                                                90 (18.6)
##      N2                                                68 (14.1)
##      N3                                                 2 ( 0.4)
##      NX                                                10 ( 2.1)
##   prior_malignancy = yes (%)                           79 (16.4)
##   primary_diagnosis (%)
##      Acinar cell carcinoma                             22 ( 4.6)
##      Adenocarcinoma with mixed subtypes               103 (21.3)
##      Adenocarcinoma, NOS                              289 (59.8)
##      Bronchio-alveolar carcinoma, mucinous              5 ( 1.0)
##      Bronchiolo-alveolar adenocarcinoma, NOS            3 ( 0.6)
##      Bronchiolo-alveolar carcinoma, non-mucinous       17 ( 3.5)
##      Clear cell adenocarcinoma, NOS                     2 ( 0.4)
##      Micropapillary carcinoma, NOS                      2 ( 0.4)
##      Mucinous adenocarcinoma                           13 ( 2.7)
##      Papillary adenocarcinoma, NOS                     21 ( 4.3)
##      Signet ring cell carcinoma                         1 ( 0.2)
##      Solid carcinoma, NOS                               5 ( 1.0)
##   time (mean (SD))                                 879.55 (873.37)
##   race (%)
##      american indian or alaska native                   1 ( 0.2)
##      asian                                              8 ( 1.7)
##      black or african american                         50 (10.4)
##      not reported                                      44 ( 9.1)
##      white                                            380 (78.7)
##   MK2.Expression (mean (SD))                         3.86 (1.05)
##                                                   Stratified by MK2_Expression_topQ_t1
```

```
##                                               0
##   n                                              319
##   age_at_diagnosis (mean (SD))                 66.11 (9.96)
##   gender = male (%)                              153 (48.0)
##   smoking = 1 (%)                                213 (66.8)
##   ajcc_pathologic_stage (%)
##      Stage I                                       3 ( 0.9)
##      Stage IA                                     81 (25.4)
##      Stage IB                                     80 (25.1)
##      Stage II                                      0 ( 0.0)
##      Stage IIA                                    31 ( 9.7)
##      Stage IIB                                    47 (14.7)
##      Stage IIIA                                   53 (16.6)
##      Stage IIIB                                    6 ( 1.9)
##      Stage IV                                     18 ( 5.6)
##   ajcc_pathologic_n (%)
##      N0                                          201 (63.0)
##      N1                                           62 (19.4)
##      N2                                           50 (15.7)
##      N3                                            0 ( 0.0)
##      NX                                            6 ( 1.9)
##   prior_malignancy = yes (%)                      50 (15.7)
##   primary_diagnosis (%)
##      Acinar cell carcinoma                        15 ( 4.7)
##      Adenocarcinoma with mixed subtypes           74 (23.2)
##      Adenocarcinoma, NOS                         187 (58.6)
##      Bronchio-alveolar carcinoma, mucinous         5 ( 1.6)
##      Bronchiolo-alveolar adenocarcinoma, NOS       2 ( 0.6)
##      Bronchiolo-alveolar carcinoma, non-mucinous  10 ( 3.1)
##      Clear cell adenocarcinoma, NOS                2 ( 0.6)
##      Micropapillary carcinoma, NOS                 0 ( 0.0)
##      Mucinous adenocarcinoma                       9 ( 2.8)
##      Papillary adenocarcinoma, NOS                11 ( 3.4)
##      Signet ring cell carcinoma                    1 ( 0.3)
##      Solid carcinoma, NOS                          3 ( 0.9)
##   time (mean (SD))                            869.21 (953.82)
##   race (%)
##      american indian or alaska native              0 ( 0.0)
##      asian                                         6 ( 1.9)
##      black or african american                    35 (11.0)
##      not reported                                 36 (11.3)
##      white                                       242 (75.9)
##   MK2.Expression (mean (SD))                    3.27 (0.60)
##                                              Stratified by MK2_Expression_topQ_t1
##                                               1               p      test
##   n                                              164
##   age_at_diagnosis (mean (SD))                 65.12 (9.99)    0.307
##   gender = male (%)                               66 (40.2)    0.129
##   smoking = 1 (%)                                118 (72.0)    0.290
##   ajcc_pathologic_stage (%)                                    0.359
##      Stage I                                       2 ( 1.2)
##      Stage IA                                     48 (29.3)
##      Stage IB                                     48 (29.3)
##      Stage II                                      1 ( 0.6)
```

```
##       Stage IIA                                     19 (11.6)
##       Stage IIB                                     20 (12.2)
##       Stage IIIA                                    16 ( 9.8)
##       Stage IIIB                                     4 ( 2.4)
##       Stage IV                                       6 ( 3.7)
##    ajcc_pathologic_n (%)                                           0.162
##       N0                                           112 (68.3)
##       N1                                            28 (17.1)
##       N2                                            18 (11.0)
##       N3                                             2 ( 1.2)
##       NX                                             4 ( 2.4)
##    prior_malignancy = yes (%)                       29 (17.7)     0.663
##    primary_diagnosis (%)                                           0.356
##       Acinar cell carcinoma                          7 ( 4.3)
##       Adenocarcinoma with mixed subtypes            29 (17.7)
##       Adenocarcinoma, NOS                          102 (62.2)
##       Bronchio-alveolar carcinoma, mucinous          0 ( 0.0)
##       Bronchiolo-alveolar adenocarcinoma, NOS        1 ( 0.6)
##       Bronchiolo-alveolar carcinoma, non-mucinous    7 ( 4.3)
##       Clear cell adenocarcinoma, NOS                 0 ( 0.0)
##       Micropapillary carcinoma, NOS                  2 ( 1.2)
##       Mucinous adenocarcinoma                        4 ( 2.4)
##       Papillary adenocarcinoma, NOS                 10 ( 6.1)
##       Signet ring cell carcinoma                     0 ( 0.0)
##       Solid carcinoma, NOS                           2 ( 1.2)
##    time (mean (SD))                             899.66 (692.84)  0.717
##    race (%)                                                        0.078
##       american indian or alaska native               1 ( 0.6)
##       asian                                          2 ( 1.2)
##       black or african american                     15 ( 9.1)
##       not reported                                   8 ( 4.9)
##       white                                        138 (84.1)
##    MK2.Expression (mean (SD))                     5.01 (0.75)   <0.001
```

| | Overall | 0 | 1 | p | test |
|---|---|---|---|---|---|
| n | 483 | 319 | 164 | | |
| age_at_diagnosis (mean (SD)) | 65.77 (9.97) | 66.11 (9.96) | 65.12 (9.99) | 0.307 | |
| gender = male (%) | 219 (45.3) | 153 (48.0) | 66 (40.2) | 0.129 | |
| smoking = 1 (%) | 331 (68.5) | 213 (66.8) | 118 (72.0) | 0.290 | |
| ajcc_pathologic_stage (%) | | | | 0.359 | |
| Stage I | 5 ( 1.0) | 3 ( 0.9) | 2 ( 1.2) | | |
| Stage IA | 129 (26.7) | 81 (25.4) | 48 (29.3) | | |
| Stage IB | 128 (26.5) | 80 (25.1) | 48 (29.3) | | |
| Stage II | 1 ( 0.2) | 0 ( 0.0) | 1 ( 0.6) | | |
| Stage IIA | 50 (10.4) | 31 ( 9.7) | 19 (11.6) | | |
| Stage IIB | 67 (13.9) | 47 (14.7) | 20 (12.2) | | |
| Stage IIIA | 69 (14.3) | 53 (16.6) | 16 ( 9.8) | | |
| Stage IIIB | 10 ( 2.1) | 6 ( 1.9) | 4 ( 2.4) | | |
| Stage IV | 24 ( 5.0) | 18 ( 5.6) | 6 ( 3.7) | | |
| ajcc_pathologic_n (%) | | | | 0.162 | |
| N0 | 313 (64.8) | 201 (63.0) | 112 (68.3) | | |
| N1 | 90 (18.6) | 62 (19.4) | 28 (17.1) | | |
| N2 | 68 (14.1) | 50 (15.7) | 18 (11.0) | | |
| N3 | 2 ( 0.4) | 0 ( 0.0) | 2 ( 1.2) | | |

| | Overall | 0 | 1 | p | test |
|---|---|---|---|---|---|
| NX | 10 ( 2.1) | 6 ( 1.9) | 4 ( 2.4) | | |
| prior_malignancy = yes (%) | 79 (16.4) | 50 (15.7) | 29 (17.7) | 0.663 | |
| primary_diagnosis (%) | | | | 0.356 | |
| Acinar cell carcinoma | 22 ( 4.6) | 15 ( 4.7) | 7 ( 4.3) | | |
| Adenocarcinoma with mixed subtypes | 103 (21.3) | 74 (23.2) | 29 (17.7) | | |
| Adenocarcinoma, NOS | 289 (59.8) | 187 (58.6) | 102 (62.2) | | |
| Bronchio-alveolar carcinoma, mucinous | 5 ( 1.0) | 5 ( 1.6) | 0 ( 0.0) | | |
| Bronchiolo-alveolar adenocarcinoma, NOS | 3 ( 0.6) | 2 ( 0.6) | 1 ( 0.6) | | |
| Bronchiolo-alveolar carcinoma, non-mucinous | 17 ( 3.5) | 10 ( 3.1) | 7 ( 4.3) | | |
| Clear cell adenocarcinoma, NOS | 2 ( 0.4) | 2 ( 0.6) | 0 ( 0.0) | | |
| Micropapillary carcinoma, NOS | 2 ( 0.4) | 0 ( 0.0) | 2 ( 1.2) | | |
| Mucinous adenocarcinoma | 13 ( 2.7) | 9 ( 2.8) | 4 ( 2.4) | | |
| Papillary adenocarcinoma, NOS | 21 ( 4.3) | 11 ( 3.4) | 10 ( 6.1) | | |
| Signet ring cell carcinoma | 1 ( 0.2) | 1 ( 0.3) | 0 ( 0.0) | | |
| Solid carcinoma, NOS | 5 ( 1.0) | 3 ( 0.9) | 2 ( 1.2) | | |
| time (mean (SD)) | 879.55 (873.37) | 869.21 (953.82) | 899.66 (692.84) | 0.717 | |
| race (%) | | | | 0.078 | |
| american indian or alaska native | 1 ( 0.2) | 0 ( 0.0) | 1 ( 0.6) | | |
| asian | 8 ( 1.7) | 6 ( 1.9) | 2 ( 1.2) | | |
| black or african american | 50 (10.4) | 35 (11.0) | 15 ( 9.1) | | |
| not reported | 44 ( 9.1) | 36 (11.3) | 8 ( 4.9) | | |
| white | 380 (78.7) | 242 (75.9) | 138 (84.1) | | |
| MK2.Expression (mean (SD)) | 3.86 (1.05) | 3.27 (0.60) | 5.01 (0.75) | <0.001 | |

### Additional models and analysis

#### Continuous MK2 expression, no R censoring: Survival ~ expression (continuous) + clinical stage (stage 1, stage 2,. . . )

In this model, we use MK2 expression as a continuous variable - in other words, don't split into low vs. high (for that, see below). The 'MK.Expression' variable is the MK2 transcript level. Here, we're going to use the full follow-up time without any R censoring.

```
# COX PH model
coxph_model_all <- coxph(Surv(time, censor.2y) ~ gender + smoking + MK2.Expression,
    id = submitter_id, data = MK2_CoxPH_quantile[MK2_CoxPH_quantile$ajcc_pathologic_stage_combined ==
        "Early Stage", ])

# examine MK2 expression values
hist(clinical$MK2.Expression)
```

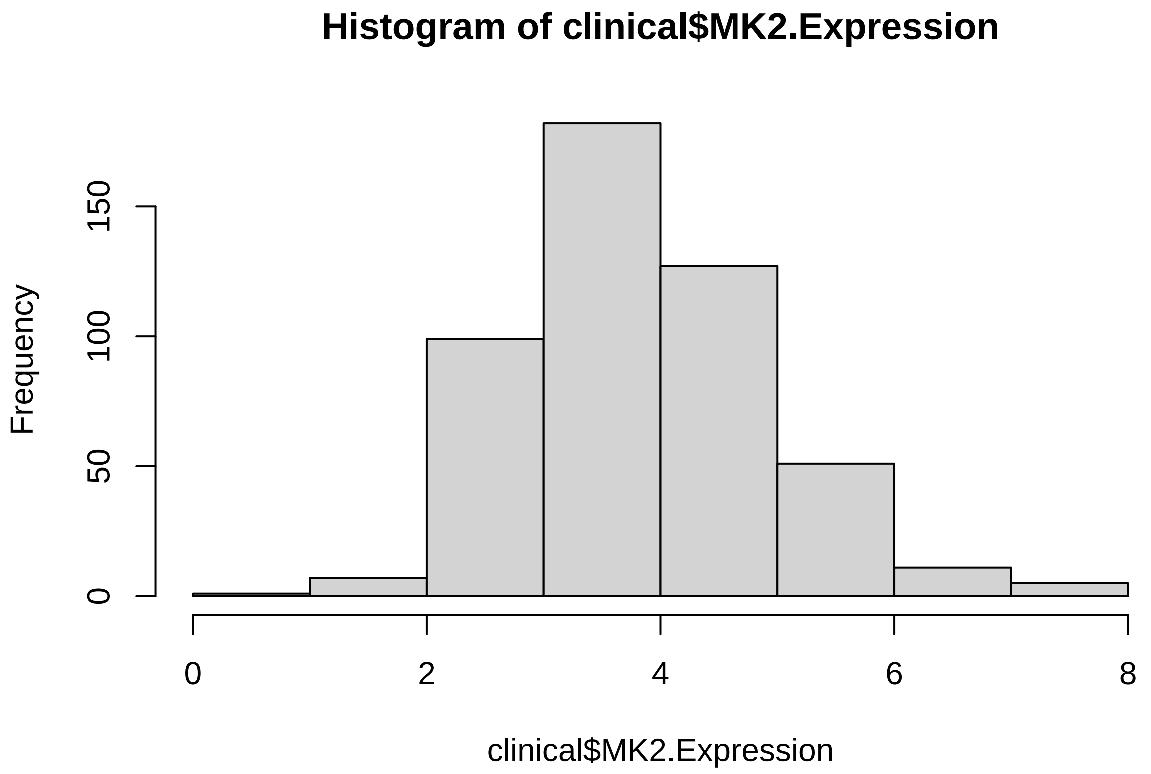

```
cox.zph(coxph_model_all)
```

```
##                 chisq df    p
## gender          2.021  1 0.16
## smoking         1.142  1 0.29
## MK2.Expression 0.142  1 0.71
## GLOBAL          3.305  3 0.35
```

```
plot(cox.zph(coxph_model_all)[3])
```

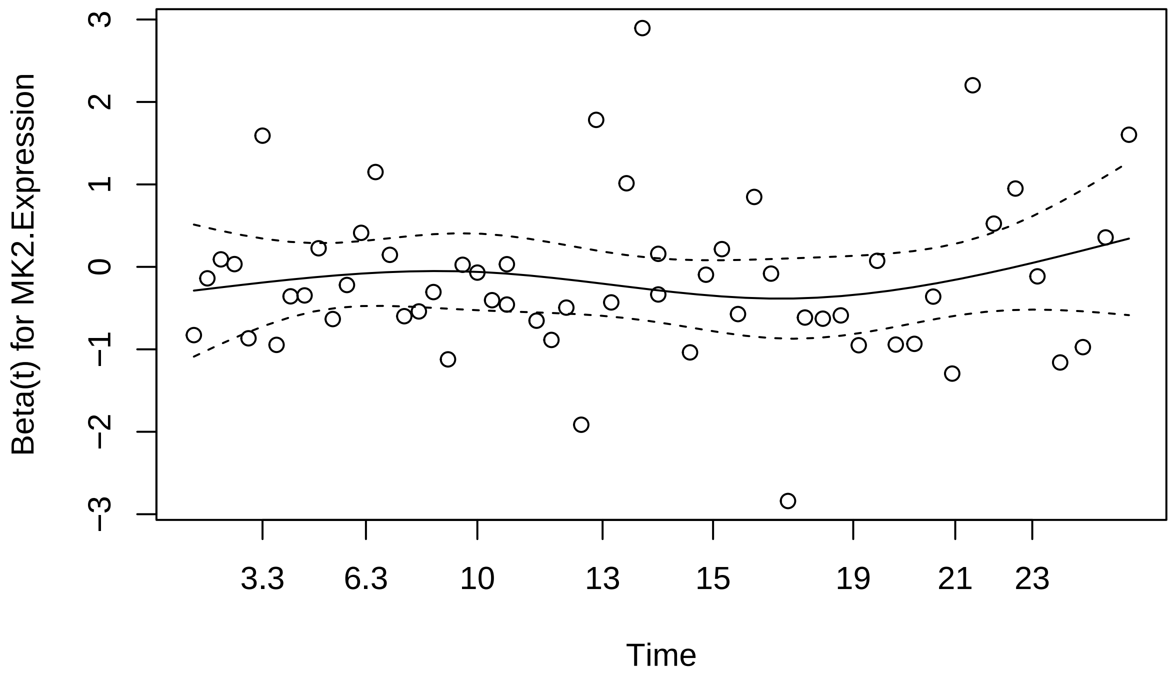

```r
summary(coxph_model_all)
```

```
## Call:
## coxph(formula = Surv(time, censor.2y) ~ gender + smoking + MK2.Expression,
##     data = MK2_CoxPH_quantile[MK2_CoxPH_quantile$ajcc_pathologic_stage_combined ==
##         "Early Stage", ], id = submitter_id)
##
##   n= 319, number of events= 58
##
##                   coef exp(coef) se(coef)      z Pr(>|z|)
## gendermale      0.54980   1.73290  0.26657  2.063   0.0392 *
## smoking         0.04049   1.04132  0.29428  0.138   0.8906
## MK2.Expression -0.16220   0.85027  0.12556 -1.292   0.1964
## ---
## Signif. codes:  0 '***' 0.001 '**' 0.01 '*' 0.05 '.' 0.1 ' ' 1
##
##                exp(coef) exp(-coef) lower .95 upper .95
## gendermale        1.7329     0.5771    1.0277     2.922
## smoking           1.0413     0.9603    0.5849     1.854
## MK2.Expression    0.8503     1.1761    0.6648     1.088
##
## Concordance= 0.605  (se = 0.037 )
## Likelihood ratio test= 6.57  on 3 df,   p=0.09
## Wald test            = 6.5  on 3 df,   p=0.09
## Score (logrank) test = 6.64  on 3 df,   p=0.08
```

##### Logistic regression for death at 1 year, stratified by specific stage

As shown above, LR using MK2 as a continuous variable but tumor stage dichotomized as early/late produced some interesting results. We wondered if we were "overlumping" stage by combining Stage I and II into "Early Stage". Here we split stages into 1-4. Of course, now the problem was low numbers. These models did not yield any additional information, so were not considered further.

```r
# LR stratified by individual stage
summary(glm(death_at_1year ~ MK2.Expression, data = MK2_CoxPH_quantile[MK2_CoxPH_quantile$ajcc_patholog
    "Stage 1", ], family = binomial(link = "logit")))
```

```
##
## Call:
## glm(formula = death_at_1year ~ MK2.Expression, family = binomial(link = "logit"),
##     data = MK2_CoxPH_quantile[MK2_CoxPH_quantile$ajcc_pathologic_stage_combined.2 ==
##         "Stage 1", ])
##
## Deviance Residuals:
##     Min       1Q   Median       3Q      Max
## -0.4456  -0.3181  -0.2899  -0.2583   2.6770
##
## Coefficients:
##                Estimate Std. Error z value Pr(>|z|)
## (Intercept)     -1.9805     1.2743  -1.554    0.120
## MK2.Expression  -0.2988     0.3375  -0.885    0.376
##
## (Dispersion parameter for binomial family taken to be 1)
##
##     Null deviance: 74.394  on 210  degrees of freedom
```

```
## Residual deviance: 73.576  on 209  degrees of freedom
## AIC: 77.576
##
## Number of Fisher Scoring iterations: 6
```

```
summary(glm(death_at_1year ~ MK2.Expression, data = MK2_CoxPH_quantile[MK2_CoxPH_quantile$ajcc_pathologi
    "Stage 2", ], family = binomial(link = "logit")))
```

```
##
## Call:
## glm(formula = death_at_1year ~ MK2.Expression, family = binomial(link = "logit"),
##     data = MK2_CoxPH_quantile[MK2_CoxPH_quantile$ajcc_pathologic_stage_combined.2 ==
##         "Stage 2", ])
##
## Deviance Residuals:
##     Min       1Q   Median       3Q      Max
## -0.9166  -0.6776  -0.5733  -0.4215   2.2349
##
## Coefficients:
##                Estimate Std. Error z value Pr(>|z|)
## (Intercept)      0.2439     1.0268   0.238   0.8123
## MK2.Expression  -0.4773     0.2768  -1.724   0.0846 .
## ---
## Signif. codes:  0 '***' 0.001 '**' 0.01 '*' 0.05 '.' 0.1 ' ' 1
##
## (Dispersion parameter for binomial family taken to be 1)
##
##     Null deviance: 100.47  on 107  degrees of freedom
## Residual deviance:  97.09  on 106  degrees of freedom
## AIC: 101.09
##
## Number of Fisher Scoring iterations: 4
```

```
summary(glm(death_at_1year ~ MK2.Expression, data = MK2_CoxPH_quantile[MK2_CoxPH_quantile$ajcc_pathologi
    "Stage 3", ], family = binomial(link = "logit")))
```

```
##
## Call:
## glm(formula = death_at_1year ~ MK2.Expression, family = binomial(link = "logit"),
##     data = MK2_CoxPH_quantile[MK2_CoxPH_quantile$ajcc_pathologic_stage_combined.2 ==
##         "Stage 3", ])
##
## Deviance Residuals:
##     Min       1Q   Median       3Q      Max
## -1.0027  -0.8448  -0.7861   1.4666   1.7445
##
## Coefficients:
##                Estimate Std. Error z value Pr(>|z|)
## (Intercept)     0.05054    1.05892   0.048    0.962
## MK2.Expression -0.27542    0.29351  -0.938    0.348
##
## (Dispersion parameter for binomial family taken to be 1)
##
##     Null deviance: 79.905  on 66  degrees of freedom
## Residual deviance: 78.958  on 65  degrees of freedom
```

```
## AIC: 82.958
##
## Number of Fisher Scoring iterations: 4
```

```
summary(glm(death_at_1year ~ MK2.Expression, data = MK2_CoxPH_quantile[MK2_CoxPH_quantile$ajcc_patholog
    "Stage 4", ], family = binomial(link = "logit")))
```

```
##
## Call:
## glm(formula = death_at_1year ~ MK2.Expression, family = binomial(link = "logit"),
##     data = MK2_CoxPH_quantile[MK2_CoxPH_quantile$ajcc_pathologic_stage_combined.2 ==
##         "Stage 4", ])
##
## Deviance Residuals:
##     Min       1Q   Median       3Q      Max
## -1.1913  -0.6872  -0.5152  -0.4273   1.8134
##
## Coefficients:
##                Estimate Std. Error z value Pr(>|z|)
## (Intercept)     -3.8831     2.4529  -1.583    0.113
## MK2.Expression   0.7118     0.6355   1.120    0.263
##
## (Dispersion parameter for binomial family taken to be 1)
##
##     Null deviance: 19.557  on 18  degrees of freedom
## Residual deviance: 18.216  on 17  degrees of freedom
## AIC: 22.216
##
## Number of Fisher Scoring iterations: 4
```

```
summary(glm(death_at_1year ~ MK2_Expression_topQ, data = MK2_CoxPH_quantile[MK2_CoxPH_quantile$ajcc_pat
    "Stage 1", ], family = binomial(link = "logit")))
```

```
##
## Call:
## glm(formula = death_at_1year ~ MK2_Expression_topQ, family = binomial(link = "logit"),
##     data = MK2_CoxPH_quantile[MK2_CoxPH_quantile$ajcc_pathologic_stage_combined.2 ==
##         "Stage 1", ])
##
## Deviance Residuals:
##     Min       1Q   Median       3Q      Max
## -0.3550  -0.3550  -0.3550  -0.1586   2.9604
##
## Coefficients:
##                      Estimate Std. Error z value Pr(>|z|)
## (Intercept)           -2.7327     0.3649  -7.490 6.91e-14 ***
## MK2_Expression_topQ1  -1.6367     1.0704  -1.529    0.126
## ---
## Signif. codes:  0 '***' 0.001 '**' 0.01 '*' 0.05 '.' 0.1 ' ' 1
##
## (Dispersion parameter for binomial family taken to be 1)
##
##     Null deviance: 74.394  on 210  degrees of freedom
## Residual deviance: 70.985  on 209  degrees of freedom
## AIC: 74.985
```

```
##
## Number of Fisher Scoring iterations: 7
```

```
summary(glm(death_at_1year ~ MK2_Expression_topQ, data = MK2_CoxPH_quantile[MK2_CoxPH_quantile$ajcc_pat
    "Stage 2", ], family = binomial(link = "logit")))
```

```
##
## Call:
## glm(formula = death_at_1year ~ MK2_Expression_topQ, family = binomial(link = "logit"),
##     data = MK2_CoxPH_quantile[MK2_CoxPH_quantile$ajcc_pathologic_stage_combined.2 ==
##         "Stage 2", ])
##
## Deviance Residuals:
##     Min       1Q   Median       3Q      Max
## -0.7339  -0.7339  -0.7339  -0.3381   2.4043
##
## Coefficients:
##                      Estimate Std. Error z value Pr(>|z|)
## (Intercept)           -1.1741     0.2775  -4.231 2.33e-05 ***
## MK2_Expression_topQ1  -1.6591     0.7787  -2.131   0.0331 *
## ---
## Signif. codes:  0 '***' 0.001 '**' 0.01 '*' 0.05 '.' 0.1 ' ' 1
##
## (Dispersion parameter for binomial family taken to be 1)
##
##     Null deviance: 100.474  on 107  degrees of freedom
## Residual deviance:  94.152  on 106  degrees of freedom
## AIC: 98.152
##
## Number of Fisher Scoring iterations: 5
```

```
summary(glm(death_at_1year ~ MK2_Expression_topQ, data = MK2_CoxPH_quantile[MK2_CoxPH_quantile$ajcc_pat
    "Stage 3", ], family = binomial(link = "logit")))
```

```
##
## Call:
## glm(formula = death_at_1year ~ MK2_Expression_topQ, family = binomial(link = "logit"),
##     data = MK2_CoxPH_quantile[MK2_CoxPH_quantile$ajcc_pathologic_stage_combined.2 ==
##         "Stage 3", ])
##
## Deviance Residuals:
##     Min       1Q   Median       3Q      Max
## -0.8898  -0.8898  -0.8898   1.4953   2.0074
##
## Coefficients:
##                      Estimate Std. Error z value Pr(>|z|)
## (Intercept)           -0.7221     0.2956  -2.443   0.0146 *
## MK2_Expression_topQ1  -1.1497     0.8151  -1.411   0.1584
## ---
## Signif. codes:  0 '***' 0.001 '**' 0.01 '*' 0.05 '.' 0.1 ' ' 1
##
## (Dispersion parameter for binomial family taken to be 1)
##
##     Null deviance: 79.905  on 66  degrees of freedom
## Residual deviance: 77.506  on 65  degrees of freedom
```

```
## AIC: 81.506
##
## Number of Fisher Scoring iterations: 4
```

```
summary(glm(death_at_1year ~ MK2_Expression_topQ, data = MK2_CoxPH_quantile[MK2_CoxPH_quantile$ajcc_patl
    "Stage 4", ], family = binomial(link = "logit")))
```

```
##
## Call:
## glm(formula = death_at_1year ~ MK2_Expression_topQ, family = binomial(link = "logit"),
##     data = MK2_CoxPH_quantile[MK2_CoxPH_quantile$ajcc_pathologic_stage_combined.2 ==
##         "Stage 4", ])
##
## Deviance Residuals:
##     Min       1Q   Median       3Q      Max
## -0.7585  -0.6681  -0.6681  -0.6681   1.7941
##
## Coefficients:
##                      Estimate Std. Error z value Pr(>|z|)
## (Intercept)           -1.3863     0.6455  -2.148   0.0317 *
## MK2_Expression_topQ1   0.2877     1.3229   0.217   0.8278
## ---
## Signif. codes:  0 '***' 0.001 '**' 0.01 '*' 0.05 '.' 0.1 ' ' 1
##
## (Dispersion parameter for binomial family taken to be 1)
##
##     Null deviance: 19.557  on 18  degrees of freedom
## Residual deviance: 19.511  on 17  degrees of freedom
## AIC: 23.511
##
## Number of Fisher Scoring iterations: 4
```

```
summary(glm(death_at_1year ~ MK2_Expression_topQ, data = MK2_CoxPH_quantile[MK2_CoxPH_quantile$ajcc_patl
    "Stage 1", ], family = binomial(link = "logit")))
```

```
##
## Call:
## glm(formula = death_at_1year ~ MK2_Expression_topQ, family = binomial(link = "logit"),
##     data = MK2_CoxPH_quantile[MK2_CoxPH_quantile$ajcc_pathologic_stage_combined.2 ==
##         "Stage 1", ])
##
## Deviance Residuals:
##     Min       1Q   Median       3Q      Max
## -0.3550  -0.3550  -0.3550  -0.1586   2.9604
##
## Coefficients:
##                      Estimate Std. Error z value Pr(>|z|)
## (Intercept)           -2.7327     0.3649  -7.490 6.91e-14 ***
## MK2_Expression_topQ1  -1.6367     1.0704  -1.529    0.126
## ---
## Signif. codes:  0 '***' 0.001 '**' 0.01 '*' 0.05 '.' 0.1 ' ' 1
##
## (Dispersion parameter for binomial family taken to be 1)
##
##     Null deviance: 74.394  on 210  degrees of freedom
```

```
## Residual deviance: 70.985  on 209  degrees of freedom
## AIC: 74.985
##
## Number of Fisher Scoring iterations: 7
```

```
summary(glm(death_at_1year ~ MK2_Expression_topQ, data = MK2_CoxPH_quantile[MK2_CoxPH_quantile$ajcc_patl
    "Stage 2", ], family = binomial(link = "logit")))
```

```
##
## Call:
## glm(formula = death_at_1year ~ MK2_Expression_topQ, family = binomial(link = "logit"),
##     data = MK2_CoxPH_quantile[MK2_CoxPH_quantile$ajcc_pathologic_stage_combined.2 ==
##         "Stage 2", ])
##
## Deviance Residuals:
##     Min       1Q   Median       3Q      Max
## -0.7339  -0.7339  -0.7339  -0.3381   2.4043
##
## Coefficients:
##                      Estimate Std. Error z value Pr(>|z|)
## (Intercept)           -1.1741     0.2775  -4.231 2.33e-05 ***
## MK2_Expression_topQ1  -1.6591     0.7787  -2.131   0.0331 *
## ---
## Signif. codes:  0 '***' 0.001 '**' 0.01 '*' 0.05 '.' 0.1 ' ' 1
##
## (Dispersion parameter for binomial family taken to be 1)
##
##     Null deviance: 100.474  on 107  degrees of freedom
## Residual deviance:  94.152  on 106  degrees of freedom
## AIC: 98.152
##
## Number of Fisher Scoring iterations: 5
```

```
summary(glm(death_at_1year ~ MK2_Expression_topQ, data = MK2_CoxPH_quantile[MK2_CoxPH_quantile$ajcc_patl
    "Stage 3", ], family = binomial(link = "logit")))
```

```
##
## Call:
## glm(formula = death_at_1year ~ MK2_Expression_topQ, family = binomial(link = "logit"),
##     data = MK2_CoxPH_quantile[MK2_CoxPH_quantile$ajcc_pathologic_stage_combined.2 ==
##         "Stage 3", ])
##
## Deviance Residuals:
##     Min       1Q   Median       3Q      Max
## -0.8898  -0.8898  -0.8898   1.4953   2.0074
##
## Coefficients:
##                      Estimate Std. Error z value Pr(>|z|)
## (Intercept)           -0.7221     0.2956  -2.443   0.0146 *
## MK2_Expression_topQ1  -1.1497     0.8151  -1.411   0.1584
## ---
## Signif. codes:  0 '***' 0.001 '**' 0.01 '*' 0.05 '.' 0.1 ' ' 1
##
## (Dispersion parameter for binomial family taken to be 1)
##
```

```
##     Null deviance: 79.905  on 66  degrees of freedom
## Residual deviance: 77.506  on 65  degrees of freedom
## AIC: 81.506
##
## Number of Fisher Scoring iterations: 4
```

```
summary(glm(death_at_1year ~ MK2_Expression_topQ, data = MK2_CoxPH_quantile[MK2_CoxPH_quantile$ajcc_patl
    "Stage 4", ], family = binomial(link = "logit")))
```

```
##
## Call:
## glm(formula = death_at_1year ~ MK2_Expression_topQ, family = binomial(link = "logit"),
##     data = MK2_CoxPH_quantile[MK2_CoxPH_quantile$ajcc_pathologic_stage_combined.2 ==
##         "Stage 4", ])
##
## Deviance Residuals:
##     Min       1Q   Median       3Q      Max
## -0.7585  -0.6681  -0.6681  -0.6681   1.7941
##
## Coefficients:
##                     Estimate Std. Error z value Pr(>|z|)
## (Intercept)          -1.3863     0.6455  -2.148   0.0317 *
## MK2_Expression_topQ1   0.2877     1.3229   0.217   0.8278
## ---
## Signif. codes:  0 '***' 0.001 '**' 0.01 '*' 0.05 '.' 0.1 ' ' 1
##
## (Dispersion parameter for binomial family taken to be 1)
##
##     Null deviance: 19.557  on 18  degrees of freedom
## Residual deviance: 19.511  on 17  degrees of freedom
## AIC: 23.511
##
## Number of Fisher Scoring iterations: 4
```

##### Jackknife-type analysis for effects due to long period of patient recruitment

It appears that patients were recruited across many years. Chemo regimens change with time. To Figure out whether this plays a role, we re-estimate the odds ratio for high MK2 levels in the multivariate Linear Regression Model (Model 2) multiple times, each time removing only the patients recruited in a particular year

```
MK2_tvary <- MK2_CoxPH_quantile
years <- as.data.frame(table(as.factor(as.character(clinical$year_of_diagnosis))))
MK2_tvary$year_of_diagnosis <- as.factor(as.character(MK2_tvary$year_of_diagnosis))

returnPointEst <- function(x) {
    modelresult <- glm(death_at_1year ~ MK2_Expression_topQ + smoking +
        age_at_diagnosis + gender, data = MK2_CoxPH_quantile[MK2_CoxPH_quantile$ajcc_pathologic_stage_c
        "Early Stage", ], family = binomial(link = "logit"))
    return(exp(modelresult$coefficients[1]))
}

years$estimate <- lapply(as.character(years$Var1), returnPointEst)
years$estimate <- lapply(years$estimate, function(x) return(x[[1]]))
```

```
returnPointEstlci <- function(x) {
    modelresult <- glm(death_at_1year ~ MK2_Expression_topQ + smoking +
        age_at_diagnosis + gender, data = MK2_CoxPH_quantile[MK2_CoxPH_quantile$ajcc_pathologic_stage_c
        "Early Stage", ], family = binomial(link = "logit"))
    return(exp(confint.lm(modelresult)[2]))
}

returnPointEstuci <- function(x) {
    modelresult <- glm(death_at_1year ~ MK2_Expression_topQ + smoking +
        age_at_diagnosis + gender, data = MK2_CoxPH_quantile[MK2_CoxPH_quantile$ajcc_pathologic_stage_c
        "Early Stage", ], family = binomial(link = "logit"))
    return(exp(confint.lm(modelresult)[7]))
}

years$estimate_lci <- lapply(as.character(years$Var1), returnPointEstlci)
years$estimate_uci <- lapply(as.character(years$Var1), returnPointEstuci)

years$estimate <- as.numeric(years$estimate)
years$estimate_lci <- as.numeric(years$estimate_lci)
years$estimate_uci <- as.numeric(years$estimate_uci)

ggplot(years, aes(x = Var1, y = estimate)) + geom_point() + geom_errorbar(aes(ymin = estimate_lci,
    ymax = estimate_uci), width = 0) + theme_bw() + ylim(0, 1.2) + geom_hline(yintercept = 1) +
    ylab("OR for death at one year as a function of High MK2 level") +
    xlab("Patient Recruitment Year that was excluded from model") + theme(axis.text.x = element_text(an
    hjust = 1))
```

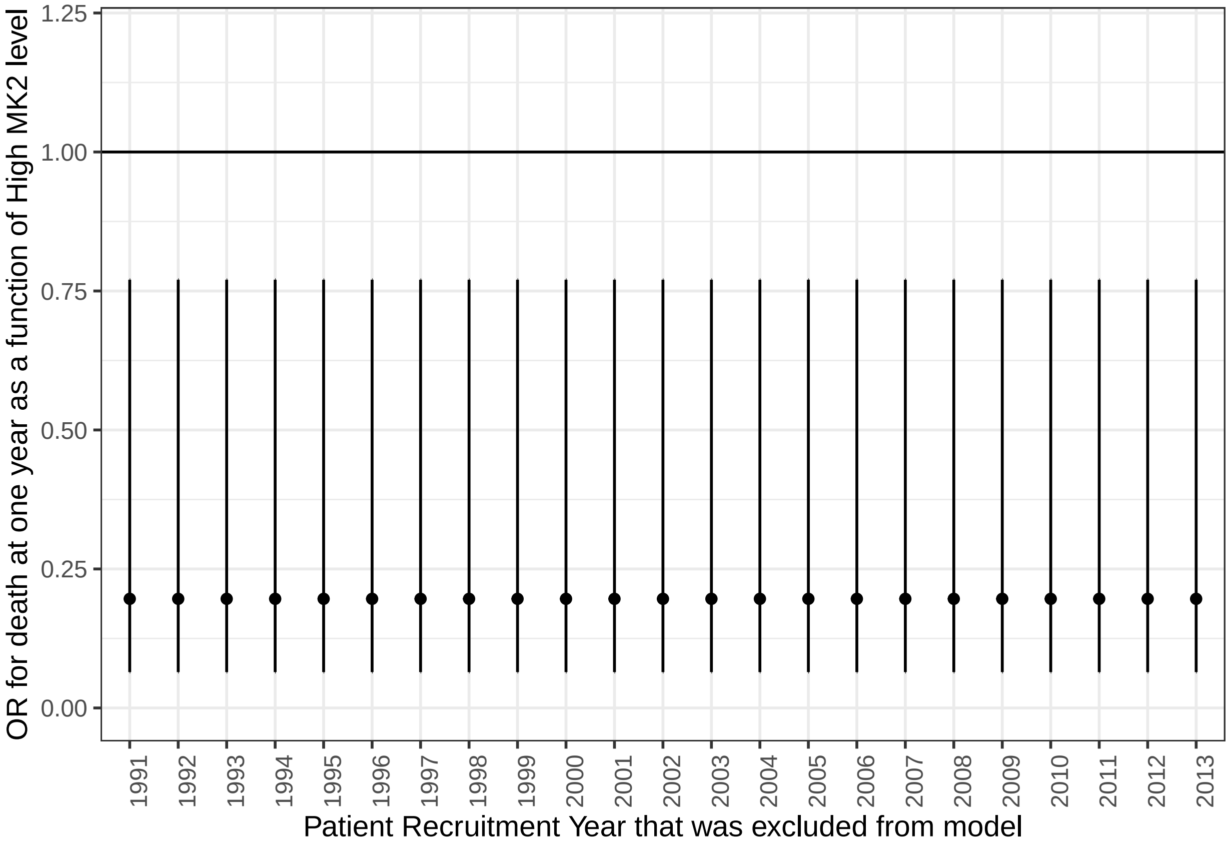

### Normal vs. tumor tissue

One of the benefits of downloading data from the TCGA is that it includes the "11" barcode, which is normal tissue

```
expr2_normalcontrols$Patient <- substring(expr2_normalcontrols$Patient,
    1, 12)
expr2_normalcontrols$sample.type = "Normal Tissue"
expr2_t <- expr3
expr2_t$sample.type = "Tumor"

exp2_paired <- left_join(expr2_normalcontrols, expr2_t, by = c("Patient"))
exp2_paired <- exp2_paired[!is.na(exp2_paired$MK2.Expression.Manual.y),
    ]
colnames(exp2_paired) <- c("MK2.Expression.Normal", "Stage.1", "Patient",
    "sample.type.1", "MK2.Expression.Tumor", "Stage.2", "sample.type.2")

# These data are plotted elsewhere. (See graphpad file)

# look at FC in MK2 as a predictor of survival

exp2_paired$MK2_diff <- exp2_paired$MK2.Expression.Tumor - exp2_paired$MK2.Expression.Normal
exp2_paired$MK2_FC <- exp2_paired$MK2.Expression.Tumor/exp2_paired$MK2.Expression.Normal
exp2_paired$Patient <- substring(exp2_paired$Patient, 1, 12)

exp2_paired <- left_join(exp2_paired, clinical, by = c(Patient = "submitter_id"))

ggplot(exp2_paired, aes(x = Stage.1, y = MK2_diff, fill = vital_status)) +
```

```
    geom_boxplot() + stat_compare_means()
```

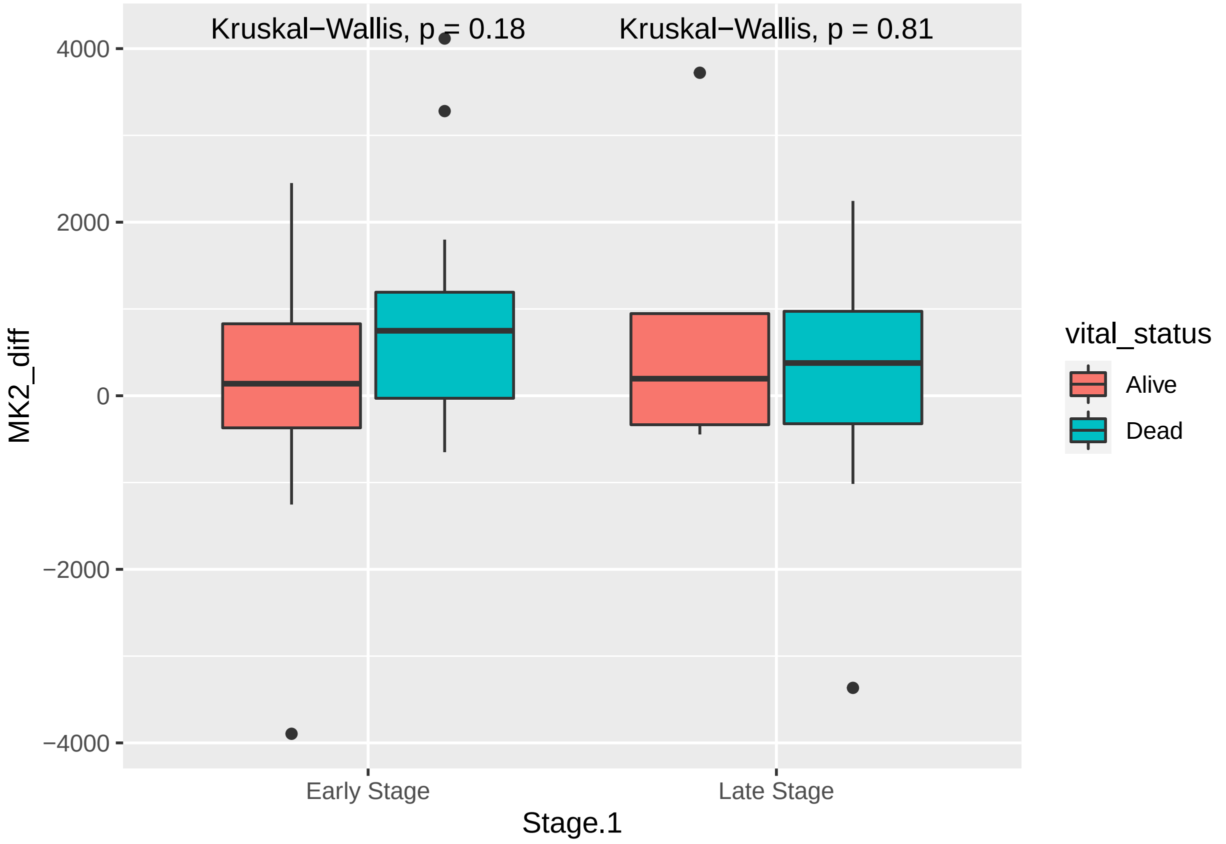

```
ggplot(exp2_paired, aes(x = Stage.1, y = MK2.Expression.Normal, fill = vital_status)) +
    geom_boxplot() + stat_compare_means()
```

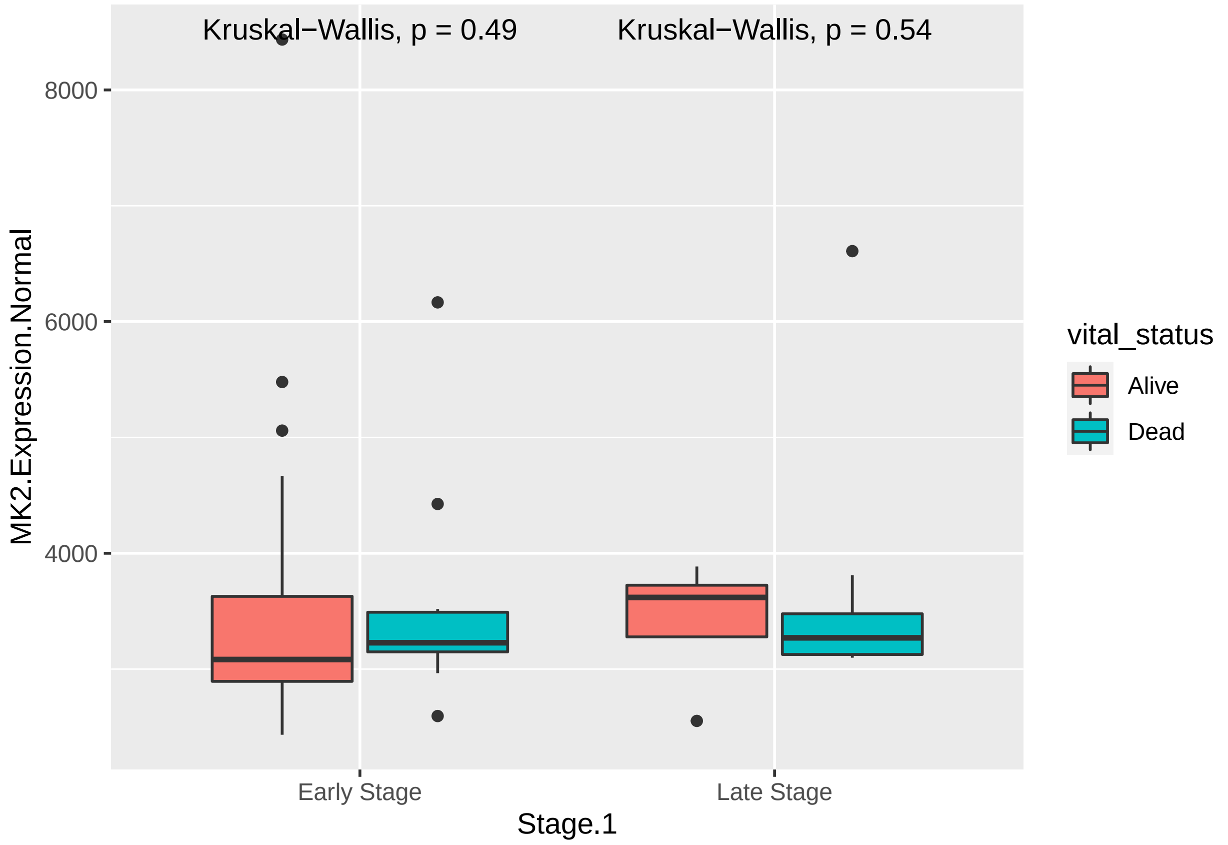

```
ggplot(exp2_paired, aes(x = Stage.1, y = MK2.Expression.Tumor, fill = vital_status)) +
    geom_boxplot() + stat_compare_means()
```

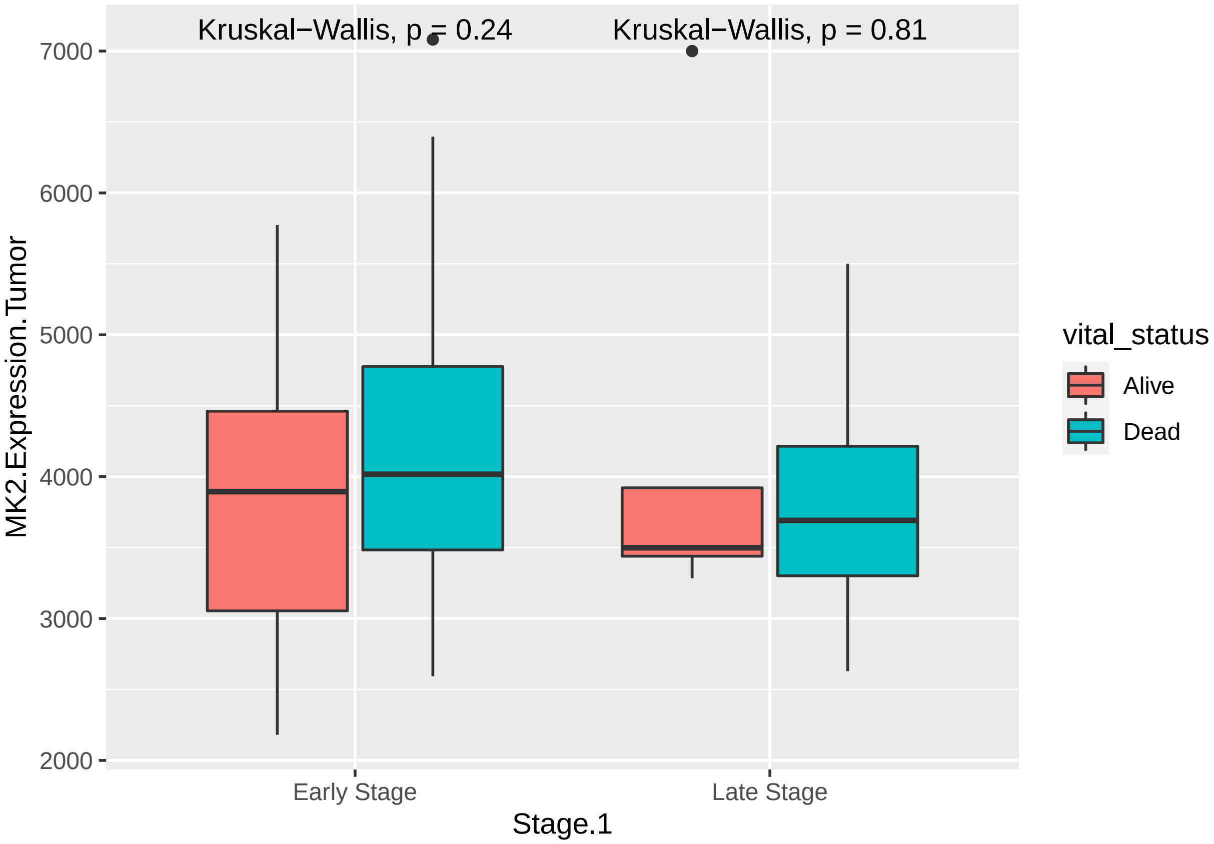

```
summary(coxph(Surv(time, censor) ~ MK2_FC, id = Patient, data = exp2_paired))
```

```
## Call:
## coxph(formula = Surv(time, censor) ~ MK2_FC, data = exp2_paired,
##     id = Patient)
##
##   n= 56, number of events= 26
##
##           coef exp(coef) se(coef)     z Pr(>|z|)
## MK2_FC 0.07524   1.07814  0.52357 0.144    0.886
##
##        exp(coef) exp(-coef) lower .95 upper .95
## MK2_FC     1.078     0.9275    0.3864     3.008
##
## Concordance= 0.461  (se = 0.057 )
## Likelihood ratio test= 0.02  on 1 df,   p=0.9
## Wald test            = 0.02  on 1 df,   p=0.9
## Score (logrank) test = 0.02  on 1 df,   p=0.9
```

#### Hsp27 and MK2

Here, we explore correlations between Hsp27 and MK2

```
HSP27Exp <- read.csv("../rawdata/LUAD.Hsp27Exp.csv")

HSP27Exp <- left_join(HSP27Exp, expr, by = c("Patient"))
library(ggpubr)
```

```r
ggplot(HSP27Exp, aes(x = log(Expression), y = log(MK2.Expression))) + geom_point() +
    geom_smooth(method = "lm") + theme_bw() + stat_cor()
```

```
## `geom_smooth()` using formula 'y ~ x'
## Warning: Removed 1 rows containing non-finite values (stat_smooth).
## Warning: Removed 1 rows containing non-finite values (stat_cor).
## Warning: Removed 1 rows containing missing values (geom_point).
```

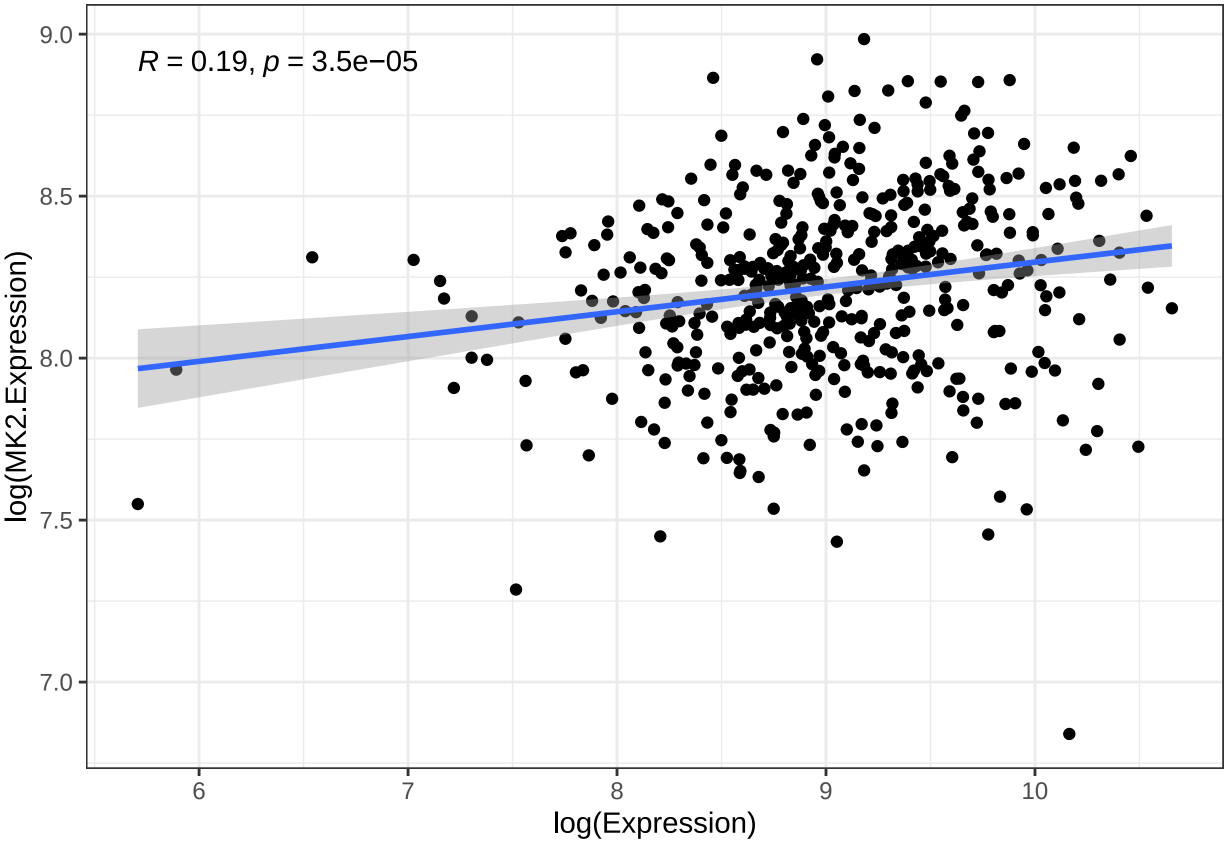

### System Info

```r
sessionInfo()
```

```
## R version 4.1.0 (2021-05-18)
## Platform: x86_64-pc-linux-gnu (64-bit)
## Running under: Ubuntu 20.04.2 LTS
##
## Matrix products: default
## BLAS:   /usr/lib/x86_64-linux-gnu/blas/libblas.so.3.9.0
## LAPACK: /usr/lib/x86_64-linux-gnu/lapack/liblapack.so.3.9.0
##
## locale:
##  [1] LC_CTYPE=en_US.UTF-8       LC_NUMERIC=C
##  [3] LC_TIME=en_US.UTF-8        LC_COLLATE=en_US.UTF-8
##  [5] LC_MONETARY=en_US.UTF-8    LC_MESSAGES=en_US.UTF-8
##  [7] LC_PAPER=en_US.UTF-8       LC_NAME=C
##  [9] LC_ADDRESS=C               LC_TELEPHONE=C
## [11] LC_MEASUREMENT=en_US.UTF-8 LC_IDENTIFICATION=C
```

```
##
## attached base packages:
## [1] parallel  stats4    stats     graphics  grDevices utils     datasets
## [8] methods   base
##
## other attached packages:
##  [1] tableone_0.13.0             ggthemes_4.2.4
##  [3] effects_4.2-0               carData_3.0-4
##  [5] ggsci_2.9                   ggpmisc_0.4.0
##  [7] ggpp_0.4.0                  tidyr_1.1.3
##  [9] survminer_0.4.9             ggpubr_0.4.0
## [11] ggplot2_3.3.5               survival_3.2-11
## [13] TCGAbiolinks_2.20.0         DT_0.18
## [15] dplyr_1.0.7                 SummarizedExperiment_1.22.0
## [17] Biobase_2.52.0              GenomicRanges_1.44.0
## [19] GenomeInfoDb_1.28.1         IRanges_2.26.0
## [21] S4Vectors_0.30.0            BiocGenerics_0.38.0
## [23] MatrixGenerics_1.4.3        matrixStats_0.60.1
## [25] knitr_1.33
##
## loaded via a namespace (and not attached):
##   [1] readxl_1.3.1                backports_1.2.1
##   [3] BiocFileCache_2.0.0         plyr_1.8.6
##   [5] splines_4.1.0               digest_0.6.27
##   [7] htmltools_0.5.2             fansi_0.5.0
##   [9] magrittr_2.0.1              memoise_2.0.0
##  [11] openxlsx_4.2.4              Biostrings_2.60.1
##  [13] readr_1.4.0                 R.utils_2.10.1
##  [15] prettyunits_1.1.1           colorspace_2.0-2
##  [17] blob_1.2.2                  rvest_1.0.0
##  [19] rappdirs_0.3.3              mitools_2.4
##  [21] haven_2.4.1                 xfun_0.25
##  [23] crayon_1.4.1                RCurl_1.98-1.3
##  [25] jsonlite_1.7.2              lme4_1.1-27.1
##  [27] zoo_1.8-9                   glue_1.4.2
##  [29] gtable_0.3.0                zlibbioc_1.38.0
##  [31] XVector_0.32.0              MatrixModels_0.5-0
##  [33] DelayedArray_0.18.0         car_3.0-11
##  [35] abind_1.4-5                 SparseM_1.81
##  [37] scales_1.1.1                DBI_1.1.1
##  [39] rstatix_0.7.0               Rcpp_1.0.7
##  [41] xtable_1.8-4                progress_1.2.2
##  [43] proxy_0.4-26                foreign_0.8-81
##  [45] bit_4.0.4                   km.ci_0.5-2
##  [47] survey_4.0                  htmlwidgets_1.5.3
##  [49] httr_1.4.2                  ellipsis_0.3.2
##  [51] pkgconfig_2.0.3             XML_3.99-0.7
##  [53] R.methodsS3_1.8.1           farver_2.1.0
##  [55] nnet_7.3-16                 dbplyr_2.1.1
##  [57] utf8_1.2.2                  tidyselect_1.1.1
##  [59] labeling_0.4.2              rlang_0.4.11
##  [61] polynom_1.4-0               AnnotationDbi_1.54.1
##  [63] munsell_0.5.0               cellranger_1.1.0
##  [65] tools_4.1.0                 cachem_1.0.6
```

```
##  [67] downloader_0.4             generics_0.1.0
##  [69] RSQLite_2.2.8              broom_0.7.8
##  [71] evaluate_0.14              stringr_1.4.0
##  [73] fastmap_1.1.0              yaml_2.2.1
##  [75] bit64_4.0.5                zip_2.2.0
##  [77] survMisc_0.5.5             purrr_0.3.4
##  [79] KEGGREST_1.32.0            nlme_3.1-152
##  [81] quantreg_5.86              formatR_1.11
##  [83] R.oo_1.24.0                xml2_1.3.2
##  [85] biomaRt_2.48.2             compiler_4.1.0
##  [87] filelock_1.0.2             curl_4.3.2
##  [89] png_0.1-7                  e1071_1.7-7
##  [91] ggsignif_0.6.2             tibble_3.1.4
##  [93] stringi_1.7.4              highr_0.9
##  [95] TCGAbiolinksGUI.data_1.12.0 forcats_0.5.1
##  [97] lattice_0.20-44            Matrix_1.3-4
##  [99] nloptr_1.2.2.2             KMsurv_0.1-5
## [101] vctrs_0.3.8                pillar_1.6.2
## [103] lifecycle_1.0.0            estimability_1.3
## [105] data.table_1.14.0          bitops_1.0-7
## [107] insight_0.14.2             conquer_1.0.2
## [109] R6_2.5.1                   gridExtra_2.3
## [111] rio_0.5.27                 boot_1.3-28
## [113] MASS_7.3-54                assertthat_0.2.1
## [115] withr_2.4.2                GenomeInfoDbData_1.2.6
## [117] mgcv_1.8-36                hms_1.1.0
## [119] labelled_2.8.0             grid_4.1.0
## [121] class_7.3-19               minqa_1.2.4
## [123] rmarkdown_2.11
```

### MK2 Validation Analysis

Karthik Suresh

9/30/2020; updated: 11/18/2021

### Data Import and cleaning

The data for these analyses came from cBioPortal. We downloaded the clinical data as tracks from the website along with mRNA data. The mRNA data was available in 2 forms: log-normalized to other genes, or non-scaled - i.e. log normalized RSEM expression that is not normalized to other genes. We chose to use the latter (although on comparison, the difference in actual values between the two forms of mRNA data was minimal, and did not significantly change our model results)

```
library(knitr)
opts_chunk$set(tidy.opts=list(width.cutoff=70), tidy=TRUE)
```

```
knitr::opts_chunk$set(echo = TRUE)
library(data.table)
library(ggplot2)
library(ggpubr)
library(dplyr)
```

```
##
## Attaching package: 'dplyr'

## The following objects are masked from 'package:data.table':
##
##     between, first, last

## The following objects are masked from 'package:stats':
##
##     filter, lag

## The following objects are masked from 'package:base':
##
##     intersect, setdiff, setequal, union
```

```
library(survival)
library(survminer)
```

```
# This is MAPKAPK2 mRNA and survival data obtained from cBioBortal -
# Study: OncoSG, Nat Genetics 2020

MK2_valid <- read.table("../rawdata/validation.dataset.MK2.Exp.tsv", sep = "\t")

# these are normalized across the genome. We want the non-scaled data
MK2_valid_nn <- read.csv("../rawdata/MAPKAPK2_valid_mRNA_nonnorm.txt",
    sep = "\t")
MK2_valid <- MK2_valid[c(1:8, 14), ]
MK2_valid_t <- transpose(MK2_valid)
colnames(MK2_valid_t) <- MK2_valid_t[1, ]
MK2_valid_t <- MK2_valid_t[-c(1:2), ]
MK2_valid_t <- left_join(MK2_valid_t, MK2_valid_nn, by = c(track_name = "Patient.ID"))

# switch the expression over to the non-scaled version
MK2_valid_t$MAPKAPK2 <- MK2_valid_t$MAPKAPK2..mRNA.Expression..log.RNA.Seq.V2.RSEM.

MK2_valid_t$Stage.simp <- lapply(MK2_valid_t$Stage, function(x) if (x ==
    "I" | x == "II") return("Early Stage") else return("Late Stage"))
MK2_valid_t$Stage.simp <- as.factor(unlist(MK2_valid_t$Stage.simp))
MK2_valid_t$MAPKAPK2 <- as.numeric(MK2_valid_t$MAPKAPK2)

# MK2 expression levels
ggplot(MK2_valid_t, aes(x = Stage.simp, y = MAPKAPK2)) + geom_boxplot(outlier.shape = NA) +
    geom_point() + theme_bw() + stat_compare_means()
```

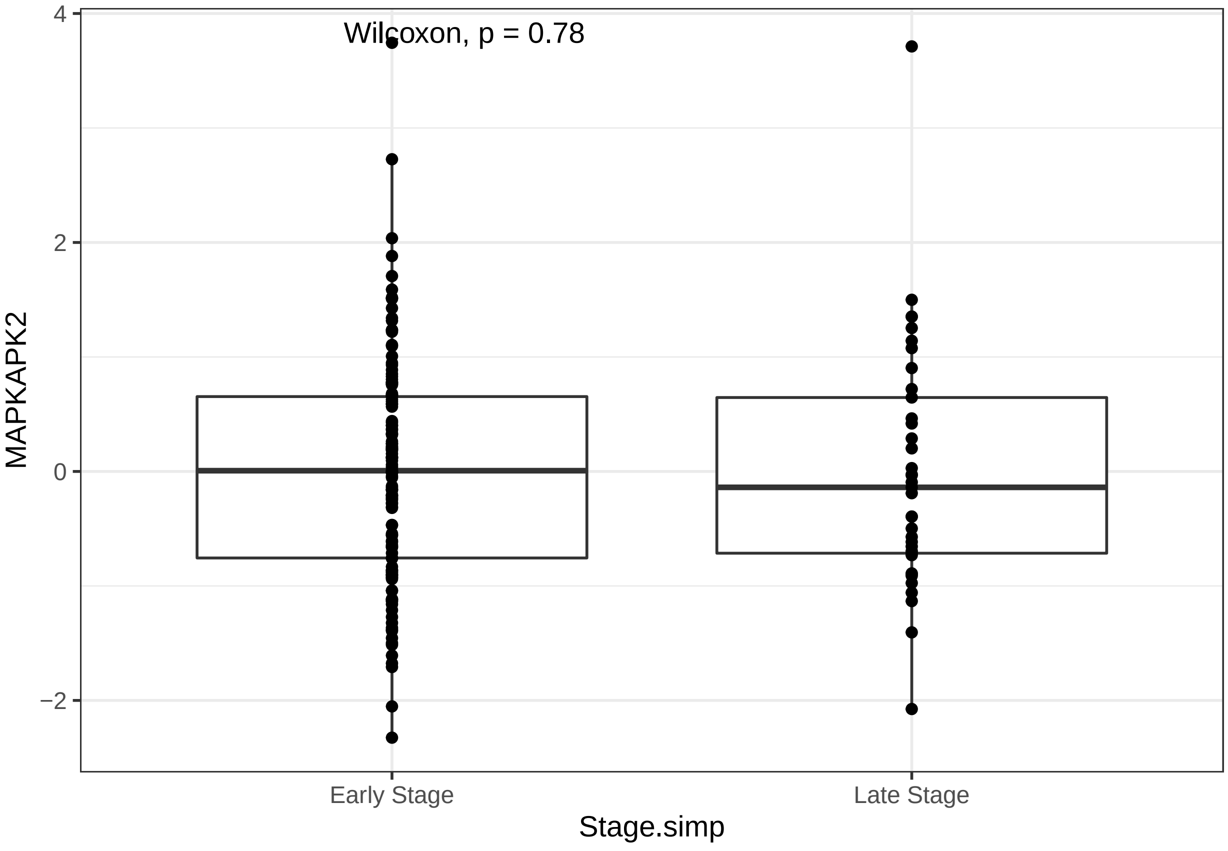

```
ggplot(MK2_valid_t, aes(x = MAPKAPK2, fill = Stage.simp)) + geom_density(adjust = 1.5,
    alpha = 0.5)
```

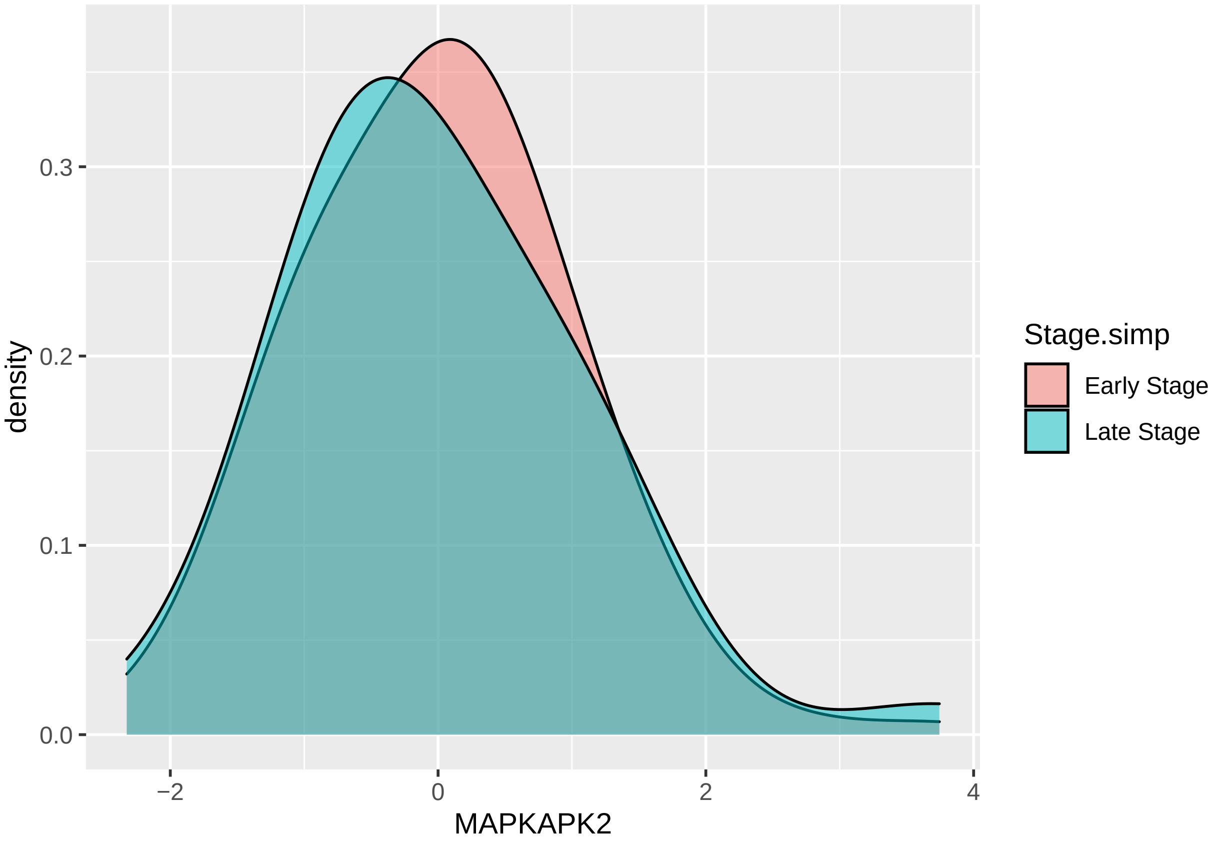

```
hist(MK2_valid_t$MAPKAPK2)
```

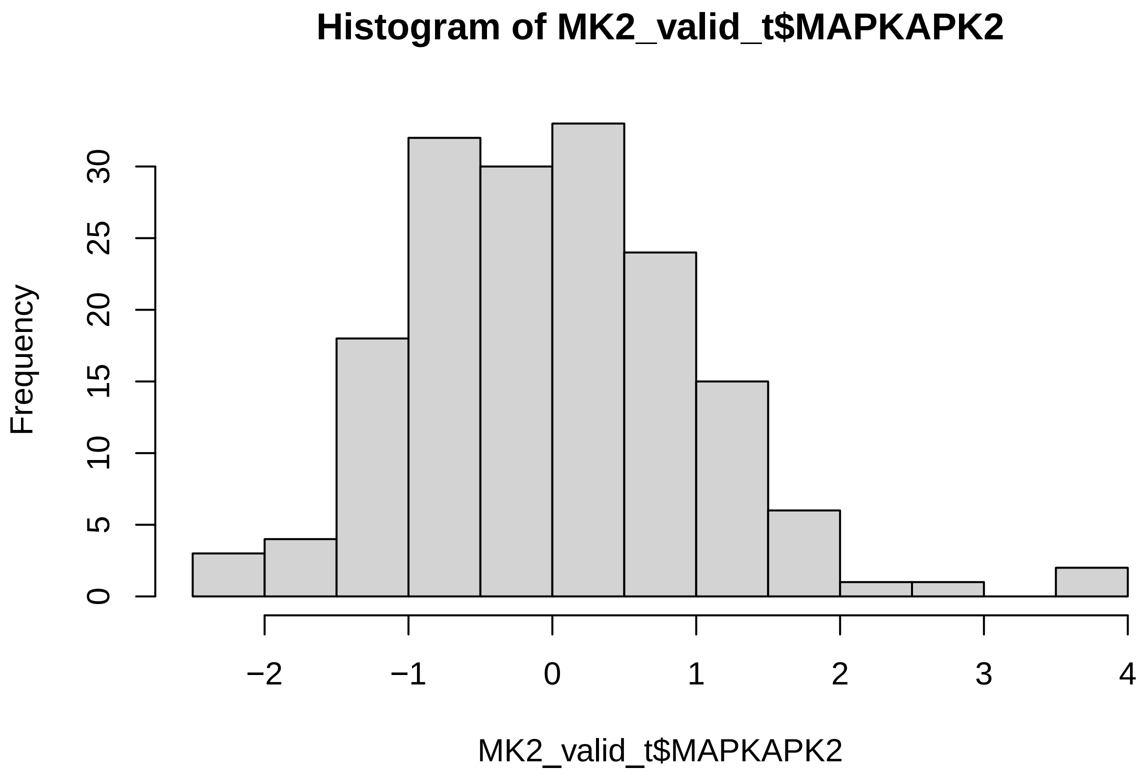

```r
# new variables for cox ph analysis
MK2_valid_t$censor <- lapply(MK2_valid_t$`Overall survival status`, function(x) if (x ==
    "0:LIVING") return(0) else return(1))
MK2_valid_t$time <- as.numeric(MK2_valid_t$`Overall survival months`)
MK2_valid_t$censor <- as.numeric(unlist(MK2_valid_t$censor))
MK2_valid_t$time <- as.numeric(MK2_valid_t$time)
MK2_valid_t$Sex <- as.factor(MK2_valid_t$Sex)
MK2_valid_t$Stage <- as.factor(MK2_valid_t$Stage)
MK2_valid_t$Smoking <- as.factor(MK2_valid_t$`Smoking status`)
MK2_valid_t$Age <- as.numeric(MK2_valid_t$Age)

# R censoring variable The problem is that there are very few events
# (deaths) at one year. So we can't R censor at 1 year like we did
# previously. So, we will try 2 years.
MK2_valid_t$censor_2yr <- mapply(function(dead, time) if (dead == 1 & time <
    25) return(1) else if (dead == 0 & time > 24) return(0) else if (dead ==
    0 & time < 25) return(0) else if (dead == 1 & time > 24) return(0),
    MK2_valid_t$censor, MK2_valid_t$time)
MK2_valid_t$censor_2yr <- as.numeric(unlist(MK2_valid_t$censor_2yr))

hist(MK2_valid_t$censor_2yr)
```

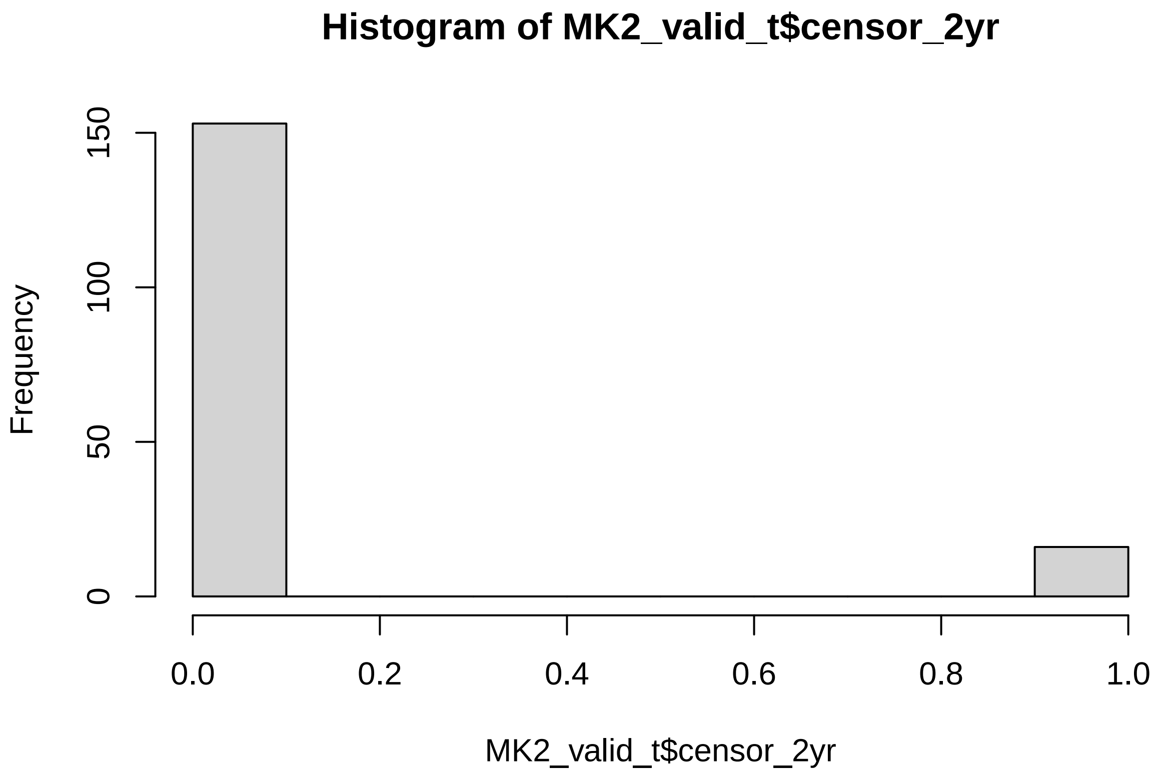

We will explore the data in 2 parts. In the first part - this is the part published in the paper - we really just look at the first two years (censoring at 2 years). This is in part to maintain symmetry with how we analyzed TCGA data.

Not shown here are our more extensive analyses sing the entire time period. The problem here is that MK2 expression (not unexpectedly) behaved in a time-varying fashion over this extended time period, and thus we used a variety of analyses to look at that relationship. If folks are interested in these analyses, they are available off the gitHub repo; you can look at the MK2analysis_validation.rmd file under ~/scripts/old.versions for these analyses.

#### Cox PH and Logistic Regression analysis with R censoring at 2 years

This will be done in two forms - using MK2 as a continuous variable, and again using top 1/3 vs. bottom 2/3 MK2 levels.

##### Cox PH with R censoring at 2 years, MK2 as a continuous variable

```
# MK2 survival analysis using R censoring at 1 year
summary(coxph(Surv(time, censor_2yr) ~ MAPKAPK2, id = Sample.ID, data = MK2_valid_t))
```

```
## Call:
## coxph(formula = Surv(time, censor_2yr) ~ MAPKAPK2, data = MK2_valid_t,
##     id = Sample.ID)
##
##   n= 169, number of events= 16
##
##              coef exp(coef) se(coef)      z Pr(>|z|)
## MAPKAPK2 -0.6548    0.5196   0.2796 -2.341   0.0192 *
```

```
## ---
## Signif. codes:  0 '***' 0.001 '**' 0.01 '*' 0.05 '.' 0.1 ' ' 1
##
##          exp(coef) exp(-coef) lower .95 upper .95
## MAPKAPK2    0.5196      1.925    0.3003    0.8988
##
## Concordance= 0.686  (se = 0.055 )
## Likelihood ratio test= 5.91  on 1 df,   p=0.02
## Wald test            = 5.48  on 1 df,   p=0.02
## Score (logrank) test = 5.38  on 1 df,   p=0.02
```

```
summary(coxph(Surv(time, censor_2yr) ~ MAPKAPK2 + Sex, id = Sample.ID,
    data = MK2_valid_t))
```

```
## Call:
## coxph(formula = Surv(time, censor_2yr) ~ MAPKAPK2 + Sex, data = MK2_valid_t,
##     id = Sample.ID)
##
##   n= 169, number of events= 16
##
##             coef exp(coef) se(coef)      z Pr(>|z|)
## MAPKAPK2 -0.7174    0.4880   0.2833 -2.532   0.0113 *
## SexMale   0.9137    2.4934   0.5204  1.756   0.0791 .
## ---
## Signif. codes:  0 '***' 0.001 '**' 0.01 '*' 0.05 '.' 0.1 ' ' 1
##
##          exp(coef) exp(-coef) lower .95 upper .95
## MAPKAPK2     0.488     2.0492    0.2801    0.8503
## SexMale      2.493     0.4011    0.8992    6.9145
##
## Concordance= 0.716  (se = 0.055 )
## Likelihood ratio test= 9.13  on 2 df,   p=0.01
## Wald test            = 8.55  on 2 df,   p=0.01
## Score (logrank) test = 8.26  on 2 df,   p=0.02
```

```
summary(coxph(Surv(time, censor_2yr) ~ MAPKAPK2 + Sex + Stage.simp, id = Sample.ID,
    data = MK2_valid_t))
```

```
## Call:
## coxph(formula = Surv(time, censor_2yr) ~ MAPKAPK2 + Sex + Stage.simp,
##     data = MK2_valid_t, id = Sample.ID)
##
##   n= 169, number of events= 16
##
##                        coef exp(coef) se(coef)      z Pr(>|z|)
## MAPKAPK2            -0.7861    0.4556   0.2984 -2.635  0.00842 **
## SexMale              0.9529    2.5933   0.5315  1.793  0.07299 .
## Stage.simpLate Stage  1.4174    4.1262   0.5032  2.816  0.00486 **
## ---
## Signif. codes:  0 '***' 0.001 '**' 0.01 '*' 0.05 '.' 0.1 ' ' 1
##
##                      exp(coef) exp(-coef) lower .95 upper .95
## MAPKAPK2                0.4556     2.1948    0.2539    0.8176
## SexMale                 2.5933     0.3856    0.9150    7.3497
## Stage.simpLate Stage    4.1262     0.2424    1.5388   11.0638
```

```
##
## Concordance= 0.795  (se = 0.044 )
## Likelihood ratio test= 16.5  on 3 df,   p=9e-04
## Wald test            = 14.81  on 3 df,   p=0.002
## Score (logrank) test = 16.91  on 3 df,   p=7e-04
```

```
summary(coxph(Surv(time, censor_2yr) ~ MAPKAPK2 + Age + Sex + Smoking +
    Stage.simp, id = Sample.ID, data = MK2_valid_t))
```

```
## Call:
## coxph(formula = Surv(time, censor_2yr) ~ MAPKAPK2 + Age + Sex +
##     Smoking + Stage.simp, data = MK2_valid_t, id = Sample.ID)
##
##   n= 169, number of events= 16
##
##                          coef exp(coef) se(coef)      z Pr(>|z|)
## MAPKAPK2             -0.90164   0.40590  0.30940 -2.914  0.00357 **
## Age                   0.03292   1.03346  0.02668  1.234  0.21722
## SexMale               0.08766   1.09162  0.61359  0.143  0.88640
## SmokingYes            1.63958   5.15300  0.65482  2.504  0.01228 *
## Stage.simpLate Stage  1.49883   4.47645  0.51553  2.907  0.00365 **
## ---
## Signif. codes:  0 '***' 0.001 '**' 0.01 '*' 0.05 '.' 0.1 ' ' 1
##
##                      exp(coef) exp(-coef) lower .95 upper .95
## MAPKAPK2                0.4059     2.4636    0.2213    0.7444
## Age                     1.0335     0.9676    0.9808    1.0889
## SexMale                 1.0916     0.9161    0.3279    3.6338
## SmokingYes              5.1530     0.1941    1.4278   18.5970
## Stage.simpLate Stage    4.4764     0.2234    1.6297   12.2958
##
## Concordance= 0.827  (se = 0.046 )
## Likelihood ratio test= 24.36  on 5 df,   p=2e-04
## Wald test            = 19.58  on 5 df,   p=0.001
## Score (logrank) test = 23.71  on 5 df,   p=2e-04
```

```
MK2_CoxPH_valid_RC <- coxph(Surv(time, censor_2yr) ~ MAPKAPK2 + Age + Sex +
    Smoking + Stage.simp, id = Sample.ID, data = MK2_valid_t)

ggplot(MK2_valid_t, aes(x = time, y = MAPKAPK2)) + geom_point() + geom_smooth(method = "lm") +
    theme_bw() + stat_cor()
```

```
## `geom_smooth()` using formula 'y ~ x'
```

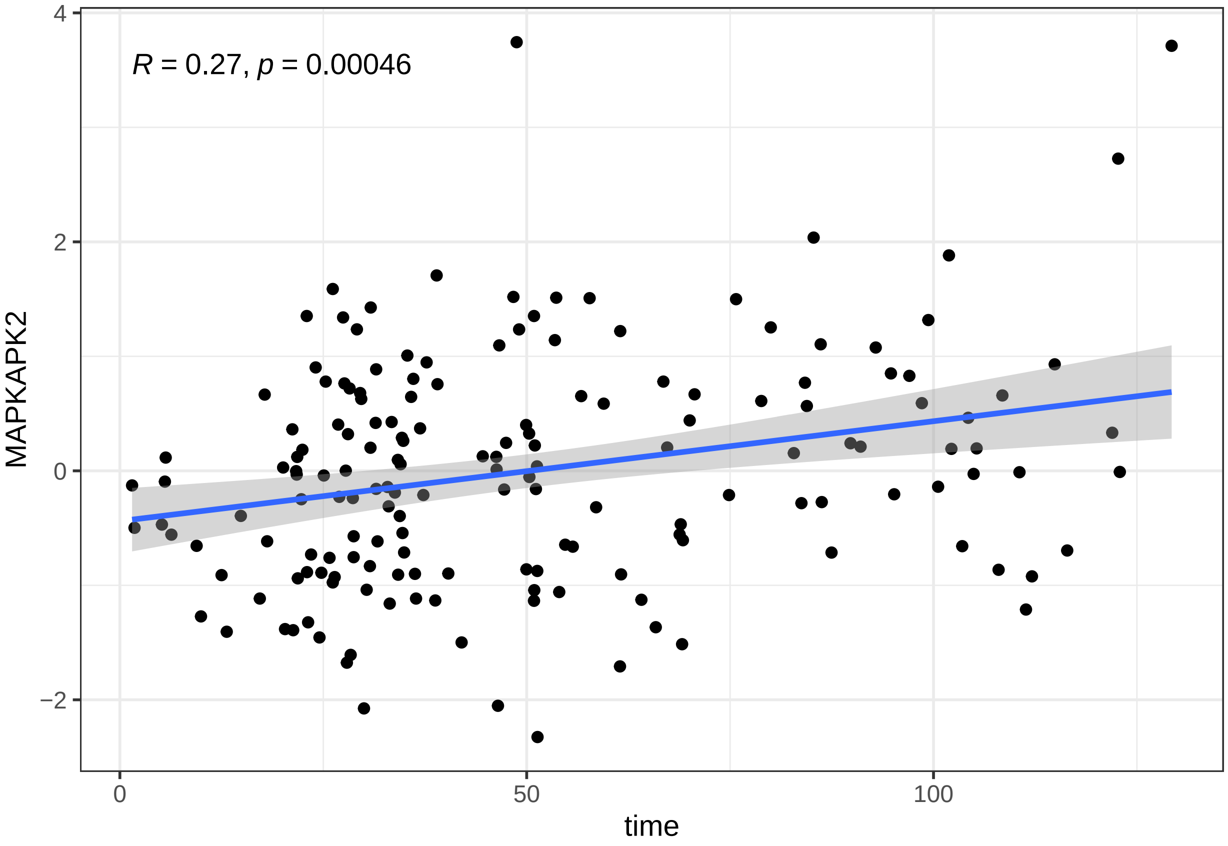

```r
cox.zph(MK2_CoxPH_valid_RC)
```

```
##             chisq df    p
## MAPKAPK2    0.117  1 0.73
## Age         0.624  1 0.43
## Sex         1.305  1 0.25
## Smoking     0.910  1 0.34
## Stage.simp 0.681  1 0.41
## GLOBAL      2.783  5 0.73
```

```r
plot(cox.zph(MK2_CoxPH_valid_RC)[1])
```

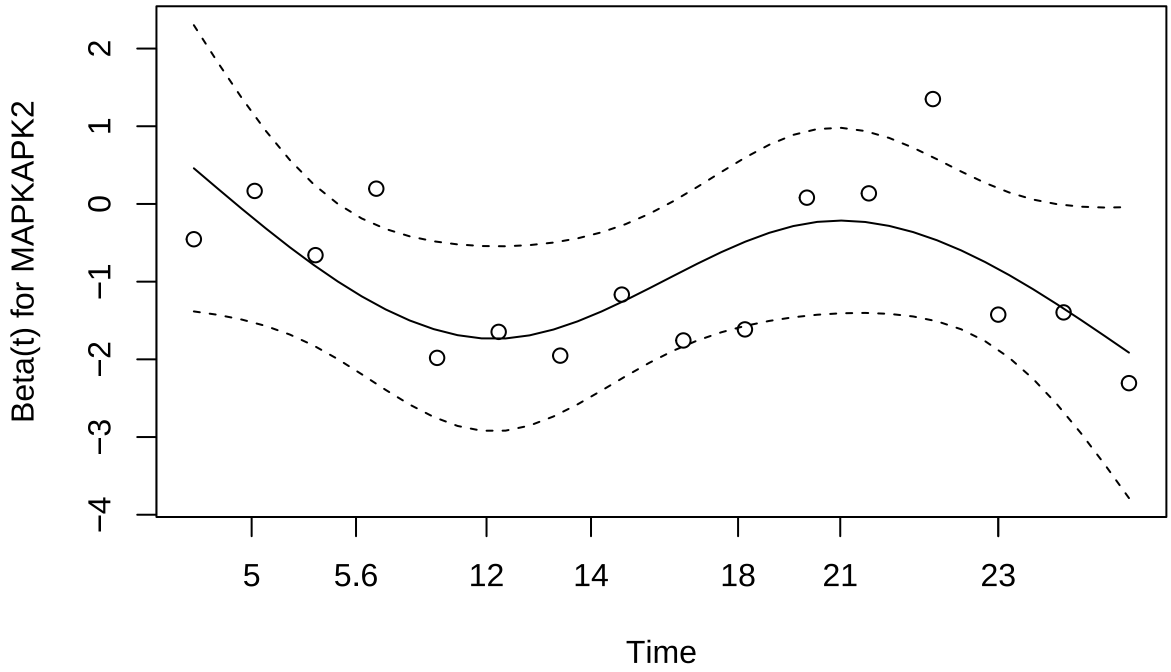

```
summary(MK2_CoxPH_valid_RC)
```

```
## Call:
## coxph(formula = Surv(time, censor_2yr) ~ MAPKAPK2 + Age + Sex +
##     Smoking + Stage.simp, data = MK2_valid_t, id = Sample.ID)
##
##   n= 169, number of events= 16
##
##                         coef exp(coef) se(coef)      z Pr(>|z|)
## MAPKAPK2             -0.90164   0.40590  0.30940 -2.914  0.00357 **
## Age                   0.03292   1.03346  0.02668  1.234  0.21722
## SexMale               0.08766   1.09162  0.61359  0.143  0.88640
## SmokingYes            1.63958   5.15300  0.65482  2.504  0.01228 *
## Stage.simpLate Stage  1.49883   4.47645  0.51553  2.907  0.00365 **
## ---
## Signif. codes:  0 '***' 0.001 '**' 0.01 '*' 0.05 '.' 0.1 ' ' 1
##
##                      exp(coef) exp(-coef) lower .95 upper .95
## MAPKAPK2                0.4059     2.4636    0.2213    0.7444
## Age                     1.0335     0.9676    0.9808    1.0889
## SexMale                 1.0916     0.9161    0.3279    3.6338
## SmokingYes              5.1530     0.1941    1.4278   18.5970
## Stage.simpLate Stage    4.4764     0.2234    1.6297   12.2958
##
## Concordance= 0.827  (se = 0.046 )
## Likelihood ratio test= 24.36  on 5 df,   p=2e-04
## Wald test            = 19.58  on 5 df,   p=0.001
## Score (logrank) test = 23.71  on 5 df,   p=2e-04
```

##### Cox PH with R censoring at 2 years, MK2 as a "hi" vs. "low" (dichotomous) variable

```
MK2_valid_t$MK2.topQ <- lapply(MK2_valid_t$MAPKAPK2, function(x) if (x >
    quantile(MK2_valid_t$MAPKAPK2, 0.66)) return(1) else return(0))
MK2_valid_t$MK2.topQ <- as.numeric(unlist(MK2_valid_t$MK2.topQ))
hist(MK2_valid_t$MK2.topQ)
```

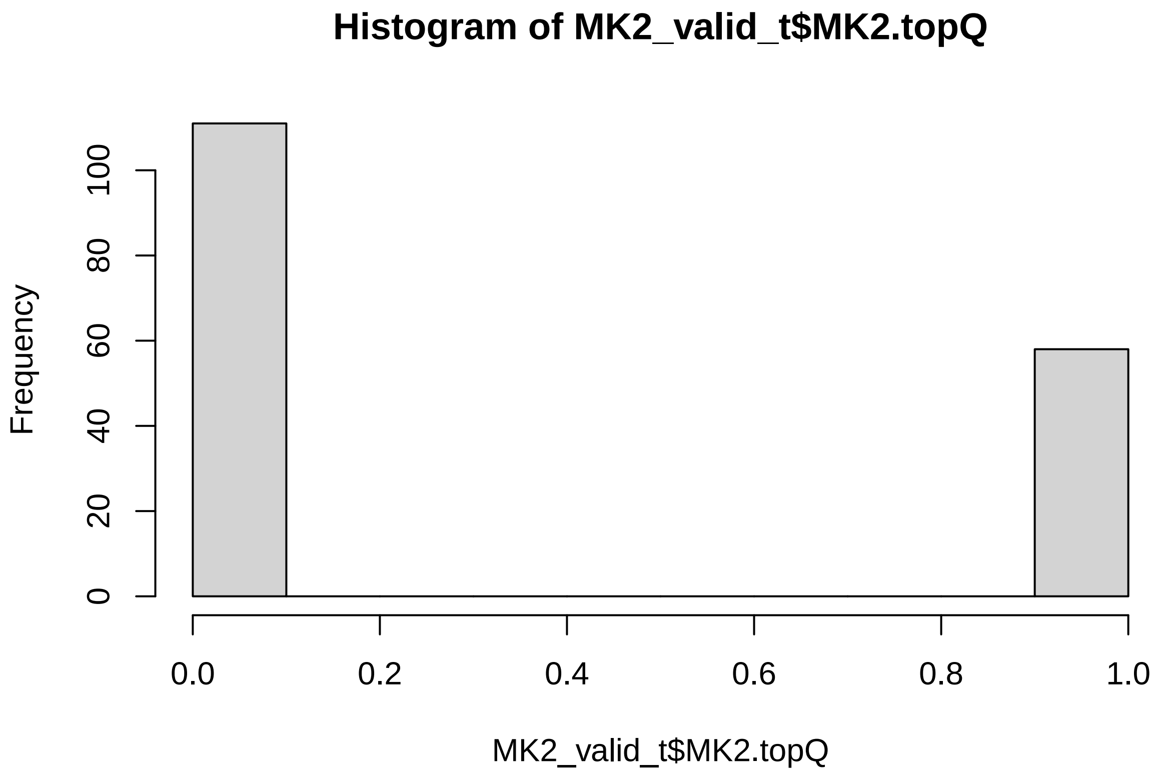

```
MK2_CoxPH_valid_RC_topQ <- coxph(Surv(time, censor_2yr) ~ MK2.topQ + Age +
    Sex + Smoking + Stage.simp, id = Sample.ID, data = MK2_valid_t)
cox.zph(MK2_CoxPH_valid_RC_topQ)
```

```
##            chisq df    p
## MK2.topQ   0.849  1 0.36
## Age        0.493  1 0.48
## Sex        1.393  1 0.24
## Smoking    0.769  1 0.38
## Stage.simp 0.596  1 0.44
## GLOBAL     3.335  5 0.65
```

```
summary(MK2_CoxPH_valid_RC_topQ)
```

```
## Call:
## coxph(formula = Surv(time, censor_2yr) ~ MK2.topQ + Age + Sex +
##     Smoking + Stage.simp, data = MK2_valid_t, id = Sample.ID)
##
##   n= 169, number of events= 16
##
##                            coef exp(coef) se(coef)      z Pr(>|z|)
## MK2.topQ               -2.24096   0.10636  1.03755 -2.160  0.03078 *
```

```
## Age                      0.02850   1.02891  0.02482  1.148  0.25087
## SexMale                  0.26098   1.29820  0.59959  0.435  0.66337
## SmokingYes               1.23779   3.44798  0.62527  1.980  0.04775 *
## Stage.simpLate Stage     1.33972   3.81799  0.50235  2.667  0.00766 **
## ---
## Signif. codes:  0 '***' 0.001 '**' 0.01 '*' 0.05 '.' 0.1 ' ' 1
##
##                        exp(coef) exp(-coef) lower .95 upper .95
## MK2.topQ                  0.1064     9.4024   0.01392    0.8127
## Age                       1.0289     0.9719   0.98005    1.0802
## SexMale                   1.2982     0.7703   0.40084    4.2045
## SmokingYes                3.4480     0.2900   1.01235   11.7435
## Stage.simpLate Stage      3.8180     0.2619   1.42637   10.2197
##
## Concordance= 0.836  (se = 0.043 )
## Likelihood ratio test= 23.95  on 5 df,   p=2e-04
## Wald test            = 19.84  on 5 df,   p=0.001
## Score (logrank) test = 24.04  on 5 df,   p=2e-04
```

##### Model graphics for the Cox PH model above - KM curves

```
km_valid <- survfit(Surv(time, censor_2yr) ~ MK2.topQ, data = MK2_valid_t,
    type = "kaplan-meier")
ggsurvplot(km_valid, conf.int = TRUE, xlim = c(0, 24), break.x.by = 2,
    pval = TRUE, font.y = 14, font.x = 14, font.tickslab = 14, ggtheme = theme_bw())
```

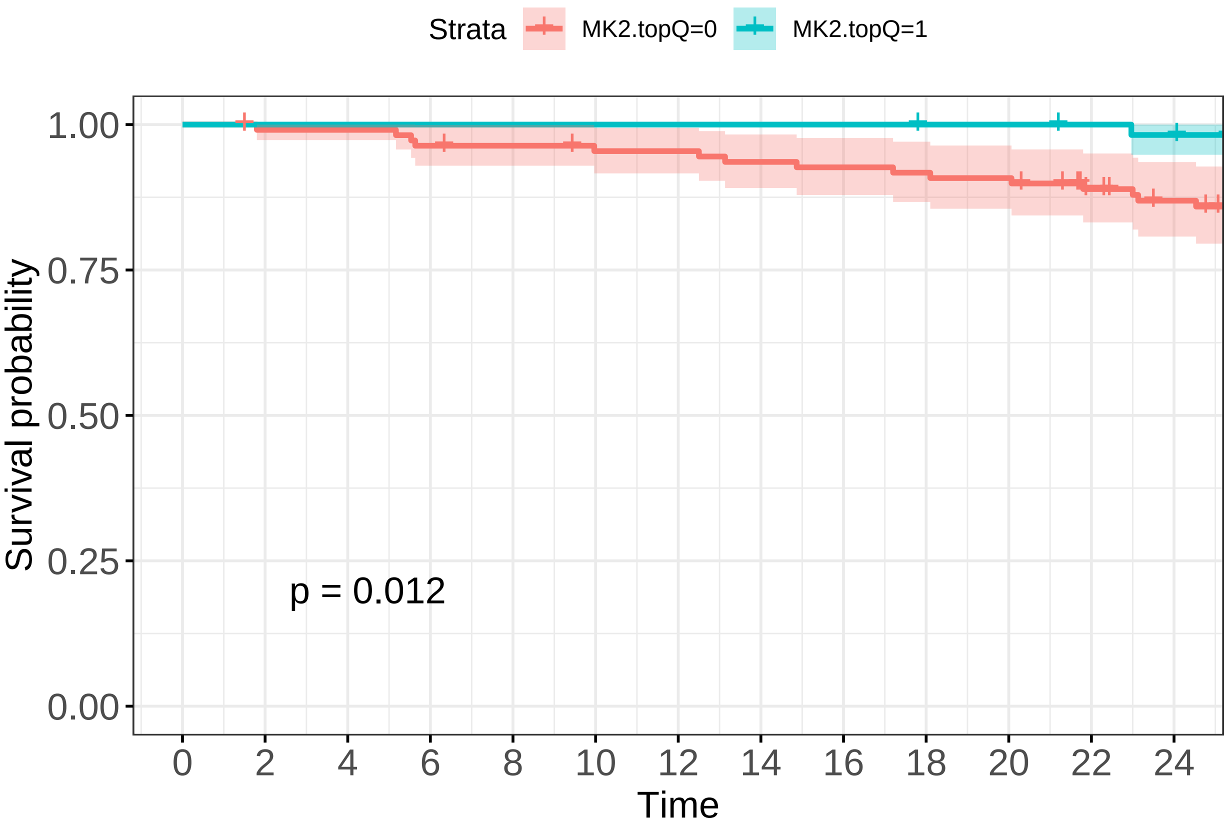

```
ggsurvplot(km_valid, conf.int = TRUE, risk.table = TRUE, pval = TRUE, font.y = 14,
    font.x = 14, font.tickslab = 14, ggtheme = theme_bw())
```

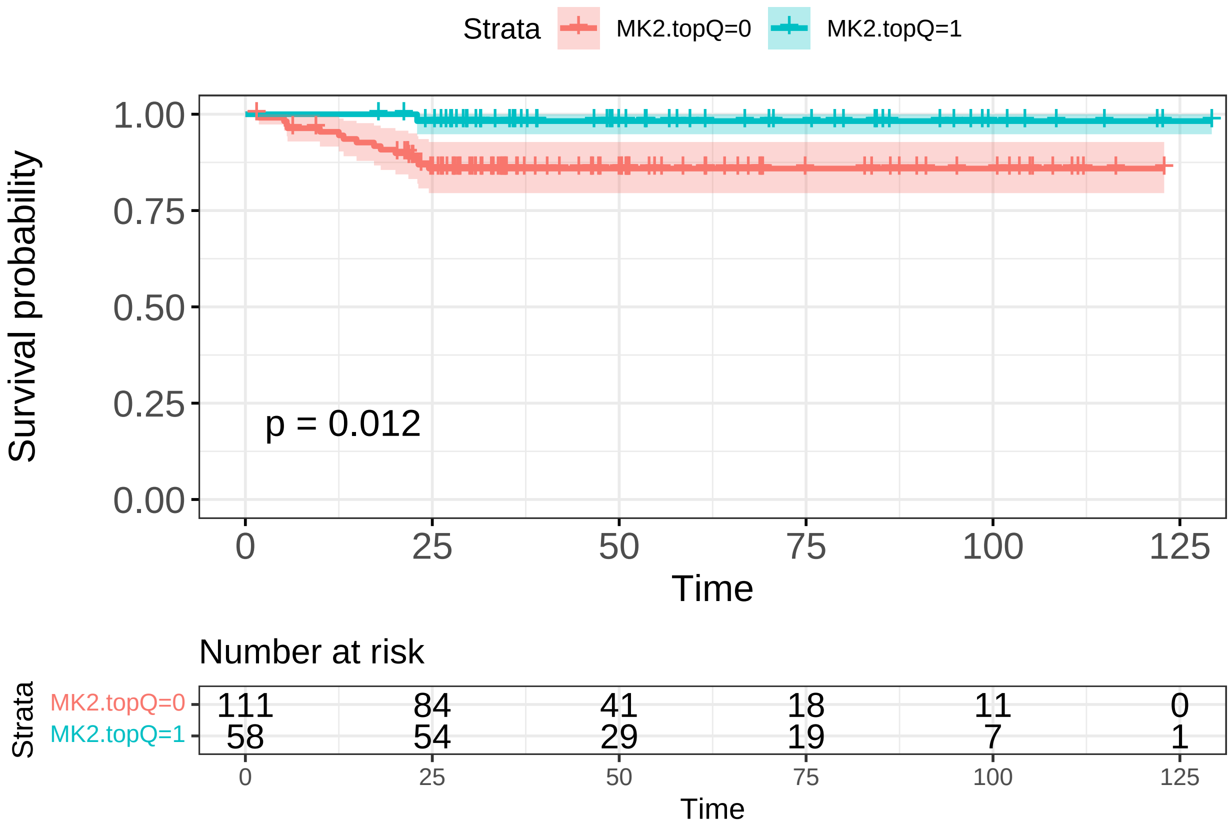

##### Logistic regression for death at 2 year

```
glm_valid_RC_topQ <- glm(censor_2yr ~ MK2.topQ + Age + Sex + Smoking +
    Stage.simp, data = MK2_valid_t, family = binomial(link = "logit"))
ResourceSelection::hoslem.test(glm_valid_RC_topQ$y, fitted(glm_valid_RC_topQ))
```

```
##
##  Hosmer and Lemeshow goodness of fit (GOF) test
##
## data:  glm_valid_RC_topQ$y, fitted(glm_valid_RC_topQ)
## X-squared = 5.1862, df = 8, p-value = 0.7375
```

```
summary(glm_valid_RC_topQ)
```

```
##
## Call:
## glm(formula = censor_2yr ~ MK2.topQ + Age + Sex + Smoking + Stage.simp,
##     family = binomial(link = "logit"), data = MK2_valid_t)
##
## Deviance Residuals:
##      Min        1Q    Median        3Q       Max
## -1.39541  -0.44702  -0.25153  -0.09478   2.67335
##
## Coefficients:
##                        Estimate Std. Error z value Pr(>|z|)
## (Intercept)            -5.58292    2.03162  -2.748  0.00600 **
```

```
## MK2.topQ             -2.43052    1.08706  -2.236  0.02536 *
## Age                   0.03575    0.02906   1.230  0.21852
## SexMale               0.27854    0.69235   0.402  0.68746
## SmokingYes            1.47837    0.70376   2.101  0.03567 *
## Stage.simpLate Stage  1.60790    0.60011   2.679  0.00738 **
## ---
## Signif. codes:  0 '***' 0.001 '**' 0.01 '*' 0.05 '.' 0.1 ' ' 1
##
## (Dispersion parameter for binomial family taken to be 1)
##
##     Null deviance: 105.869  on 168  degrees of freedom
## Residual deviance:  81.567  on 163  degrees of freedom
## AIC: 93.567
##
## Number of Fisher Scoring iterations: 7
```

```
exp(coefficients(glm_valid_RC_topQ))
```

```
##          (Intercept)            MK2.topQ                  Age
##          0.003761567         0.087990908          1.036401141
##              SexMale          SmokingYes Stage.simpLate Stage
##          1.321193700         4.385791796          4.992317722
```

```
exp(confint(glm_valid_RC_topQ))
```

```
## Waiting for profiling to be done...
```

```
##                             2.5 %     97.5 %
## (Intercept)          4.839724e-05  0.1541378
## MK2.topQ             4.595678e-03  0.4995502
## Age                  9.810628e-01  1.1011159
## SexMale              3.394220e-01  5.2842893
## SmokingYes           1.153863e+00 18.8912207
## Stage.simpLate Stage 1.545490e+00 16.8396107
```

```
listVars <- c("Age", "MAPKAPK2", "Chemotherapy", "Stage.simp", "Sex", "Smoking")
catVars <- c("Chemotherapy", "Stage.simp", "Sex", "Smoking")
table1_validationcohort <- tableone::CreateTableOne(listVars, MK2_valid_t,
    catVars, strata = c("MK2.topQ"), addOverall = TRUE)
print(table1_validationcohort)
```

```
##                               Stratified by MK2.topQ
##                                Overall       0             1             p
##    n                             169           111            58
##    Age (mean (SD))             64.00 (9.47)  63.96 (10.23) 64.07 (7.91)   0.946
##    MAPKAPK2 (mean (SD))         0.00 (1.00)  -0.56 (0.60)   1.07 (0.69)  <0.001
##    Chemotherapy = Yes (%)         53 (31.4)     29 (26.1)     24 (41.4)   0.064
##    Stage.simp = Late Stage (%)    37 (21.9)     25 (22.5)     12 (20.7)   0.938
##    Sex = Male (%)                 75 (44.4)     44 (39.6)     31 (53.4)   0.121
##    Smoking = Yes (%)              61 (36.1)     41 (36.9)     20 (34.5)   0.883
##                               Stratified by MK2.topQ
##                                test
##    n
##    Age (mean (SD))
##    MAPKAPK2 (mean (SD))
##    Chemotherapy = Yes (%)
##    Stage.simp = Late Stage (%)
```

```
##   Sex = Male (%)
##   Smoking = Yes (%)
```

```
## R version 4.1.0 (2021-05-18)
## Platform: x86_64-pc-linux-gnu (64-bit)
## Running under: Ubuntu 20.04.2 LTS
##
## Matrix products: default
## BLAS:   /usr/lib/x86_64-linux-gnu/blas/libblas.so.3.9.0
## LAPACK: /usr/lib/x86_64-linux-gnu/lapack/liblapack.so.3.9.0
##
## locale:
##  [1] LC_CTYPE=en_US.UTF-8       LC_NUMERIC=C
##  [3] LC_TIME=en_US.UTF-8        LC_COLLATE=en_US.UTF-8
##  [5] LC_MONETARY=en_US.UTF-8    LC_MESSAGES=en_US.UTF-8
##  [7] LC_PAPER=en_US.UTF-8       LC_NAME=C
##  [9] LC_ADDRESS=C               LC_TELEPHONE=C
## [11] LC_MEASUREMENT=en_US.UTF-8 LC_IDENTIFICATION=C
##
## attached base packages:
## [1] stats     graphics  grDevices utils     datasets  methods   base
##
## other attached packages:
## [1] survminer_0.4.9   survival_3.2-11   dplyr_1.0.7       ggpubr_0.4.0
## [5] ggplot2_3.3.5     data.table_1.14.0 knitr_1.33
##
## loaded via a namespace (and not attached):
##  [1] tidyr_1.1.3             splines_4.1.0           carData_3.0-4
##  [4] assertthat_0.2.1        highr_0.9               cellranger_1.1.0
##  [7] yaml_2.2.1              pillar_1.6.2            backports_1.2.1
## [10] lattice_0.20-44         glue_1.4.2              digest_0.6.27
## [13] ggsignif_0.6.2          gridtext_0.1.4          ResourceSelection_0.3-5
## [16] colorspace_2.0-2        htmltools_0.5.2         Matrix_1.3-4
## [19] survey_4.0              pkgconfig_2.0.3         labelled_2.8.0
## [22] broom_0.7.8             haven_2.4.1             purrr_0.3.4
## [25] xtable_1.8-4            scales_1.1.1            km.ci_0.5-2
## [28] openxlsx_4.2.4          rio_0.5.27              KMsurv_0.1-5
## [31] proxy_0.4-26            tibble_3.1.4            mgcv_1.8-36
## [34] generics_0.1.0          farver_2.1.0            car_3.0-11
## [37] ellipsis_0.3.2          withr_2.4.2             magrittr_2.0.1
## [40] crayon_1.4.1            readxl_1.3.1            ggtext_0.1.1
## [43] evaluate_0.14           fansi_0.5.0             nlme_3.1-152
## [46] MASS_7.3-54             class_7.3-19            rstatix_0.7.0
## [49] forcats_0.5.1           xml2_1.3.2              foreign_0.8-81
## [52] tableone_0.13.0         tools_4.1.0             hms_1.1.0
## [55] mitools_2.4             formatR_1.11            lifecycle_1.0.0
## [58] stringr_1.4.0           munsell_0.5.0           zip_2.2.0
## [61] e1071_1.7-7             compiler_4.1.0          rlang_0.4.11
## [64] grid_4.1.0              labeling_0.4.2          rmarkdown_2.11
## [67] gtable_0.3.0            abind_1.4-5             DBI_1.1.1
## [70] curl_4.3.2              markdown_1.1            R6_2.5.1
## [73] gridExtra_2.3           zoo_1.8-9               fastmap_1.1.0
## [76] survMisc_0.5.5          utf8_1.2.2              stringi_1.7.4
## [79] Rcpp_1.0.7              vctrs_0.3.8             tidyselect_1.1.1
```

```
## [82] xfun_0.25
```

Note that the `echo = FALSE` parameter was added to the code chunk to prevent printing of the R code that generated the plot.

### Pan Cancer MK2 Survival Analysis

Karthik Suresh

3/11/2020

Updates in Revision 2:

1. We updated the Cox PH model to the model that was used for the LUAD dataset.
2. We updated the figures for the HR across cancer datasets to also include metrics of the Cox PH models used in each dataset.

The MasterMK2data.csv file contains MK2 transcript levels + clinical data across a variety of cancer datasets. Please see the MK2__panCA__annotation pdf for the code that we used to pull the clinical data from the TCGA. The MK2 transcript data was pulled from OncoLnc (manually).

```
library(knitr)
opts_chunk$set(tidy.opts=list(width.cutoff=70), tidy=TRUE)
```

#### Create MK2 categories (low vs. high)

An added issue here is that we need to define quantiles WITHIN tumor groups. So the quantile for each tumor group is different.

```
MK2_data <- MK2_data[!is.na(MK2_data$Expression), ]
# create MK2 quantiles
bottom_quant <<- 0.33
top_quant <<- 0.66

BelowBottomQuantile <- function(cancer.type, x) {
    if (x < quantile(MK2_data[MK2_data$cancer.type.x == cancer.type, ]$Expression,
        bottom_quant))
        return(1) else return(0)
}
AboveTopQuantile <- function(cancer.type, x) {
    if (x > quantile(MK2_data[MK2_data$cancer.type.x == cancer.type, ]$Expression,
        top_quant))
        return(1) else return(0)
}
MK2_data$MK2_Expression_bottomQ <- mapply(BelowBottomQuantile, MK2_data$cancer.type.x,
    MK2_data$Expression)
MK2_data$MK2_Expression_topQ <- mapply(AboveTopQuantile, MK2_data$cancer.type.x,
    MK2_data$Expression)

MK2_data_tb <- MK2_data
MK2_data_tb$MK2_Expression_topQ <- as.numeric(MK2_data_tb$MK2_Expression_topQ)
MK2_data_tb$MK2_Expression_bottomQ <- as.numeric(MK2_data_tb$MK2_Expression_bottomQ)
```

```r
# re-categorize the variable as MK2_tv where 0 = lower 10th, 1= top
# 10th
MK2_data_tb$MK2_tv <- mapply(function(bottomQ, topQ) if (topQ == 1) return("High") else return("Low"),
    MK2_data_tb$MK2_Expression_bottomQ, MK2_data_tb$MK2_Expression_topQ)
MK2_data_tb$MK2_tv <- as.factor(MK2_data_tb$MK2_tv)

MK2_data_tb <- MK2_data_tb[!is.na(MK2_data_tb$time), ]
# create a 2 year censor
MK2_data_tb$death_at_2year <- mapply(function(dead, time) if (dead == 1 &
    time < 366 * 2) return(1) else if (dead == 0 & time > 366 * 2) return(0) else if (dead ==
    0 & time < 366 * 2) return(0) else if (dead == 1 & time > 366 * 2) return(0),
    MK2_data_tb$censor, as.numeric(MK2_data_tb$time))
MK2_data_tb$death_at_2year <- as.numeric(unlist(as.character(MK2_data_tb$death_at_2year)))
```

```
## Warning: NAs introduced by coercion
```

```r
MK2_data_tb <- MK2_data_tb[MK2_data_tb$death_at_2year == 0 | MK2_data_tb$death_at_2year ==
    1, ]
```

#### Run Cox PH on the entire dataset

So, here we look at the effect of MK2 on the entire dataset, adjusting for cancer type and stage.

```r
hist(MK2_data_tb$Expression)
```

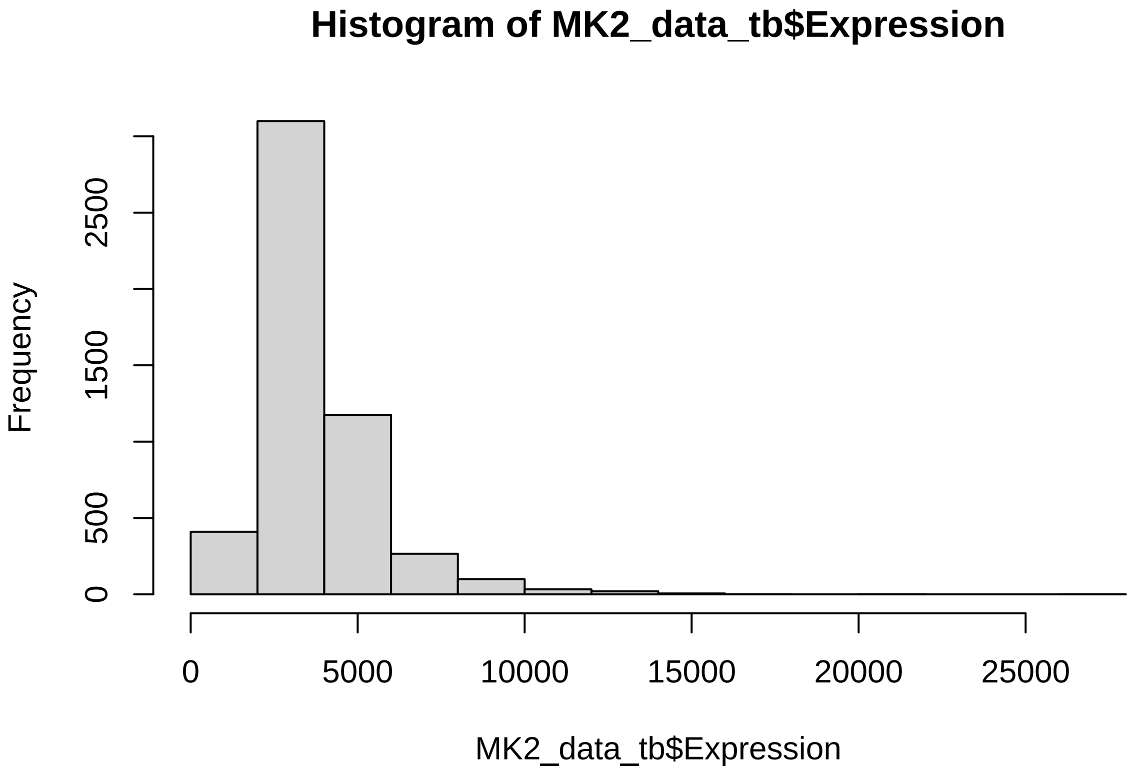

```r
MK2_data_tb$Expression_logt <- log(MK2_data_tb$Expression)

coxph_full <- coxph(Surv(time, death_at_2year) ~ Expression_logt + ajcc_pathologic_stage_combined +
    age_at_diagnosis + gender + smoking + as.factor(cancer.type.x), data = MK2_data_tb)
```

```r
cox.zph(coxph_full)
```

```
##                                    chisq df       p
## Expression_logt                    0.453  1    0.50
## ajcc_pathologic_stage_combined     0.557  1    0.46
## age_at_diagnosis                  17.907  1 2.3e-05
## gender                             1.237  1    0.27
## smoking                            1.343  1    0.25
## as.factor(cancer.type.x)          46.437 12 5.8e-06
## GLOBAL                            67.180 17 6.6e-08
```

```r
summary(coxph_full)
```

```
## Call:
## coxph(formula = Surv(time, death_at_2year) ~ Expression_logt +
##     ajcc_pathologic_stage_combined + age_at_diagnosis + gender +
##     smoking + as.factor(cancer.type.x), data = MK2_data_tb)
##
##   n= 5078, number of events= 989
##    (35 observations deleted due to missingness)
##
##                                              coef exp(coef) se(coef)      z
## Expression_logt                           0.17235   1.18810  0.09891  1.742
## ajcc_pathologic_stage_combinedLate Stage  1.03678   2.82011  0.07077 14.650
## age_at_diagnosis                          0.02365   1.02393  0.00299  7.910
## gendermale                                0.05464   1.05616  0.06985  0.782
## smoking1                                 -0.08319   0.92018  0.09720 -0.856
## as.factor(cancer.type.x)BRCA             -1.82178   0.16174  0.20620 -8.835
## as.factor(cancer.type.x)COAD             -0.75287   0.47101  0.16045 -4.692
## as.factor(cancer.type.x)ESCA              0.29541   1.34367  0.18692  1.580
## as.factor(cancer.type.x)KIRC             -0.72921   0.48229  0.15386 -4.739
## as.factor(cancer.type.x)KIRP             -1.06740   0.34390  0.23383 -4.565
## as.factor(cancer.type.x)LIHC              0.15580   1.16859  0.15296  1.019
## as.factor(cancer.type.x)LUAD             -0.13029   0.87784  0.14033 -0.928
## as.factor(cancer.type.x)LUSC              0.12708   1.13551  0.13467  0.944
## as.factor(cancer.type.x)PAAD              1.13055   3.09736  0.15347  7.367
## as.factor(cancer.type.x)READ             -1.31750   0.26780  0.29513 -4.464
## as.factor(cancer.type.x)SKCM             -0.64071   0.52692  0.15691 -4.083
## as.factor(cancer.type.x)STAD              0.20768   1.23082  0.13384  1.552
##                                          Pr(>|z|)
## Expression_logt                            0.0814 .
## ajcc_pathologic_stage_combinedLate Stage  < 2e-16 ***
## age_at_diagnosis                         2.57e-15 ***
## gendermale                                 0.4340
## smoking1                                   0.3921
## as.factor(cancer.type.x)BRCA              < 2e-16 ***
## as.factor(cancer.type.x)COAD             2.70e-06 ***
## as.factor(cancer.type.x)ESCA               0.1140
## as.factor(cancer.type.x)KIRC             2.14e-06 ***
## as.factor(cancer.type.x)KIRP             5.00e-06 ***
## as.factor(cancer.type.x)LIHC               0.3084
## as.factor(cancer.type.x)LUAD               0.3532
## as.factor(cancer.type.x)LUSC               0.3454
## as.factor(cancer.type.x)PAAD             1.75e-13 ***
```

```
## as.factor(cancer.type.x)READ            8.04e-06 ***
## as.factor(cancer.type.x)SKCM            4.44e-05 ***
## as.factor(cancer.type.x)STAD              0.1207
## ---
## Signif. codes:  0 '***' 0.001 '**' 0.01 '*' 0.05 '.' 0.1 ' ' 1
##
##                                         exp(coef) exp(-coef) lower .95
## Expression_logt                            1.1881     0.8417    0.9787
## ajcc_pathologic_stage_combinedLate Stage    2.8201     0.3546    2.4549
## age_at_diagnosis                           1.0239     0.9766    1.0179
## gendermale                                 1.0562     0.9468    0.9210
## smoking1                                   0.9202     1.0867    0.7606
## as.factor(cancer.type.x)BRCA               0.1617     6.1829    0.1080
## as.factor(cancer.type.x)COAD               0.4710     2.1231    0.3439
## as.factor(cancer.type.x)ESCA               1.3437     0.7442    0.9315
## as.factor(cancer.type.x)KIRC               0.4823     2.0734    0.3567
## as.factor(cancer.type.x)KIRP               0.3439     2.9078    0.2175
## as.factor(cancer.type.x)LIHC               1.1686     0.8557    0.8659
## as.factor(cancer.type.x)LUAD               0.8778     1.1392    0.6668
## as.factor(cancer.type.x)LUSC               1.1355     0.8807    0.8721
## as.factor(cancer.type.x)PAAD               3.0974     0.3229    2.2928
## as.factor(cancer.type.x)READ               0.2678     3.7341    0.1502
## as.factor(cancer.type.x)SKCM               0.5269     1.8978    0.3874
## as.factor(cancer.type.x)STAD               1.2308     0.8125    0.9468
##                                         upper .95
## Expression_logt                            1.4423
## ajcc_pathologic_stage_combinedLate Stage    3.2397
## age_at_diagnosis                           1.0299
## gendermale                                 1.2111
## smoking1                                   1.1133
## as.factor(cancer.type.x)BRCA               0.2423
## as.factor(cancer.type.x)COAD               0.6451
## as.factor(cancer.type.x)ESCA               1.9382
## as.factor(cancer.type.x)KIRC               0.6520
## as.factor(cancer.type.x)KIRP               0.5438
## as.factor(cancer.type.x)LIHC               1.5771
## as.factor(cancer.type.x)LUAD               1.1558
## as.factor(cancer.type.x)LUSC               1.4785
## as.factor(cancer.type.x)PAAD               4.1843
## as.factor(cancer.type.x)READ               0.4776
## as.factor(cancer.type.x)SKCM               0.7166
## as.factor(cancer.type.x)STAD               1.6000
##
## Concordance= 0.741  (se = 0.007 )
## Likelihood ratio test= 751.5  on 17 df,   p=<2e-16
## Wald test            = 629.1  on 17 df,   p=<2e-16
## Score (logrank) test = 753.2  on 17 df,   p=<2e-16
```

### Collect Cox PH info

Now, we run our pre-specified Cox PH model iteratively, for each dataset, and the functions below return either the model result or model metrics.

```r
# Some functions to pull CoxPH models and metrics for all cancers.

returnModel <- function(cancer.type) {
    surv_df <- MK2_data_tb[MK2_data_tb$cancer.type.x == cancer.type, ]
    MK2_model <- coxph(Surv(time, death_at_2year) ~ MK2_Expression_topQ +
        ajcc_pathologic_stage_combined + age_at_diagnosis + gender + smoking,
        data = surv_df)

    return(MK2_model)

}

returnModel_metrics <- function(cancer.type) {
    surv_df <- MK2_data_tb[MK2_data_tb$cancer.type.x == cancer.type, ]
    MK2_model <- coxph(Surv(time, death_at_2year) ~ MK2_Expression_topQ +
        ajcc_pathologic_stage_combined + age_at_diagnosis + gender + smoking,
        data = surv_df)

    return(list(anova(MK2_model), cox.zph(MK2_model)))

}

returnModel_MK2cont <- function(cancer.type) {
    surv_df <- MK2_data_tb[MK2_data_tb$cancer.type.x == cancer.type, ]
    MK2_model <- coxph(Surv(time, death_at_2year) ~ Expression_logt + ajcc_pathologic_stage_combined +
        age_at_diagnosis + gender + smoking, data = surv_df)

    return(MK2_model)

}
returnModel_MK2cont_metrics <- function(cancer.type) {
    surv_df <- MK2_data_tb[MK2_data_tb$cancer.type.x == cancer.type, ]
    MK2_model <- coxph(Surv(time, death_at_2year) ~ Expression_logt + ajcc_pathologic_stage_combined +
        age_at_diagnosis + gender + smoking, data = surv_df)

    return(list(anova(MK2_model), cox.zph(MK2_model)))

}
```

Now that we have some prepared functions, we use them to call the model and retrieve model metrics.

Cox PH Model 1: MK2 as a dichotomous variable + covariates (including stage) Cox PH Model 2: MK2 as a continuous variable + covariates (including stage)

Cancer type is not included because, in essence, we're stratifying by cancer type here.

```r
# return model fit parameters (cox.zph)
MK2_model_df <- MK2_means_wide

# model MK2 as a dichotomous var
MK2_model_df$model1.results <- lapply(MK2_model_df$cancer.type.x, returnModel)

MK2_model_df$model1.metrics <- lapply(MK2_model_df$cancer.type.x, returnModel_metrics)
```

```
MK2_model_df$cox.zph_MK2 <- sapply(MK2_model_df$model1.metrics, function(x) return(x[[2]][[1]][13]))
MK2_model_df$cox.zph_global <- sapply(MK2_model_df$model1.metrics, function(x) return(x[[2]][[1]][18]))
MK2_model_df$wald_pval <- sapply(MK2_model_df$model1.results, function(x) return(broom::glance(x)$p.valu

MK2_model_df$MK2 <- lapply(MK2_model_df$model1.results, function(x) return((x$coefficients[[1]][1])))
MK2_model_df$MK2.lci <- lapply(MK2_model_df$model1.results, function(x) return(exp(confint(x)[1])))
MK2_model_df$MK2.uci <- lapply(MK2_model_df$model1.results, function(x) return(exp(confint(x)[6])))

MK2_model_df$MK2 <- exp(as.numeric(MK2_model_df$MK2))

# model MK2 as a continuous var
MK2_model_df$model2.results <- lapply(MK2_model_df$cancer.type.x, returnModel_MK2cont)
MK2_model_df$MK2_model2 <- lapply(MK2_model_df$model2.results, function(x) return(x$coefficients[[1]][1
MK2_model_df$MK2_model2.lci <- lapply(MK2_model_df$model2.results, function(x) return(exp(confint(x)[1]
MK2_model_df$MK2_model2.uci <- lapply(MK2_model_df$model2.results, function(x) return(exp(confint(x)[6]
MK2_model_df$MK2_model2 <- exp(as.numeric(MK2_model_df$MK2_model2))

MK2_model_df$model2.metrics <- lapply(MK2_model_df$cancer.type.x, returnModel_MK2cont_metrics)
MK2_model_df$cox.zph_MK2_model2 <- sapply(MK2_model_df$model2.metrics,
    function(x) return(x[[2]][[1]][13]))
MK2_model_df$cox.zph_global_model2 <- sapply(MK2_model_df$model2.metrics,
    function(x) return(x[[2]][[1]][18]))
MK2_model_df$wald_pval_model2 <- sapply(MK2_model_df$model2.results, function(x) return(broom::glance(x)
```

#### Graphical output: graphs of MK2 transcript levels and HR for MK2 across datasets

```
ggplot(MK2_model_df, aes(x = reorder(cancer.type.x, -as.numeric(MK2)),
    y = as.numeric(MK2))) + geom_bar(stat = "identity") + geom_hline(yintercept = 1) +
    theme_bw() + coord_flip() + ylab("HR for high MK2 transcript level in Cox PH model adjusted for stag
    xlab("Cancer Type")
```

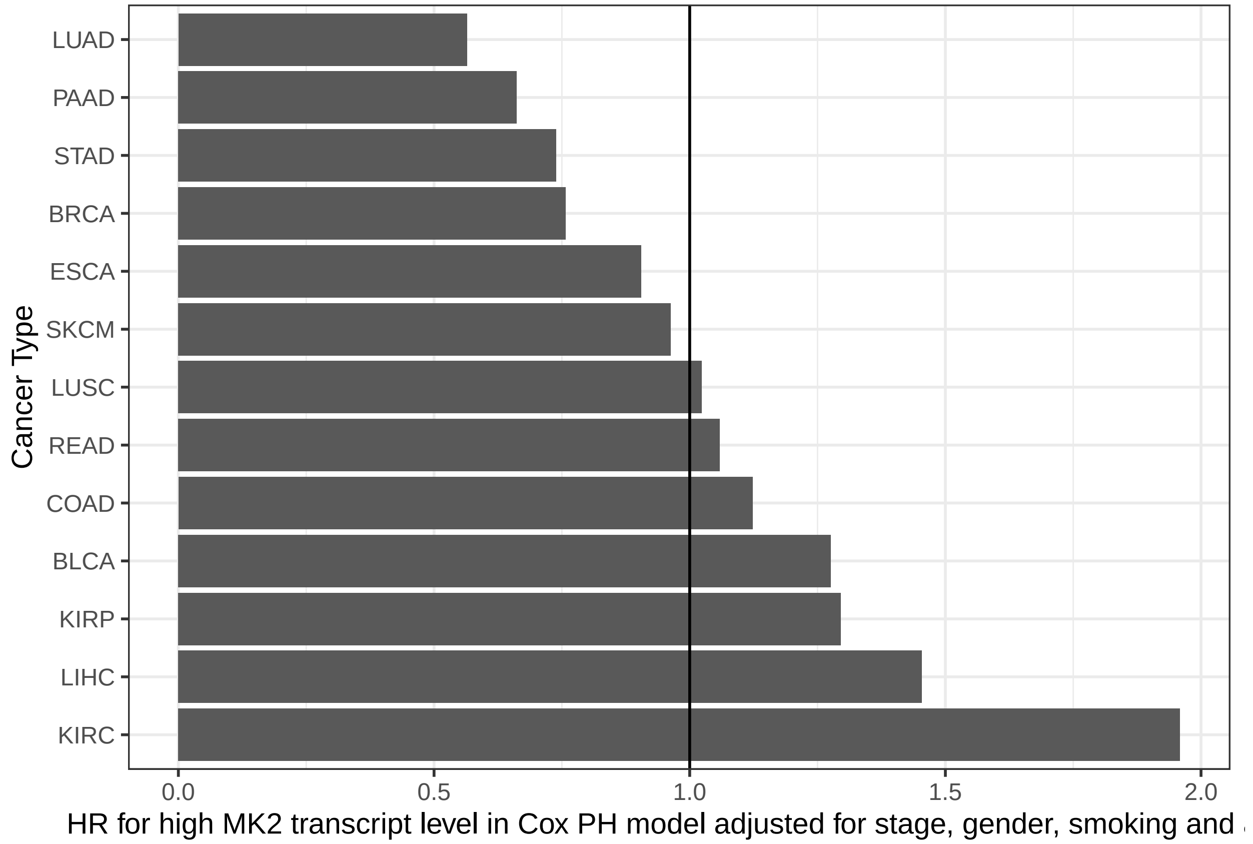

```
ggplot(MK2_model_df, aes(x = reorder(cancer.type.x, -diff), y = diff)) +
    geom_bar(stat = "identity") + theme_bw() + coord_flip() + ylab("Difference in mean MK2 transcript le
    xlab("TCGA Cancer dataset")
```

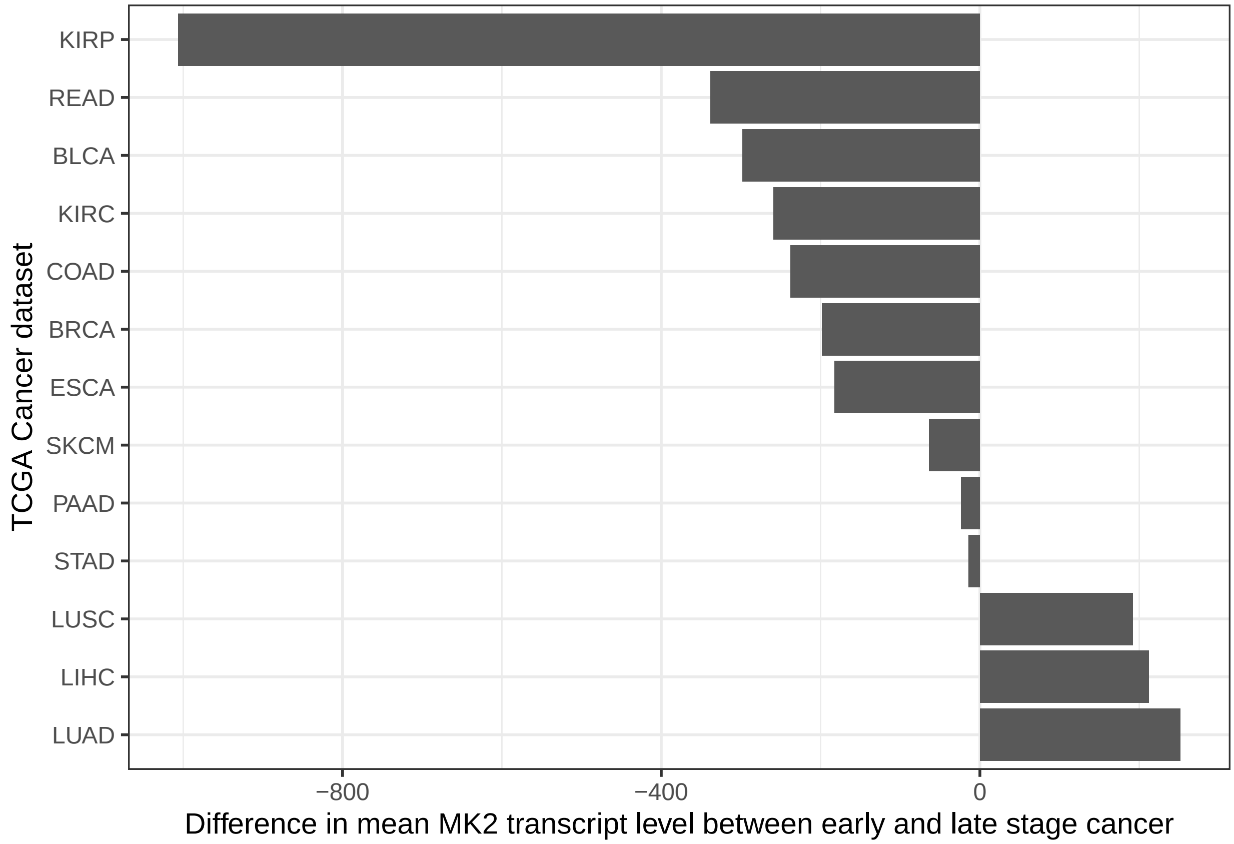

```
MK2_model_df$cancer.type.x.label <- sapply(MK2_model_df$cancer.type.x,
    function(x) return(pts_percatype[pts_percatype$cancer_type == x, ]$num))
MK2_model_df$cancer.type.x.label <- paste0(MK2_model_df$cancer.type.x,
    " (n=", MK2_model_df$cancer.type.x.label, ")")

# Model 1: MK2 as a dichotomous variable

HR_plot <- ggplot(MK2_model_df, aes(x = reorder(cancer.type.x.label, -wald_pval),
    y = as.numeric(MK2))) + geom_point() + geom_errorbar(aes(ymin = as.numeric(MK2.lci),
    ymax = as.numeric(MK2.uci)), width = 0) + geom_hline(yintercept = 1) +
    theme_bw() + ylab("HR for high MK2 transcript level in Cox PH model adjusted for stage, gender, smol
    xlab("Cancer Type") + coord_flip()

model_stats_df <- MK2_model_df[c("cancer.type.x", "cox.zph_MK2", "wald_pval")]
model_stats_df <- arrange(model_stats_df, wald_pval)
metrics_plot_wald <- ggplot(model_stats_df, aes(x = reorder(cancer.type.x,
    -wald_pval), y = log(wald_pval))) + geom_point(size = 4) + geom_hline(yintercept = log(0.05)) +
    geom_segment(aes(x = cancer.type.x, xend = cancer.type.x, y = log(0.05),
        yend = log(wald_pval))) + theme_bw() + xlab("") + ylab("log p value") +
    coord_flip()

metrics_plot_PH <- ggplot(model_stats_df, aes(x = reorder(cancer.type.x,
    -wald_pval), y = cox.zph_MK2)) + geom_point(size = 4) + geom_hline(yintercept = 0.05) +
    geom_segment(aes(x = cancer.type.x, xend = cancer.type.x, y = 0.05,
        yend = cox.zph_MK2)) + theme_bw() + xlab("") + ylab("p value") +
```

```
    coord_flip()

gridExtra::grid.arrange(HR_plot, metrics_plot_wald, metrics_plot_PH, ncol = 3,
    widths = c(4, 1, 1))
```

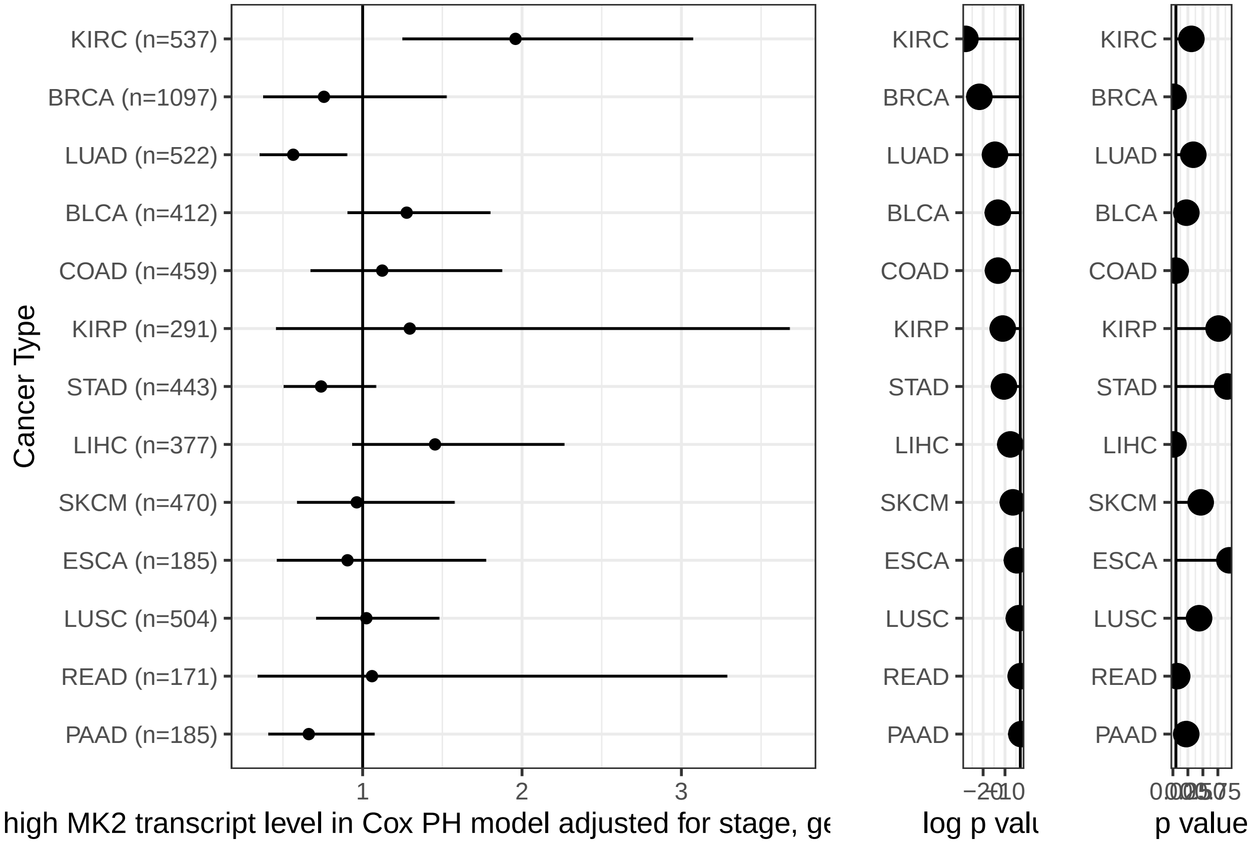

```
# model 2 results plots

model2_stats_df <- MK2_model_df[c("cancer.type.x", "cox.zph_MK2_model2",
    "wald_pval_model2")]

HR_plot_model2 <- ggplot(MK2_model_df, aes(x = reorder(cancer.type.x.label,
    -wald_pval_model2), y = as.numeric(MK2_model2))) + geom_point() + geom_errorbar(aes(ymin = as.numeri
    ymax = as.numeric(MK2_model2.uci)), width = 0) + geom_hline(yintercept = 1) +
    theme_bw() + ylab("HR for MK2 transcript level in Cox PH model adjusted for stage, gender, smoking a
    xlab("Cancer Type") + coord_flip()

metrics_plot_wald_model2 <- ggplot(model2_stats_df, aes(x = reorder(cancer.type.x,
    -wald_pval_model2), y = log(wald_pval_model2))) + geom_point(size = 4) +
    geom_hline(yintercept = log(0.05)) + geom_segment(aes(x = cancer.type.x,
    xend = cancer.type.x, y = log(0.05), yend = log(wald_pval_model2))) +
    theme_bw() + xlab("") + ylab("log p value") + coord_flip()

metrics_plot_PH_model2 <- ggplot(model2_stats_df, aes(x = reorder(cancer.type.x,
    -wald_pval_model2), y = cox.zph_MK2_model2)) + geom_point(size = 4) +
    geom_hline(yintercept = 0.05) + geom_segment(aes(x = cancer.type.x,
    xend = cancer.type.x, y = 0.05, yend = cox.zph_MK2_model2)) + theme_bw() +
```

```
    xlab("") + ylab("p value") + coord_flip()

gridExtra::grid.arrange(HR_plot_model2, metrics_plot_wald_model2, metrics_plot_PH_model2,
    ncol = 3, widths = c(4, 1, 1))
```

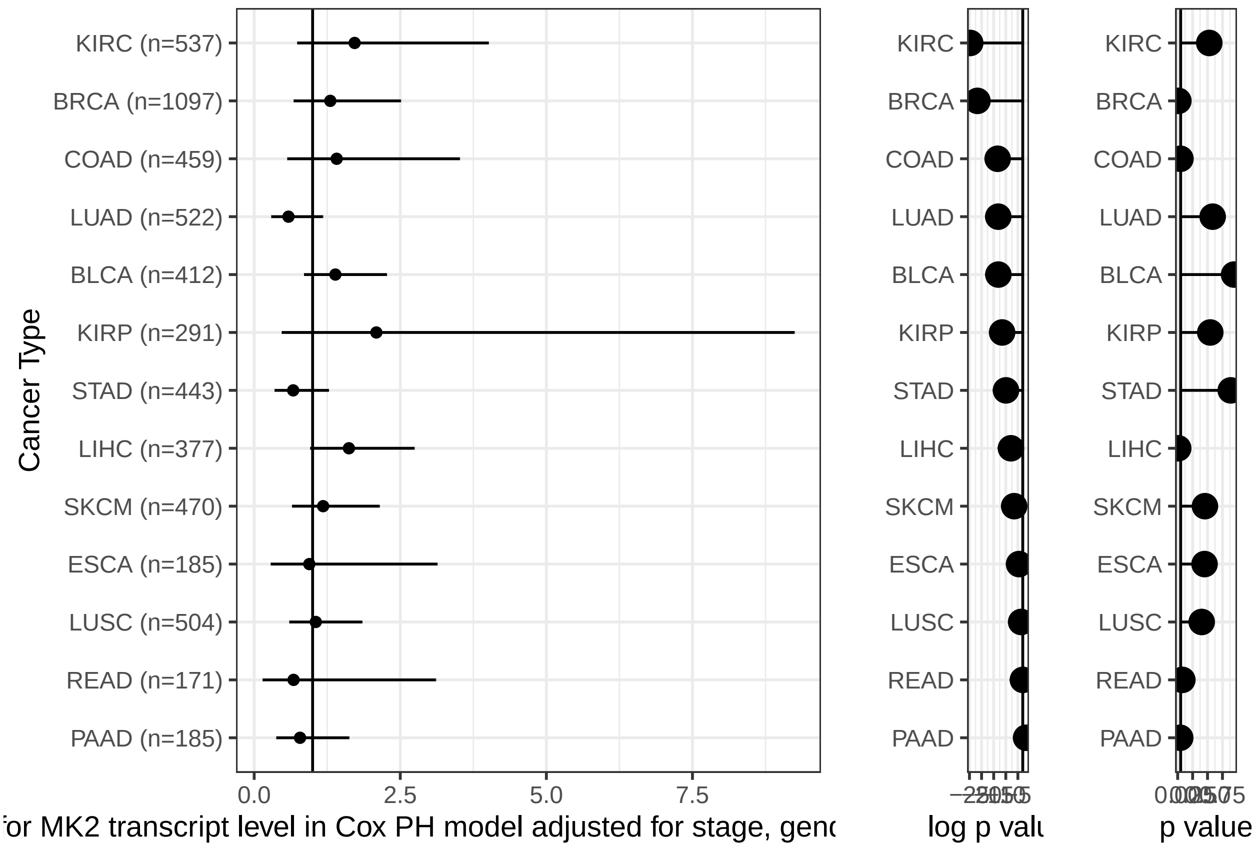

#### R Markdown

This is an R Markdown document. Markdown is a simple formatting syntax for authoring HTML, PDF, and MS Word documents. For more details on using R Markdown see http://rmarkdown.rstudio.com.

When you click the **Knit** button a document will be generated that includes both content as well as the output of any embedded R code chunks within the document. You can embed an R code chunk like this:

```
summary(cars)
```

```
##      speed           dist
##  Min.   : 4.0   Min.   :  2.00
##  1st Qu.:12.0   1st Qu.: 26.00
##  Median :15.0   Median : 36.00
##  Mean   :15.4   Mean   : 42.98
##  3rd Qu.:19.0   3rd Qu.: 56.00
##  Max.   :25.0   Max.   :120.00
```

#### Including Plots

You can also embed plots, for example:

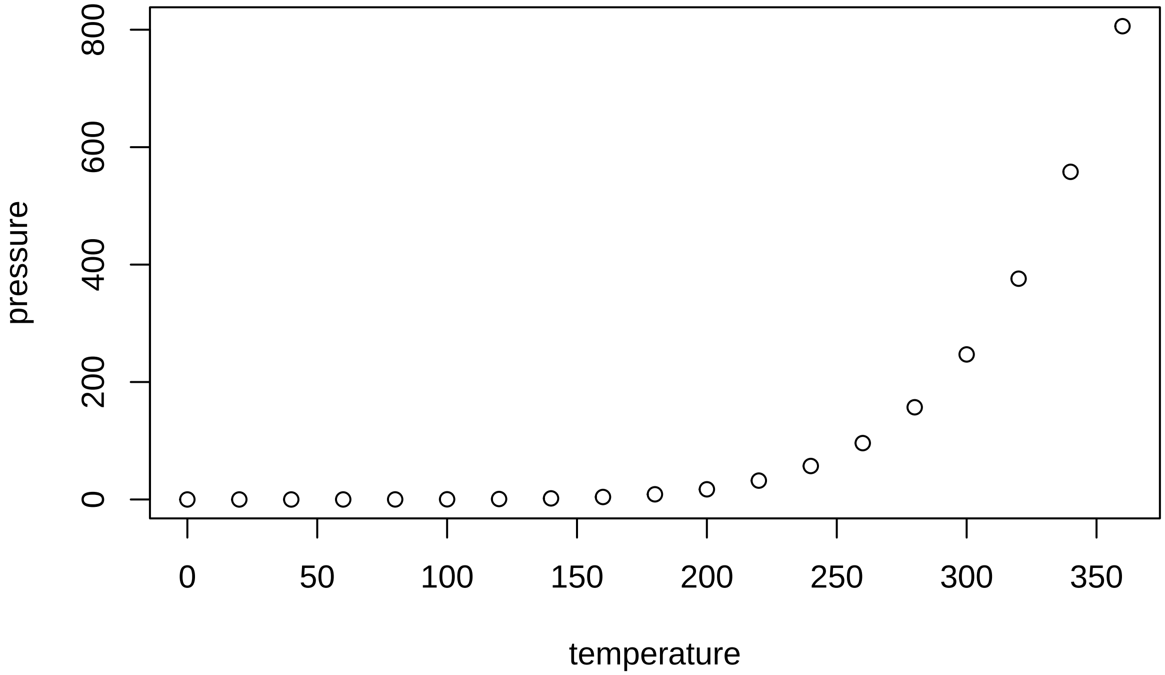

Note that the `echo = FALSE` parameter was added to the code chunk to prevent printing of the R code that generated the plot.

### Pan Cancer MK2 transcript clinical annotation

Karthik Suresh

3/11/2020

We have already downloaded transcript data for MK2 across a variety of cancers. Here, we need to collate all the transcript data, go find all the relevant clinical data using GDC_query, annotate, and save a giant file containing transcript and clinical data across many cancers.

```
library(knitr)
opts_chunk$set(eval=FALSE, echo=TRUE)
```

```
knitr::opts_chunk$set(echo = TRUE, eval=FALSE)
library("SummarizedExperiment")
library("dplyr")
library("DT")
library("TCGAbiolinks")
library("plyr")
library("dplyr")
rm(list=ls())

filelist <- as.data.frame(list.files("../rawdata/panca.mk2.oncolnc/"))
colnames(filelist)<- c("filename")
filelist$cancer.type <- lapply(as.character(filelist$filename), function(x) return(unlist(strsplit(x,"_

GetTranscriptData <- function(file, ca.type) {

if(!exists("MasterTranscriptList")) {
  MasterTranscriptList <<- read.csv(paste0("../rawdata/panca.mk2.oncolnc/", file))
  MasterTranscriptList$cancer.type <<- ca.type
}
else {
 currTranscriptList <<- read.csv(paste0("../rawdata/panca.mk2.oncolnc/", file))
  currTranscriptList$cancer.type <<- ca.type
  MasterTranscriptList <<- rbind(MasterTranscriptList, currTranscriptList)
  }
return(1)
}

filelist$collectTranscript <- mapply(GetTranscriptData, filelist$filename, filelist$cancer.type)

GetClinicalData <- function(ca.type) {
 if(!exists("MasterClinicalList")) {

  MasterClinicalList <<-  GDCquery_clinic(project = paste0("TCGA-",ca.type), type = "clinical")
  MasterClinicalList$cancer.type <<- ca.type
```

```
 }
  else {
   currClinicalList <<-  GDCquery_clinic(project = paste0("TCGA-",ca.type), type = "clinical")
   currClinicalList$cancer.type <<- ca.type
   MasterClinicalList <<- rbind.fill(MasterClinicalList, currClinicalList)

  }

 return(1)

}

filelist$collectClinical <- lapply(filelist$cancer.type, GetClinicalData)
MasterList <- left_join(MasterClinicalList, MasterTranscriptList, by=c("submitter_id"="Patient"))

write.table(MasterList, file = "../rawdata/MasterMK2data.csv", quote=FALSE, sep="\t", col.names=TRUE, r
```

#### R Markdown

This is an R Markdown document. Markdown is a simple formatting syntax for authoring HTML, PDF, and MS Word documents. For more details on using R Markdown see http://rmarkdown.rstudio.com.

When you click the **Knit** button a document will be generated that includes both content as well as the output of any embedded R code chunks within the document. You can embed an R code chunk like this:
